# GSK3 inhibition reduces ECM production and prevents age-related macular degeneration-like pathology

**DOI:** 10.1101/2023.12.14.571757

**Authors:** Sophia M. DiCesare, Antonio J. Ortega, Gracen E. Collier, Steffi Daniel, Krista N. Thompson, Melissa K. McCoy, Bruce A. Posner, John D. Hulleman

## Abstract

Malattia Leventinese/Doyne Honeycomb Retinal Dystrophy (ML/DHRD) is an age-related macular degeneration (AMD)-like retinal dystrophy caused by an autosomal dominant R345W mutation in the secreted glycoprotein, fibulin-3 (F3). To identify new small molecules that reduce F3 production from retinal pigmented epithelium (RPE) cells, we knocked-in a luminescent peptide tag (HiBiT) into the endogenous F3 locus which enabled simple, sensitive, and high throughput detection of the protein. The GSK3 inhibitor, CHIR99021 (CHIR), significantly reduced F3 burden (expression, secretion, and intracellular levels) in immortalized RPE and non-RPE cells. Low-level, long-term CHIR treatment promoted remodeling of the RPE extracellular matrix (ECM), reducing sub-RPE deposit-associated proteins (e.g., amelotin, complement component 3, collagen IV, and fibronectin), while increasing RPE differentiation factors (e.g., tyrosinase, and pigment epithelium derived factor). In vivo, treatment of 8 mo R345W^+/+^ knockin mice with CHIR (25 mg/kg i.p., 1 mo) was well tolerated and significantly reduced R345W F3-associated AMD-like basal laminar deposit number and size, thereby preventing the main pathological feature in these mice. This is the first demonstration of small molecule-based prevention of AMD-like pathology in ML/DHRD mice and may herald a rejuvenation of interest in GSK3 inhibition for the treatment of neurodegenerative diseases, including, potentially AMD itself.

## INTRODUCTION

Fibulin-3 (F3, aka EFEMP1) is a secreted extracellular matrix (ECM) glycoprotein that is a member of the core matrisome. F3 is produced in a variety of ocular tissues including the corneal epithelium, trabecular meshwork ring, optic nerve, and neural retina/retinal pigment epithelium (RPE) (1). Upon secretion from the RPE, F3 is incorporated as a primary component of the RPE basal lamina (BL) meshwork, which acts as a sieve between the RPE and the underlying layers of Bruch’s membrane (BrM) (2). Appropriate maintenance of the RPE BL (3, 4), as well as permeability of BrM as a whole (5, 6), are important determinants for retinal health.

Whereas loss-of-function mutations or premature stop codons in *EFEMP1* (the gene that encodes for F3) have been associated with the development of connective tissue diseases resembling Marfan syndrome (7–9), increased copy number/expression of *EFEMP1* correlates with increased risk for age-related macular degeneration (AMD) (10, 11). Moreover, autosomal dominant mutations in *EFEMP1* have been linked to eye diseases including juvenile glaucoma (12, 13), primary open angle glaucoma (14), retinal degeneration (15), and a juvenile form of AMD called Malattia Leventinese/Doyne Honeycomb Retinal Dystrophy (ML/DHRD) (16, 17). Presumably autosomal dominant F3 mutations initiate their indicated associated disease by causing different degrees of protein misfolding (15, 18), increased F3 burden (i.e., altered extracellular/intracellular steady-state levels) (12, 19–21), changes to epithelial to mesenchymal transition (EMT) (22, 23), complement activation (4, 24, 25), or a combination thereof.

Yet, given F3’s broad ocular distribution, it may be surprising that both mice and humans lacking F3 show no clear defects in retinal structure or function (1). Moreover, removal of F3 appears to protect mice from environmentally-induced basal laminar deposits (BLamDs) (26), extracellular masses which are harbingers of RPE stress in aging and in AMD patients (2, 6). These results suggest that genetic or pharmacologic knockdown of F3 would be well-tolerated in the eye, and that its removal may in fact protect against both autosomal dominant F3 diseases (e.g., glaucoma or ML/DHRD) and age-related diseases influenced by F3, such as AMD.

However, to date, no small molecule therapeutics have been identified that can reduce the production of F3 from cells. Herein, we used CRISPR/Cas9 to genomically edit and tag endogenous F3 in cultured cells with an easily-detectable peptide tag followed by high throughput screening for novel reducing compounds. We discovered that glycogen synthase kinase 3 (GSK3) appears to be a key node in regulating F3 production and is also responsible for controlling the expression of additional ECM proteins, particularly those involved in sub-RPE deposits. Excitingly, treatment of 8 mo ML/DHRD R345W^+/+^ knockin mice with CHIR for 1 mo significantly reduced R345W F3-associated BLamD number and size, thereby preventing the main pathological feature in these mice. Overall, these data strongly support the use of GSK3 inhibitors for reducing sub-RPE pathology associated with misfolded F3 and possibly in idiopathic AMD.

## MATERIALS AND METHODS

### Cell culture

ARPE-19 cells (CRL-2302, ATCC) were cultured in DMEM/F12 (Corning) with 10% FBS (Omega Scientific) and 1% PSQ (ThermoFisher). Cells were kept at 37°C with 5% CO_2_, and passages were performed every 3-4 days, or when confluent. For experiments, ARPE-19 cells were plated at confluence (unless otherwise noted), typically at a concentration of ∼100,000 cells/cm^2^, which equates to ∼200,000 cells/mL. HEK-293T cells (Life Technologies) were cultured in DMEM High Glucose (4.5 g/L) supplemented with 10% FBS and 1% PSQ. Mouse NIH-3T3 fibroblasts (CRL-1658, ATCC) were maintained in DMEM high-glucose with 10% calf serum and 1% PSQ. Primary dermal fibroblasts (PCS-201-012, ATCC) were maintained in either low glucose DMEM with 10% FBS and 1% PSQ, or fibroblast growth media (Fibroblast Basal Medium [PCS-201-012, ATCC] supplemented with Low-Serum Fibroblast Growth Kit [PCS-201-041, ATCC]. Cells were periodically confirmed to be free of mycoplasma contamination using the Universal Mycoplasma Detection Kit (30-1012k, ATCC).

### HiBiT F3 cell line generation

Low passage ARPE-19 cells were genomically edited to introduce a 2xFLAG-VS-HiBiT sequence immediately after the signal sequence cleavage site (Ser16) of F3 (Figure 1A). This area in F3 was specifically chosen because an extreme N-terminal tag would be cleaved off after co-translational import into the endoplasmic reticulum (ER), and because we have anecdotally noticed that C-terminal appendages (of any size) compromise the secretion efficiency of F3, in line with recent observations that a stop codon mutation in *EFEMP1* results in intracellular retention and causes a juvenile form of glaucoma (12). Introduction of the 29 amino acid 2xFLAG-VS-HiBiT tag at this position was not predicted to affect the native F3 signal sequence cleavage (Supplemental Figure 1, A and B), and FLAG tags have been used in a similar manner to force signal sequence cleavage immediately prior to the tag (27). Briefly, a CRISPR/Cas9 ribonucleoprotein (RNP) was generated using Alt-R Sp. Cas9 Nuclease V3 (Integrated DNA Technologies, IDT) loaded with a crRNA/tracrRNA duplex (Supplemental Table 1). This RNP, combined with a single-stranded oligodeoxynucleotide (ssODN) repair template (Supplemental Table 1) were introduced into ARPE-19 cells using electroporation (1400 V, 20 ms, 2 pulses, Neon Transfection System, Life Technologies). After electroporation, cells were incubated in antibiotic-free media containing the DNA ligase IV inhibitor, SCR7 (1 μM, Sigma) for 48 h to promote homology-directed repair (HDR). Heterogenous cultures were expanded, assayed for secreted HiBiT via a luciferase assay and HiBiT blotting, and verified genomically for insertion of the 2xFLAG-VS-HiBiT sequence (primers listed in Supplemental Table 2). Full, uncropped gel images are located in Supplemental Figure 12.

**Figure 1:**
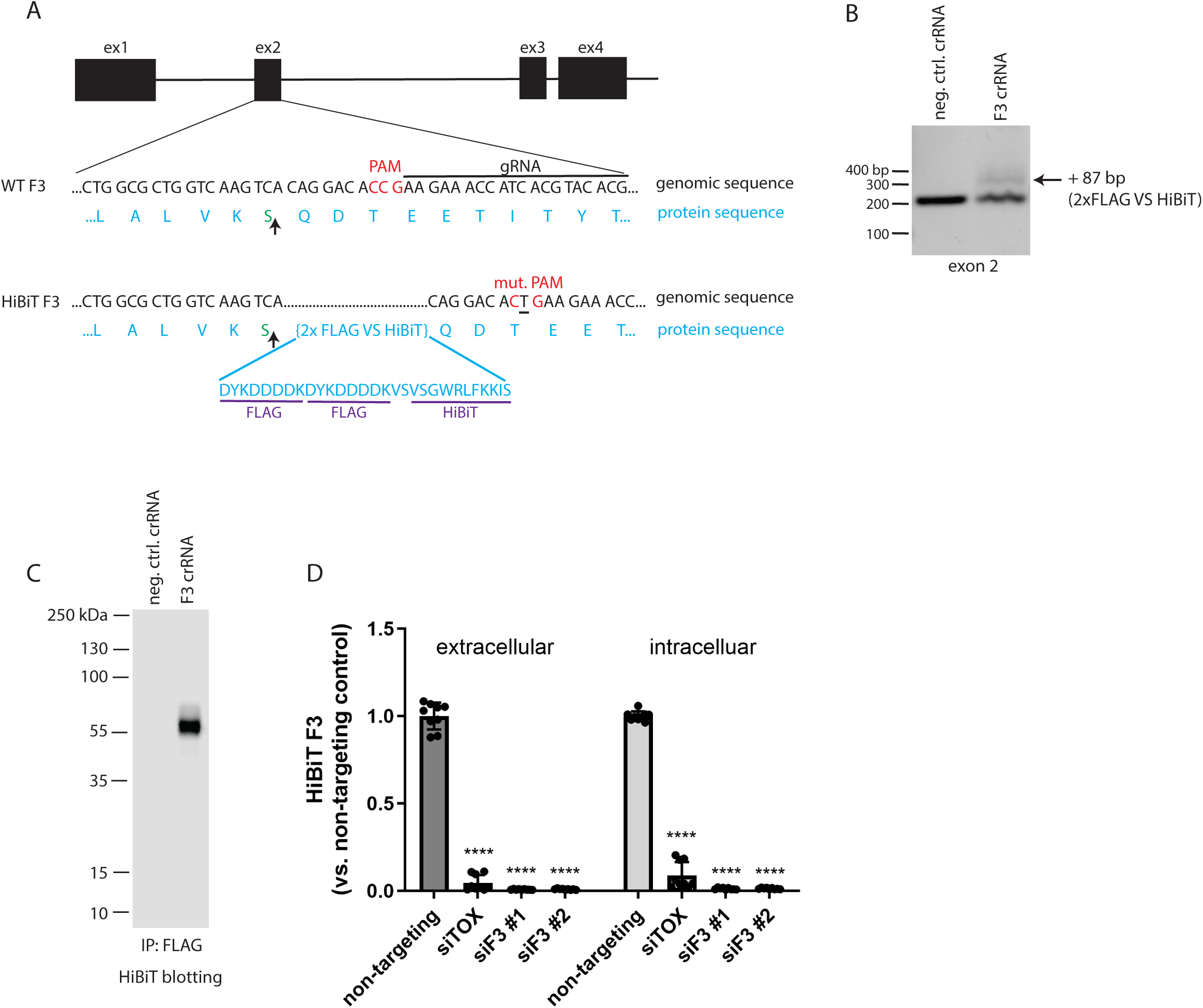
Design and validation of fibulin-3 (F3) HiBiT tagging in ARPE-19 cells using CRISPR. (A) Schematic of CRISPR editing of exon 2 of the F3 gene to knockin a 2xFLAG-VS-HiBiT sequence immediately proceeding the signal sequence cleavage site (upwards arrow). (B) Successful editing was verified by gDNA amplification of exon 2. An additional band corresponding to insertion of the 87 bp 2x FLAG VS HiBiT tag was identified with a calculated editing efficiency of ∼15%. (C) F3 HiBiT tagging results in a single extracellular protein species of correct molecular weight (∼55 kDa) as identified by immunoprecipitation (FLAG beads) followed by elution and HiBiT blotting. (D) siRNA verifies that >95% of the HiBiT signal can be attributed to F3 gene translation. n = 3 independent experiments performed in biological triplicates, **** p ≤ 0.0001, t-test vs. non-targeting siRNA.

### High-throughput screening (HTS)

Heterogenous ARPE-19 edited cells were used for all screening experiments due to their growth properties in comparison to single colony clones, which generally had much slower growth, making them subpar for large-scale HTS purposes. HiBiT F3 ARPE-19 cells were seeded at a density of ∼5,000 cells/well (in 30 μL with a MultiFlo, BioTek) in a white 384 well plate (781098, Greiner), and incubated at 37°C for 24 h. Media was exchanged using an EL406 microplate washer (BioTek). Compounds from the Prestwick Compound Library (1,200 compounds, Prestwick) and the National Institutes of Health clinical collection (446 compounds) were added using an Echo 655 (Beckman Coulter) at a final concentration of 5 μM and 0.1% DMSO. As a positive control for reduction of F3 secretion/production, brefeldin A (BFA, 50 µM) was used for the first column of each screening plate. Twenty-four hours after compound addition, plates were cooled to RT on the benchtop for 20 min followed by assaying for HiBiT signal in a whole well reading. LgBiT and lytic substrate (Promega) were added 1:100 and 1:50, respectively, into lytic buffer to generate a master mix. Fifteen microliters of the lytic/LgBiT/substrate master mix was dispensed into each plate using a MultiFlo, shaken for 5 min, and luminescence was detected on an Envision plate reader (PerkinElmer).

HiBiT assay performance was calculated by determining the Z factor (Z’) using the following equation (28):

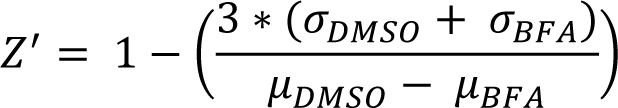

A counter screen was used to identify toxic compounds that would be predicted to yield a false positive result (i.e., reduction in F3 signal due to cell death). Cells were plated using the same methods as the initial screen. Cherry-picked hit compounds were added the following day at the standard screening concentration (5 µM) and an additional dose that was 0.5 log below the screening dose. The Cell Titer Glo 2.0 (Promega) viability assay was performed after 24 h of treatment. A whole-well analysis was completed by adding 10 µL of Cell Titer Glo diluted 1:1 with PBS + 0.1% TX-100 to the plate and shaking for 5 min. Luminescence was again read on the Envision plate reader (PerkinElmer).

In addition to the cell viability counter screen, media alone was treated with the hit compounds to identify any false reductions in luminescent signal caused by NanoBiT luciferase inhibition. Briefly, conditioned media was collected from ARPE-19 HiBiT F3 cells, spun, and plated into 384-well plates. Hit compounds were added in a dose-response format and incubated for 24 h. Lytic buffer/LgBiT/substrate master mix was added to the plates, shaken for 5 min, then read on an Envision plate reader. Wells that had significantly lower amounts of HiBiT signal were noted and the corresponding compounds were removed from subsequent experiments. Remaining hit compounds were verified in a confirmatory screen using fresh compound from a new source plate. Additional secondary metrics verified the effect of a hit compound in multiple assays and in different cell lines.

### Short interfering RNA (siRNA)

siRNA knockdown was used to verify the specificity of CRISPR HiBiT editing of F3 or fibulin-5 (F5). siRNAs (Silencer Select, Ambion, Supplemental Table 3) were introduced into cells containing HiBiT-tagged human F3, mouse F3, or human F5 by reverse transfection. For a 24 well plate, 2.14 μL of DharmaFECT4 (Horizon) was diluted into 250 µL OptiMEM (Thermo Fisher Scientific). siRNAs were added at a concentration of 100 nM. All samples were vortexed for 15 sec, then incubated at RT for 20 min. During this incubation period, ARPE-19 cells were trypsinized and normalized to a density of 466,000 cells/mL in full DMEM/F12 media. Two-hundred and fifty microliters of the siRNA/OptiMEM/DharmaFECT4 complex was added to a 24-well plate, after which an equivalent volume of cell suspension was added, yielding a final siRNA concentration of 50 nM. Plates were rocked to ensure cell distribution and to prevent clumping. After 24 h, the media was changed with full DMEM/F12. Seventy-two hours post knockdown, the media was changed again, and the HiBiT levels were determined the next day (96 h post knockdown) via an extracellular and intracellular HiBiT assay.

### Non-HTS HiBiT assay

Cells were plated at a high density of 200,000 cells/mL for all non-HTS HiBiT assay experiments. The following day, media containing the indicted compound was added and incubated for either 72 h or 1 week. For one-week treatment wells, media was replaced after 96 h with fresh media containing compound. After treatment, extracellular and intracellular assays were performed to determine levels of HiBiT-tagged F3 using either the Nano-Glo HiBiT Extracellular Detection System or the Nano-Glo HiBiT Lytic Detection System (Promega). For the extracellular HiBiT protocol, 25 µL of the conditioned media was reacted with an equal volume of HiBiT master mix (extracellular or lytic buffer, 1:100 LgBiT, and 1:50 extracellular or lytic substrate) in a black-bottomed 96-well plate for 5 min with shaking. For intracellular HiBiT detection, cells were washed in well with HBSS (Gibco), followed by addition of 50 µL of lytic HiBiT master mix (described above) and 5 min shaking. Forty microliters of the lysate was transferred to a black 96-well plate for a luminescent reading on a Synergy 2 (BioTek) or a GloMax (Promega).

### Western blotting

Cells for western blotting were rinsed with HBSS followed by lysis in buffer containing radioimmunoprecipitation buffer (RIPA, Santa Cruz), protease inhibitor (Pierce) and benzonase (Millipore Sigma), or the Nano-Go HiBiT Lytic Detection System. Lysis buffer was added to each well at a volume of 50 µL (for a 24 well plate, for example) followed by shaking for 5-15 min, and lysate was collected into Eppendorf tubes on ice. The samples were spun at 4°C for 10 min at 21,000 x g, and the soluble supernatant was collected. For cells lysed in RIPA buffer, a bicinchoninic acid assay (BCA, Pierce Thermo Scientific) was used to quantify protein levels and normalize them to 20 μg. For cells lysed in HiBiT Lytic buffer, samples were normalized by volume due to the presence of DTT, which interferes with the BCA assay. Samples were boiled in 1x reducing Laemmli buffer for 5 min, then loaded onto a 4-20% Tris-Gly SDS-PAGE gel and run at 140 V for 80 min. Proteins were transferred to 0.2 µm nitrocellulose membrane using a semidry apparatus (P0 protocol, iBlot2, Life Technologies). Total transferred protein was visualized with Ponceau S (Sigma-Aldrich) and incubated overnight in blocking buffer (Intercept Blocking Buffer, LI-COR). The following day, blots were incubated for 1 h at RT with primary antibody [GSK3α (sc-5264, Santa Cruz), GSK3β (9315S, Cell Signaling), or GAPDH (sc-47724, Santa Cruz)] diluted in 5% bovine serum albumin (BSA) in Tris-buffered saline (TBS) and 0.05% NaN_3_. Blots were rinsed in TBS with 0.05% Tween (TBS-T) and incubated for 40 min at RT in an appropriate anti-mouse or anti-rabbit secondary antibody (1:10,000-15,000, LI-COR) dissolved in 5% milk in TBS-T. Blots were again washed in TBS-T and imaged on an Odyssey CLx (LI-COR), followed by analysis using the corresponding ImageStudio Software (LI-COR). Full, uncropped blot images are located in Supplemental Figure 12.

### HiBiT blotting

Samples were lysed, prepared, and separated by SDS-PAGE as described above for western blotting. Normalized protein was reduced, denatured, and run through an SDS-PAGE gel, followed by transfer to a nitrocellulose membrane (P0 program, iBlot2). Once transferred, the membrane was incubated in TBS-T (10 min to 1 h, RT) to expose the HiBiT epitope. Blots were then incubated with blotting buffer containing 1:200 LgBiT for 1-2 h with rocking at RT (N2410, Promega). Next, NanoGlo substrate was added (1:500) and the blots were incubated for 5-10 min at RT. The resulting chemiluminescence signal was imaged on an Odyssey Fc (LI-COR). Full, uncropped blot images are located in Supplemental Figure 12.

### Lactate dehydrogenase (LDH) release assay

Cells were treated with CHIR for either 72 h or 1 week. Media was then collected and spun at 1,000 RPM for 5 min to remove potential floating cells. Fifty microliters (technical triplicates) of cleared media was mixed with an equal amount of LDH reaction buffer (LDH Cytotoxicity Assay Kit, Pierce) in a 96-well plate for 30 min at 37°C. A DMSO-treated media sample served as a control while cells treated with cell lysis buffer (provided in the LDH Cytotoxicity Kit) served as a positive control. Fifty microliters of stopping solution was added to each well and the absorbance at 490 nm was read on a GloMax (Promega).

### 7xTCF-eGFP mCherry (7TGC) and 7xTCF-firefly luciferase puromycin (7TFP) lentivirus production

VSV-g-pseudotyped replication incompetent lentivirus was generated as we have described previously (29, 30). Briefly, lentiviral plasmids (Addgene plasmids #24304 and #24308, kind gifts of Roel Nusse (31)) were co-transfected with psPAX2 and VSV-g plasmids (Addgene plasmids #12260 and #12259, gifts of Didier Trono) into low passage HEK-293T cells plated at 1×10^6^ cells/well of a poly-D-lysine-coated 6-well plate using Lipofectamine 3000 (Life Technologies). The next day, media was discarded and replaced with fresh media. Media collected 24 h and 48 h later was pooled and filtered through a 0.45 μm filter. Lenti-X GoStix (Takeda) were used to confirm the production of virus. To establish 7TGC and 7TFP cell lines, HiBiT F3 ARPE-19 cells were plated at 1×10^6^ cells/well of a 6 well plate and infected with lentivirus in full media containing polybrene for 24 h. 7TGC cells were not put under selective pressure whereas 7TFP cells were then selected with 1 μg/mL of puromycin for 1-2 weeks.

### Firefly luciferase (FLuc) assay

The HiBiT F3 7TFP ARPE-19 cells were treated with CHIR for 1 week (with a 72 h media change) and assayed for FLuc expression. Briefly, after a total of 1 week of treatment, media was aspirated and cells were washed with HBSS. Fifty microliters lysis buffer (Pierce Firefly Assay) was added to each well, then the plate was rocked for 15 min. To monitor FLuc activity, 50 µL of working solution containing luciferin and Firefly Glow Assay Buffer was added to a black 96-well plate. Ten microliters of the lysate was added to respective wells, then read on a Synergy2 plate reader.

### MMP2 zymography

HiBiT WT F3 or HiBiT R345W F3 ARPE-19 cells were plated at a density of ∼100,000 cells per well of a 12-well, or ∼30,000 cells per well of a 24-well polyester transwell plate (Corning), and media was changed to serum free media the following day and every 3-4 days thereafter. After 1 week growing on transwells, cells were treated with either DMSO (0.1%) or 1 μM CHIR in serum free media. After 2 weeks on transwells (1 week of treatment), an extracellular HiBiT assay and MMP2 zymography were performed. Briefly, for zymography, media from the apical and basal chamber was collected and combined with non-reducing SDS buffer followed by running on a 10% gelatin gel (Novex) for 90 minutes at 140 V. MMP2 was renatured in buffer (G-biosciences) for 30 min at RT. Gels were then changed to developing buffer (G-Biosciences) for an additional 30 min at RT. After changing to fresh developing buffer, gels were gently shook overnight at 37°C. The following day, the gels were stained in Coomassie R-250 for 1 h, then destained and imaged on an Odyssey Clx (LI-COR). Bands were quantified using Image Studio software (LI-COR). Full, uncropped gel images are located in Supplemental Figure 12.

### Secreted proteome visualization and mass spectrometry

HiBiT ARPE-19 cells were treated with CHIR for 72 h in serum-free DMEM/F12 media. To visualize separated total protein, conditioned media was concentrated (Amicon Ultra 0.5 mL centrifuge unit, 3,000 MWCO, Millipore Sigma) and run on a 4-20% Tris-Gly SDS-PAGE gel for 80 min at 140 V. Protein bands were imaged by silver staining (SilverQuest Silver Staining Kit, Invitrogen). In parallel, to prepare samples for mass spectrometry, an aliquot of the same concentrated media sample was run for 10 min at 140 V on a 4-20% Tris-Gly SDS-PAGE gel, stained with Coomassie Blue, and the single band corresponding to total secreted protein was excised and submitted for mass spectrometry. Briefly, samples were digested overnight with trypsin (Pierce) following reduction and alkylation with DTT and iodoacetamide (Sigma Aldrich). The samples then underwent solid-phase extraction cleanup with an Oasis HLB plate (Waters) and 2 μL of each sample was injected onto a QExactive HF mass spectrometer (Thermo Fisher Scientific) coupled to an Ultimate 3000 RSLC-Nano liquid chromatography system (Thermo Fisher Scientific). Samples were injected onto a 75 μm i.d., 15-cm long EasySpray column (Thermo) and eluted with a gradient from 0-28% buffer B over 90 min with a flow rate of 250 nL/min. Buffer A contained 2% (v/v) ACN and 0.1% formic acid in water, and buffer B contained 80% (v/v) ACN, 10% (v/v) trifluoroethanol, and 0.1% formic acid in water. The mass spectrometer operated in positive ion mode with a source voltage of 2.5 kV and an ion transfer tube temperature of 275°C. MS scans were acquired at 120,000 resolution in the Orbitrap and up to 20 MS/MS spectra were obtained for each full spectrum acquired using higher-energy collisional dissociation (HCD) for ions with charges 2-8. Dynamic exclusion was set for 20 s after an ion was selected for fragmentation. Raw MS data files were analyzed using Proteome Discoverer v. 2.4 SP1 (Thermo), with peptide identification performed using Sequest HT searching against the human protein database from UniProt (https://www.uniprot.org/). Fragment and precursor tolerances of 10 ppm and 0.6 Da were specified, and three missed cleavages were allowed. Carbamidomethylation of Cys was set as a fixed peptide modification, with oxidation of Met set as a peptide variable modification. The false-discovery rate (FDR) cutoff was 1% for all peptides.

### Sample preparation for RNAseq

HiBiT F3 ARPE-19 cells were plated at 200,000 cells/mL in DMEM/F12 media with 10% FBS and 1% PSQ. The following day, media containing DMSO (1:1000) or 1 μM CHIR was added to the cells. Cells were treated for 96 h before an additional media change. Seven days after beginning treatment, RNA from vehicle (DMSO)-treated and 1 μM CHIR-treated cells was extracted using an Aurum Total RNA Kit (BioRad) and stored at -80°C. Fifteen microliters of sample was provided to Novogene. A total amount of 1 μg RNA per sample was used as input material for the RNA sample preparations. Sequencing libraries were generated using NEBNext UltraTM RNA Library Prep Kit for Illumina (NEB) following manufacturer’s recommendations and index codes were added to attribute sequences to each sample. Briefly, mRNA was purified from total RNA using poly-T oligo-attached magnetic beads. Fragmentation was carried out using divalent cations under elevated temperature in NEBNext First Strand Synthesis Reaction Buffer (5X). First strand cDNA was synthesized using random hexamer primer and M-MuLV Reverse Transcriptase (RNase H). Second strand cDNA synthesis was subsequently performed using DNA Polymerase I and RNase H. Remaining overhangs were converted into blunt ends via exonuclease/polymerase activities. After adenylation of 3’ ends of DNA fragments, NEBNext Adaptor with hairpin loop structure were ligated to prepare for hybridization. In order to select cDNA fragments of preferentially 150∼200 bp in length, the library fragments were purified with AMPure XP system (Beckman Coulter). Then 3 μL USER Enzyme (NEB) was used with size-selected, adaptor ligated cDNA at 37 °C for 15 min followed by 5 min at 95 °C before PCR. Then PCR was performed with Phusion High-Fidelity DNA polymerase, Universal PCR primers and Index (X) Primer. At last, PCR products were purified (AMPure XP system) and library quality was assessed on the Agilent Bioanalyzer 2100 system.

### CHIR PK experiment

C57BL/6J (originally from Jackson Laboratories) were dosed via intraperitoneal injection with CHIR trihydrochloride (Tocris) dissolved in 2% DMSO with 5.7% Captisol (Ligand Pharmaceuticals) in 1x PBS at a concentration of 5 mg/mL. Mice were dosed at 25 mg/kg (average weight of 24.7 g). The dosing time points were 30, 180, 360, 960, and 1440 min, with a 0-min control group. Each timepoint included three mice, with at least one male and one female in each group. At select time points, isoflurane was used to anesthetize mice prior to tissue collection. Plasma, liver, and neural retina (both eyes pooled) were collected, and the samples were flash frozen in liquid nitrogen. All samples were then sent to mass spectroscopy and the amount of CHIR was measured in each tissue using LC-MS/MS (see below).

### LC-MS/MS analysis of CHIR

Retina tissue, liver tissue and plasma were analyzed for CHIR concentrations using an LC-MS/MS method. Retinas and livers were homogenized in PBS. Retina homogenates were made using BeadBug prefilled tubes with 3.0 mm Zirconium beads (Z763802, Sigma) and a BeadBug microtube homogenizer run for 1 min at 2800 rpm. For standards, blank commercial plasma (Bioreclamation) or untreated liver or brain tissue homogenate was spiked with varying concentrations of compound. Standards and samples were mixed with three-fold volume of 100% acetonitrile containing 0.133% formic acid and 33.3 ng internal standard (tolbutamide, Sigma), vortexed, and then spun 5 min at 16,100 x g. Supernatant was removed and spun again and the resulting second supernatant was put into an HPLC 96-well plate and analyzed by LC-MS/MS using a Sciex Triple Quad™ 4500 mass spectrometer coupled to a Shimadzu Prominence LC. CHIR was detected with the mass spectrometer in MRM (multiple reaction monitoring) mode by following the precursor to fragment ion transition 465 → 146.1. An Agilent Zorbax XDB-C18 column (50 x 4.6mm, 5 micron packing) was used for chromatography with the following conditions: Buffer A: dH20 + 0.1% formic acid, Buffer B: MeOH + 0.1% formic acid, 1.5mL/min flow rate, 0-1.5 min 3%B, 1.5-2.0 min gradient to 100%B, 2.0-3.5 min 100%B, 3.5-3.6 min gradient to 3%B, 3.6-4.5 3%B. Tolbutamide (transition 271.2 → 91.2) was used as an internal standard. Back-calculation of standard curve and quality control samples were accurate to within 15% for 85-100% of these samples at concentrations ranging from 0.5 ng/ml to 10000 ng/ml.

### In vivo treatment of CHIR

As the PK experiment demonstrated successful penetration of CHIR into the retina, we used the same CHIR formulation for in vivo treatment of R345W^+/+^ knockin mice. Eight-month-old R345W^+/+^ mice (25) were divided into a treatment group and control group (n = 4 mice/group, 2 male and 2 female mice). CHIR (25 mg/kg) or PBS (vehicle) was delivered via i.p. injections given every 24 h during the 5-day work week for one month.

### Electroretinogram (ERG)

After the one-month of vehicle or CHIR treatment, scotopic ERG was performed. Mice were dark-adapted the night before ERG. A-waves and b-waves were monitored in a dark-adapted intensity series using 6 different light intensities (0.003-30 cd.s/m^2^, Celeris, Diagnosys). Each mouse was weighed and given a ketamine/xylazine solution diluted 1:1 in biostatic water (120 mg/kg, 16 mg/kg respectively, final concentration). Once mice were unresponsive to a toe pinch, tropicamide was administered to dilate the pupil. A probe was placed into the skin between the eyes, and a ground wire was placed into the skin near the base of the tail. GenTeal Severe Dry Eye gel (Alcon) was placed onto each eye, and electrodes were aligned to face the retina. Interference was reduced to below 8.0 kΩ before starting the scan. This process was repeated for each mouse, and the a- and b-wave amplitudes were provided by the Espion software (Diagnosys). Outlier data points were identified using GraphPad Prism.

### Sample processing for Transmission Electron Microscopy (TEM) and histology

Nine-month-old vehicle or CHIR-treated mice were deeply anesthetized using an i.p. ketamine/xylazine injection. Once unresponsive to foot pinch, they were immediately transcardially perfused with 4% paraformaldehyde. An incineration mark was made temporally on the cornea prior to enucleation using a fine tip cautery to provide a reference for subsequent eye orientation. One eye was processed for histology using a freeze substitution protocol (32, 33). Briefly, the eye was enucleated and held in cold isopentane kept on dry ice for one min. The flash frozen eye was then placed in methoxy acetic acid and frozen at -80°C. Samples were slowly warmed to RT for histology submission over a few days. Briefly, after 48 h at -80°C, samples were moved to -20°C for 48 h, 4°C for 24 h, and finally to RT for at least 1 h prior to submission. The whole globe was transferred to an Eppendorf tube filled with 1.5 mL of 100% ethanol and submitted to the UT Southwestern Histo Pathology Core. Sections were taken through the optic nerve and images were recorded on a NanoZoomer S60 microscope (Hamamatsu).

For the TEM ocular sample, the contralateral eye was placed in half strength Karnovsky’s fixative (2% formaldehyde + 2.5% glutaraldehyde, in 0.1 M sodium cacodylate buffer, pH 7.4) on a rocker at RT. After 4 h of rocking in Karnovsky’s fixative, incisions were made at the incineration mark and along the ora serrata removing the lens and majority of the cornea, while leaving a corneal flap over the ventral temporal quadrant of the eye denoting the underlying retinal region of interest. The eye cups were placed in fresh half strength Karnovsky’s fixative overnight at 4°C. Eye cups were prepared for TEM by then rinsing three times in 0.1M sodium cacodylate buffer followed by 1% OsO_4_ in buffer (2 parts 0.1M Na cacodylate buffer pH 7.2, 1 part 4% osmium, 1 part 6% potassium ferrocyanide) for 2 h, washed 3 times in 0.1M sodium cacodylate buffer (pH 7.2) for 30 min. The sample was then rinsed twice with nanopure water, placed in 2.5% Uranyl Acetate in deionized water for 20 min followed by 25% ethanol for 30 min, 50% ethanol for 30 min, 75% ethanol for 30 min, 95% ethanol for 45 min, then two serial 100% ethanol incubations for 45 min each and kept overnight at 4°C.

The samples were then resin infiltrated by first replacing the ethanol by performing two 100% propylene oxide washes for 20 min, then 2 parts 100% propylene oxide per 1 part Eponate 12 resin (Ted Pella) for an 1 h, 1 part 100% propylene oxide per 2 parts Eponate 12 resin for 1 h, then left overnight in 100% Eponate 12 at RT. The next day the samples were placed in fresh Eponate 12 for 2 h then embedded with fresh Eponate 12 resin in a mold and placed in a 60°C oven overnight. Sections were cut using the corneal flap to orient the sample to section the underlying retinal region of interest. Mounted grids were stained with uranyl acetate and lead citrate contrast. The grids were imaged using a 1400+ TEM (Jeol). TIFF file images were acquired using the interfaced BIOSPRINT 16M-ActiveVu mid mount CCD camera (Advanced Microscopy Technologies).

TEM images were analyzed with ImageJ software. Brightness and contrast were adjusted for best visualization of Bruch’s membrane (BrM). Images were excluded from analysis if the membrane and/or deposits could not be reasonably delineated. The images were split into eight subfields of view using the grid function. A masked observer determined whether a deposit was present in each subfield of view with binary classification (1 = deposit, 0 = no deposit). Once the images were analyzed, a chi-square test was performed comparing treated and untreated mice, with p < 0.05 as the cutoff for significant values.

BrM and the deposit(s) were traced using the ImageJ tracing tool for some of the subfields with deposits. Some subfields of view did not have adequate resolution for this analysis and were not included. This exclusion resulted in 22 subfields of treated mice images that could be traced and analyzed. For comparison, 22 subfields from untreated mice images with deposits were randomly selected to be traced and analyzed. Prior to tracing, pixels were converted to µm. The areas of the traced deposits and membranes were reported in µm^2^.

## RESULTS

### Genome editing with HiBiT produces a sensitive method for following F3 production

As a secreted, extracellular matrix glycoprotein, endogenous F3 is typically difficult to detect in cultured cells (9) and in vivo (34). The challenge of monitoring F3 is compounded by its low levels, potential incorporation into the ECM, its monomeric molecular weight (55 kDa, the approximate size of BSA), and a dearth of F3 knockout-validated antibodies. We therefore designed a method to label endogenous F3 with an 11 amino acid HiBiT tag (35) (Figure 1A), which we theorized would facilitate quick and easy detection of endogenous F3 after complementation of HiBiT-tagged F3 with LgBiT, producing a bright and stable NanoBiT luminescent signal (35) proportional to F3 abundance. Using Sp. Cas9 ribonucleoprotein (RNP), we introduced a 2xFLAG-VS-HiBiT tag immediately after the F3 signal sequence through homology-directed repair (HDR, Figure 1A) in human adult retinal pigmented epithelial (RPE) cells (ARPE-19). This insertion was predicted to have no effect on F3 signal sequence processing (Supplemental Figure 1, A and B), and avoided potential disruptions to F3 secretion by appending additional amino acids to the C-terminus of F3 (vis-à-vis select glaucoma mutations (12)). Insertion of the HiBiT tag was verified by genomic DNA analysis, resulting in an amplicon with 87 additional base pairs (Figure 1B, ∼15% editing efficiency). Validation of the specificity of the edit was assessed by using a scrambled negative control crRNA/tracrRNA duplex and performing immunoprecipitation of the 2xFLAG tag followed by HiBiT blotting, demonstrating a single species of ∼55 kDa (Figure 1C), and short interfering RNA (siRNA) knockdown of F3 (siF3 #1, siF3 #2) vs. control siRNAs (non-targeting, Figure 1D). These observations suggest that our HiBiT editing approach is specific for F3 with little or no detectable off target effects.

### Miniaturization of the HiBiT F3 assay enables small molecule high-throughput screening (HTS)

Increased F3 production has been associated with cancers such as gliomas (36) and prevalent vision disorders such as AMD (10, 11). Additionally, rare autosomal dominant, presumably gain-of-function, mutations in F3 have been associated with diseases such as juvenile open angle glaucoma (12, 13) and the early onset AMD-like disease, ML/DHRD (17). Accordingly, identifying genetic or small molecule therapeutics designed to reduce F3 production could be therapeutically useful in these particular diseases. To facilitate this goal, we miniaturized the HiBiT assay into a 384-well format and performed HTS. An example mock plate (Supplemental Figure 2A) demonstrated consistent values in DMSO-treated wells (0.1%) with ∼7% coefficient of variation (C.V.) for a whole-well analysis (HiBiT signal originating from both media and cells). We next screened HiBiT F3 ARPE-19 cells against a Prestwick Chemical Library (Alsace, France) and a National Institutes of Health (NIH) clinical collection. While we acknowledge that ARPE-19 cells do not accurately represent ‘true’ RPE cells as would be found in a human (37), we rationalized that they are appropriate to use from a simple cell biology perspective; they are a cell line that produces endogenous F3, adhere well to HTS plates (to increase consistency), and can be used at scale in HTS applications. Example data from two HTS plates (Supplemental Figure 2, B and C) demonstrate the consistency of the assay and highlight potential reducers and enhancers of F3 production in RPE cells. Average Z’ score across all compound plates was excellent at 0.59 ± 0.06. One-hundred and twenty-seven F3 reducing compounds (Z-score < -3) were identified in the primary screen (hit percentage = 127/1646, 7.9% hit percentage). From these hits, 50 molecules were selected for confirmatory and counter screening. Compounds that affected HiBiT/LgBiT complementation or interfered with substrate binding to NanoBiT were identified in a counter screen using only HiBiT F3-containing media. Moreover, compounds that caused a >10% reduction in ATP levels (determined by a Cell Titer Glo 2.0 assay, Promega, Supplemental Table 4) over 48 h at both 1.66 μM and 5 μM were excluded. For example, MG-132 was toxic at 5 μM, but not at 1.66 μM, and was therefore retained for subsequent dose response assays. This triaging process yielded 8 hits (Supplemental Table 4), 7 of which were verified by dose-response (Supplemental Figure 3). Within this set of verified hits, two compounds, AZD2858 and CHIR98014, were both identified as GSK3 inhibitors (38, 39). These compounds demonstrated dose-responsiveness (Supplemental Figure 3) and had favorable cytotoxicity profiles (Supplemental Table 4). Based on this enrichment within our dataset, as well as the extensive use of GSK3 inhibitors for diverse diseases, including neurodegeneration (40, 41), we focused our efforts on characterizing inhibition of the GSK3 pathway as a unique way to regulate F3 production in cells.

In confirmatory assays, we validated the importance of the GSK3 pathway in regulating F3 production from RPE cells by expanding the diversity of GSK3 inhibitors tested, this time deconvoluting the whole well assay into secreted HiBiT F3 and intracellular HiBiT F3. Seventy-two-hour treatment with GSK3 inhibitors 6-bromoindirubin-3-oxime (BIO), CHIR98014, CHIR99021 (CHIR), and lithium chloride (LiCl) all significantly reduced HiBiT F3 secretion and intracellular levels (Figure 2, A and B) in a dose-responsive manner. Of these compounds, CHIR yielded the most consistent and effective results, reducing HiBiT F3 secretion up to 76% (10 μM, Figure 2A) and reducing intracellular HiBiT F3 up to 87% (10 μM, Figure 2B). Therefore, we prioritized subsequent testing of CHIR in blotting and quantitative PCR (qPCR) secondary assays. HiBiT blotting of conditioned media from RPE cells treated with CHIR (Figure 2C) confirmed the reduction in F3 secretion observed by HiBiT assay (Figure 2A). Moreover, it appears that CHIR reduces F3 secretion and intracellular levels by decreasing F3 transcription (23 ± 2% of vehicle-treated levels, Figure 2D). Moreover, lactate dehydrogenase (LDH) release assays confirmed that CHIR was non-toxic during these experiments (Figure 2E).

**Figure 2:**
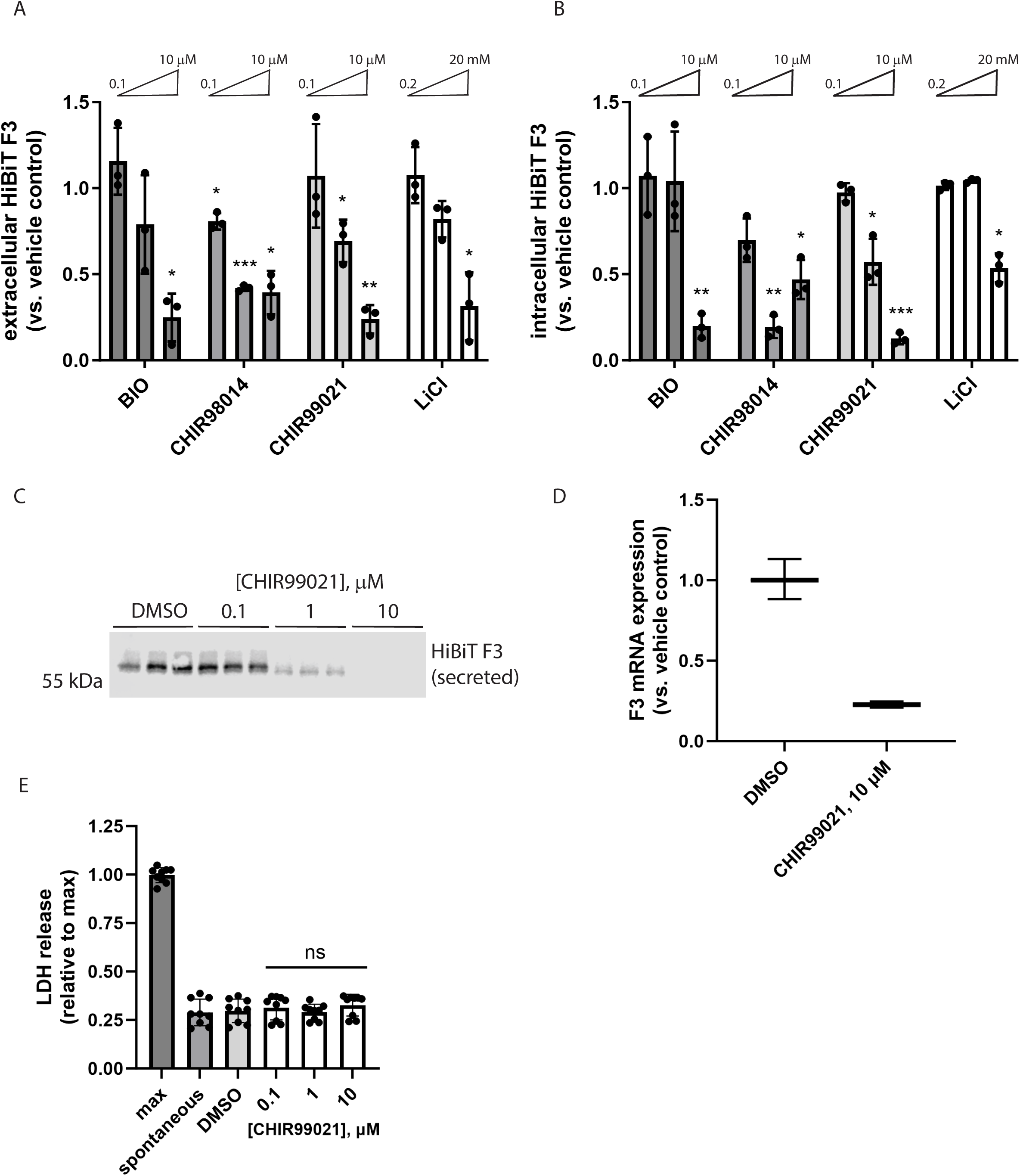
Structurally diverse glycogen synthase kinase 3 (GSK3) inhibitors reduce F3 transcripts and extracellular/intracellular levels without causing toxicity. (A, B) A series of chemically unrelated GSK3 inhibitors significantly reduce (A) extracellular and (B) intracellular HiBiT F3 levels after 72 h of treatment. n = 3 independent experiments, mean of each plotted here, * p ≤ 0.05, ** p ≤ 0.01, *** p ≤ 0.001, one sample t-test vs. hypothetical unchanged value of 1 (vehicle-treated). (C) CHIR99021-dependent HiBiT assay results in (A) were confirmed at the protein level by HiBiT blotting. Representative image of n ≥ 3 independent experiments. (D) Seventy-two hour CHIR99021 treatment reduces F3 mRNA expression. Representative data of n = 3 independent experiments, average ± S.D. of technical triplicates. (E) Treatment with CHIR does not elevate release of cytosolic lactate dehydrogenase (LDH). n = 3 independent experiments performed in biological triplicate, ns = not significant, t-test vs. vehicle-treated levels.

### siRNA knockdown of both GSK3α and GSK3β synergistically reduces F3 production

GSK3 has two structurally similar and potentially functionally redundant isoforms (42), GSK3α and GSK3β, both of which are produced in ARPE-19 cells (Figure 3, A and B), and both of which are targeted by many GSK3 inhibitors, such as CHIR, when it is used at elevated concentrations. To determine whether one GSK3 isoform was primarily responsible for reducing F3 production, we knocked down GSK3α and GSK3β individually and in combination by siRNA. Individual knockdown of GSK3α or GSK3β resulted in significant and nearly identical effects on HiBiT F3 secretion (∼30% reduction, Figure 3A) and intracellular levels (∼40% reduction, Figure 3A). Combination of GSK3α and GSK3β siRNAs further reduced HiBiT F3 levels in both the conditioned media (∼55% reduction, Figure 3A) and cell lysates (∼77% reduction, Figure 3A). GSK3α/β knockdown and HiBiT assay results were verified by western blotting (Figure 3B), demonstrating effective mRNA knockdown and providing an orthogonal confirmatory measurement of HiBiT F3 reduction. These results are a genetic confirmation that the CHIR-mediated effects on F3 are most likely occurring primarily through GSK3 inhibition.

**Figure 3:**
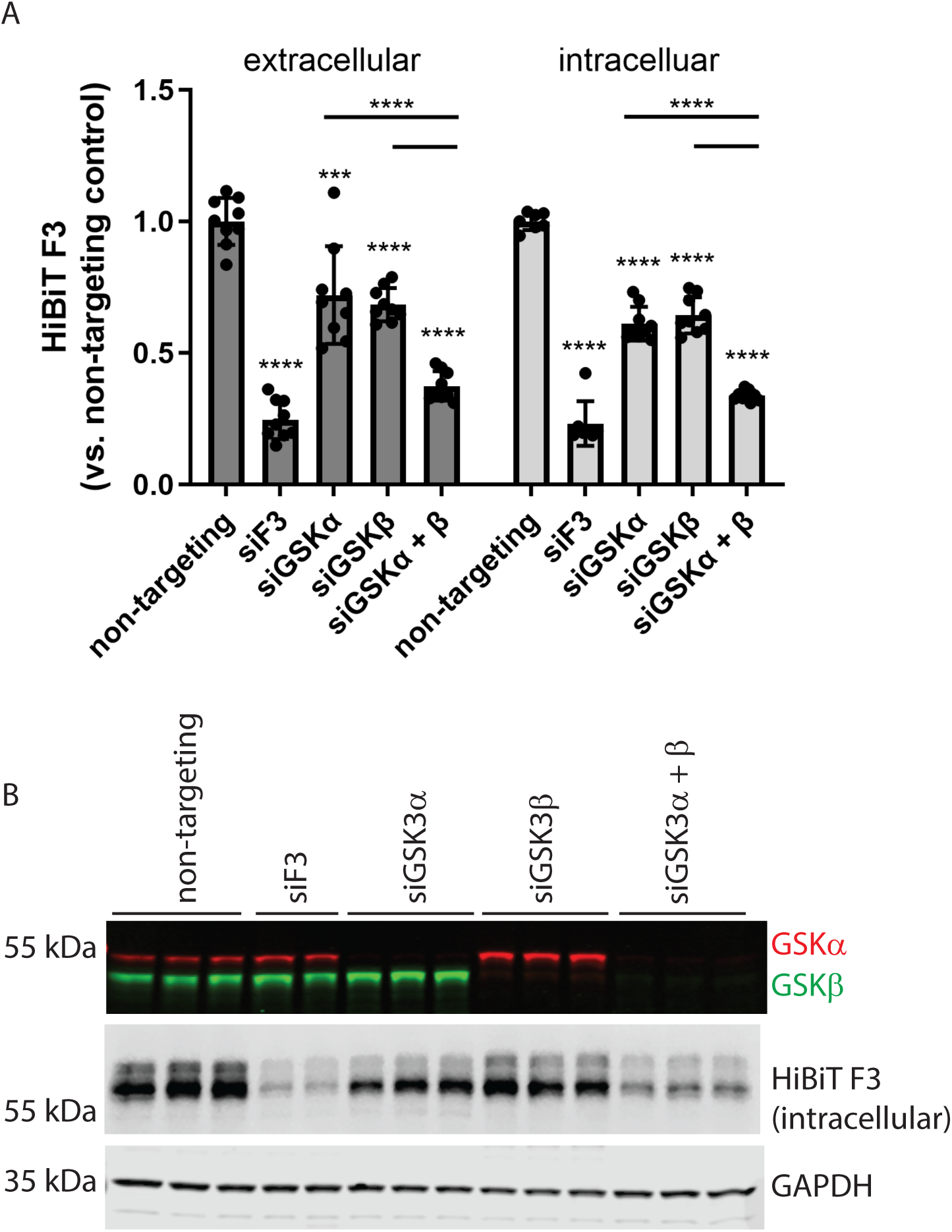
Genetic knockdown of GSK3 isoforms also reduces HiBiT F3 levels, paralleling pharmacologic GSK3 inhibitor effects. (A) Knockdown (96 h) of either GSK3α or GSK3β significantly lowers F3 production in ARPE-19 cells and the effects of knocking down both α and β isoforms are additive. n = 3 independent experiments performed in biological triplicate, *** p ≤ 0.001, **** p ≤ 0.0001, t-test vs. either vehicle-treated levels or single isoform knockdowns as indicated. (B) Protein-level knockdown effects were verified by western and HiBiT blotting. Representative images of n = 3 independent experiments.

### Testing the effects of CHIR in additional HiBiT-edited cell lines

To determine whether the effects of GSK3 inhibition on F3 production were specific to ARPE-19 cells, or represented a broader universal phenomenon conserved across cell lines, we genomically engineered two additional cell lines to express HiBiT F3: primary dermal fibroblasts and NIH-3T3 fibroblasts. Similar to ARPE-19 cells, CHIR significantly reduced both secreted and intracellular HiBiT F3 in human dermal fibroblasts in a dose-responsive manner (Supplemental Figure 4). Reassuringly, CHIR reduced F3 production in NIH-3T3 mouse cells also (Supplemental Figure 5, A and B, and Supplemental Figure 6, A-C), demonstrating a cross-cellular and cross-species effects of CHIR.

Since F3 belongs to a family of similarly structured fibulin proteins (43), we subsequently tested if CHIR could also regulate other fibulin proteins, such as the highly homologous, short fibulin, fibulin-5 (F5 or *FBLN5*), or if its effect was more specific to F3. Using an identical HiBiT editing strategy (Supplemental Figure 7, A and B, Supplemental Figure 8, A-D), we introduced a 2xFLAG HiBiT sequence onto the F5 protein in ARPE-19 cells and validated the specificity of editing using RNAi (Supplemental Figure 8B). Treatment of HiBiT F5 expressing cells with CHIR for 72 h resulted in dose-responsive reduction of secreted F5 across all concentrations used (Supplemental Figure 7C). An extended, week-long treatment with CHIR at the same doses also significantly reduced F5 secretion, but only at 1 and 10 μM (Supplemental Figure 8D). However, high concentrations of certain GSK3 inhibitors can lead to a ‘rebound’ effect, instead elevating F5 (Supplemental Figure 8D) or F3 levels (cf. Figure 2B, CHIR98014), possibly promoting intracellular retention or overriding the transcriptional reduction effect. Overall, these results suggest that the effects of CHIR on F3 appear to be cell line and species independent, and that this compound may be acting more broadly on secreted or ECM proteins (e.g., fibulins) than initially thought.

### Low level, long-term CHIR reduces HiBiT F3 secretion without triggering canonical Wnt signaling as indicated by T-cell factor (TCF) activation

Regulation of GSK3 kinase activity is an important determinant in epithelial to mesenchymal transition (EMT) decisions in multiple cell and in vivo contexts (44, 45). GSK3, in combination with casein kinase I (CKI), phosphorylates β-catenin and promotes its eventual degradation through the β-catenin destruction complex (46, 47). Thus, inhibition of GSK3 favors nuclear accumulation of β-catenin, which subsequently complexes with the transcriptional regulators lymphoid enhancing factor-1 and TCF, upregulating Wnt genes typically associated with increased EMT. Furthermore, complete genetic loss of *Gsk3*, and by extension, complete GSK3 inhibition, in mouse progenitor cells leads to microphthalmia and overt ocular morphological defects (42). Moreover, EMT triggered by sustained high levels of GSK3 inhibition of RPE cells may play an important role in the pathology of retinal diseases (22, 48). Thus, we sought to determine whether low level, longer term CHIR treatment could still reduce F3 production, but not trigger potential EMT through Wnt pathway activation.

HiBiT F3-expressing RPE cells were transduced with lentivirus encoding for constitutively expressed mCherry and a green fluorescent protein (GFP) driven by seven repeats of the TCF promoter (7TGC) or a puromycin-selectable lentivirus encoding for firefly luciferase (FLuc) driven by the same seven TCF repeats (7TFP) (31). Treatment of the 7TGC HiBiT F3 cells with CHIR (0.675 – 10 μM) for 72 h resulted in a predictable reduction in HiBiT F3 secretion (Figure 4A). Yet only concentrations greater than or equal to 2.5 μM resulted in any detectable GFP signal indicative of Wnt pathway activation (Figure 4B). Based on these findings, we next asked whether low level CHIR treatment (≤1 µM) for longer periods of time would further reduce F3 production without triggering TCF activation. One week treatment with CHIR significantly reduced F3 secretion at 1 µM (by 49.9 ± 7.0%, Figure 4C) without triggering TCF activation or toxicity (Figure 4, D and E). These data suggest that CHIR can be used at low concentrations to reduce F3 production while not detectably activating potentially detrimental Wnt/EMT-related signaling.

**Figure 4:**
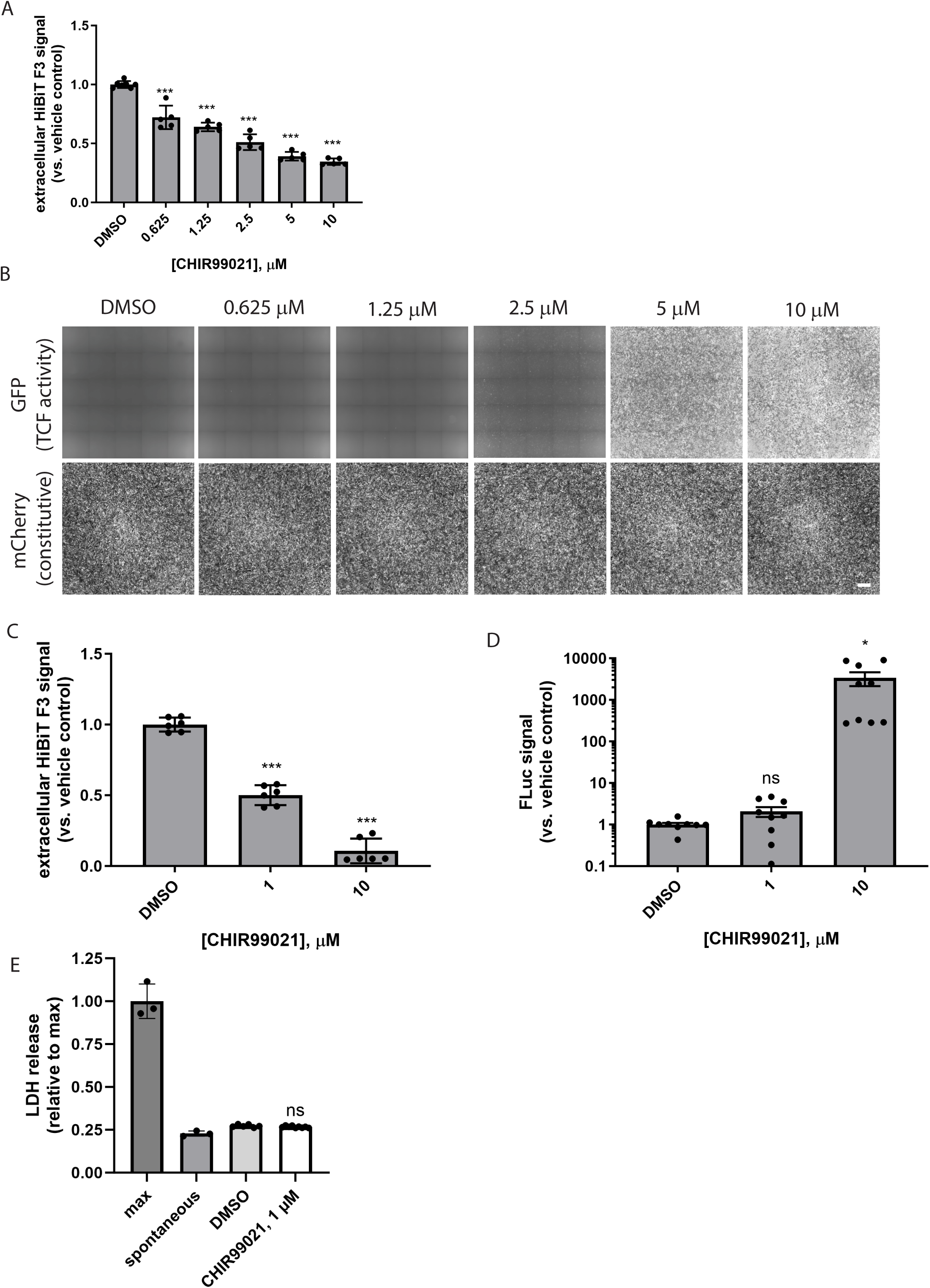
Low level, prolonged CHIR99021 treatment reduces F3 production while avoiding triggering TCF4-dependent Wnt activation. (A) Seventy-two-hour CHIR99021 treatment dose-dependently reduces HiBiT F3 extracellular levels without necessarily triggering TCF4-dependent green fluorescent protein (GFP) expression (B). n = 3 independent experiments performed in at least biological triplicate, with representative data presented. *** p ≤ 0.001, t-test vs. vehicle treatment. Scale bar ∼ 200 μm (C) One-week treatment with CHIR99021 (1 μM) significantly lowers HiBiT F3 extracellular levels without activating TCF4-dependent firefly luciferase (FLuc). n = 3 independent experiments performed in biological replicates. * p ≤ 0.05, *** p ≤ 0.001, t-test vs. vehicle treatment. (E) Prolonged low level CHIR99021 does not increase cell death as indicated by LDH release. n = 3 independent experiments performed in biological replicates. ns = not significant, t-test vs. vehicle treatment.

### Proteomics and transcriptomics confirm that low level GSK3 inhibition reduces production of collagen, basement membrane, and ECM components

Using our optimized CHIR dosage and treatment window, we next determined what additional secreted proteins were altered by low-level GSK3 inhibition. SDS-PAGE-separated concentrated media from CHIR-treated HiBiT F3 cells demonstrated a clear increase in total protein secretion at 10 μM, but relatively little detectable change in banding pattern or intensity at 0.1 or 1 μM (Supplemental Figure 9). Proteomics unbiasedly determined that a total of 40 proteins were decreased by at least 50% (≥ 2-fold) compared to the vehicle control after 1 week of CHIR treatment (1 μM, Table 1, top 50 reduced proteins shown). F3 secretion was decreased in the proteomic data set by 37%, on par with our previous HiBiT observations (Figure 4C) but did not meet the 50% reduction inclusion criteria for this data set (Table 1). Nonetheless, notable proteins identified in this downregulated list include i) collagens [collagen alpha-1(V), collagen alpha-2(IV), collagen alpha-1(IV), collagen alpha-3(IV)], ii) a major ECM protease [matrix metalloproteinase 2 (MMP2)], iii) transforming growth factor beta (TGFβ)-related proteins [plasminogen activator inhibitor 1 (PAI-1 or SERPINE1), latent-TGFβ-binding protein 2 (LTBP2), and TGFβ1], iv) inflammatory/complement markers [high mobility group protein B1 (HMGB1), HMGB2, and complement component 3 (C3)], and v) other members of the fibulin protein family [fibulin-1 (FBLN1) and fibulin-2 (FBLN2)]. Gene ontology (GO) analysis of the top 50 decreased proteins identified significant enrichment of proteins involved in collagen trimer formation, basement membrane, and extracellular matrix (Table 2). Conversely, more proteins were increased (154) by ≥ 2-fold (Supplemental Table 5, top 50 increased proteins shown) than were decreased. GO analysis of the top 50 of these proteins identified a range of cellular components including granule formation, chaperone complex, and ribosomal subunits (Supplemental Table 6).

**Table 1:**
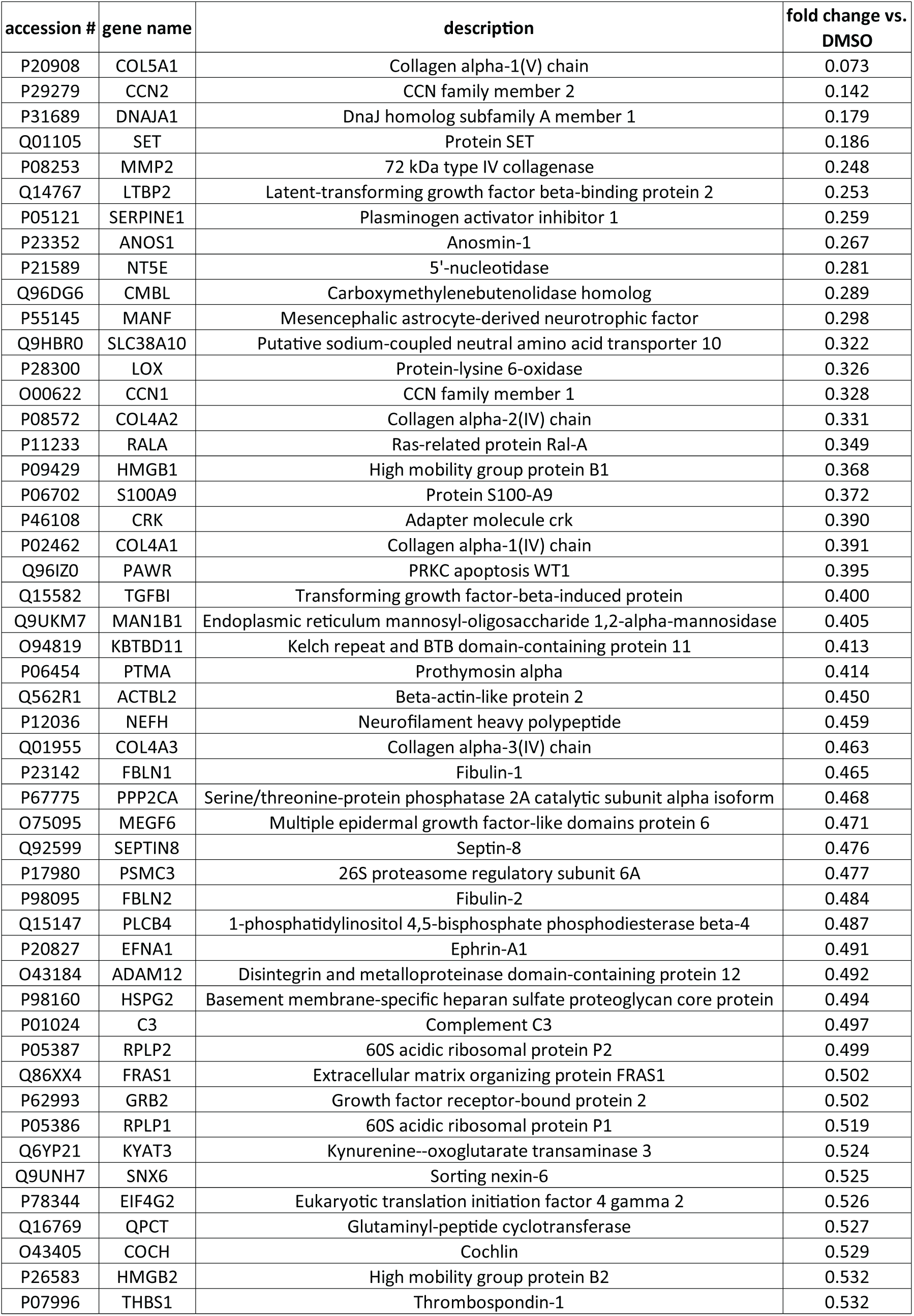
Top 50 proteins reduced (40 by ≥ 2-fold) after one-week CHIR99021 treatment of ARPE-19 cells.

**Table 2:** GO analysis of significantly reduced proteins.

| GO cellular component | # of genes in H. sapiens reference list | # of genes in dataset | expected frequency | fold enrichment | raw p value | FDR |
| --- | --- | --- | --- | --- | --- | --- |
| collagen type IV trimer (GO:0005587) | 7 | 3 | 0.01 | > 100 | 8.03E-07 | 1.49E-04 |
| basement membrane collagen trimer (GO:0098651) | 9 | 3 | 0.02 | > 100 | 1.47E-06 | 2.14E-04 |
| collagen network (GO:0098645) | 9 | 3 | 0.02 | > 100 | 1.47E-06 | 2.00E-04 |
| network-forming collagen trimer (GO:0098642) | 9 | 3 | 0.02 | > 100 | 1.47E-06 | 1.87E-04 |
| complex of collagen trimers (GO:0098644) | 22 | 4 | 0.04 | 93.59 | 1.76E-07 | 3.59E-05 |
| basement membrane (GO:0005604) | 101 | 7 | 0.2 | 35.67 | 1.43E-09 | 4.88E-07 |
| collagen trimer (GO:0005581) | 88 | 5 | 0.17 | 29.25 | 9.70E-07 | 1.52E-04 |
| collagen-containing extracellular matrix (GO:0062023) | 434 | 15 | 0.84 | 17.79 | 2.26E-15 | 4.62E-12 |
| extracellular matrix (GO:0031012) | 575 | 16 | 1.12 | 14.32 | 5.60E-15 | 3.82E-12 |
| external encapsulating structure (GO:0030312) | 576 | 16 | 1.12 | 14.3 | 5.75E-15 | 2.94E-12 |

In parallel experiments, RNA from HiBiT F3 cells treated identically with vehicle or CHIR for 1 week was sent for RNAseq analysis (Novogene). One-hundred and sixty-five genes were identified as being significantly downregulated by at least 50% (≥ 2-fold, Figure 5A, Supplemental Table 7), while 81 genes were identified as significantly upregulated by ≥ 2-fold (Figure 5A, Supplemental Table 8). Many of the proteins identified in the proteomic data set (e.g., COL4, F3 (EFEMP1), C3, etc.) were confirmed by RNAseq analysis (Supplemental Tables 7 and 8). Moreover, pathway analysis of genes that were significantly changed in either direction in the RNAseq dataset further confirm that low level GSK3 inhibition with CHIR results in substantial changes to the production and organization of the ECM as well as the basement membrane composition (Figure 5B). Additionally, genes of interest pertaining to sub-RPE deposits including amelotin (AMTN (49)) and fibronectin (FN (50)), ECM crosslinking enzymes such as lysyl oxidase (LOX (51)), and pro-EMT regulators such as WNT3 (52) and WNT5A (53) (Figure 5A) were all downregulated. Interestingly, in the upregulated gene set, two important genes for pigment formation in the RPE (54), tyrosinase (TYR) and TYR-related protein 1 (TYRP1), were significantly upregulated by CHIR treatment (Figure 5A). An additional, potentially beneficial neuroprotective protein produced in the RPE (55), pigment epithelium-derived factor (PEDF, a.k.a. SERPINF1), was also significantly upregulated in both the proteomics and RNAseq datasets by CHIR treatment (Supplemental Table 5, Figure 5A). Similar observations regarding increases in TYR and PEDF have been made after supplementing CHIR during a human embryonic stem cell (hESC)-to-RPE differentiation protocol (56).

**Figure 5:**
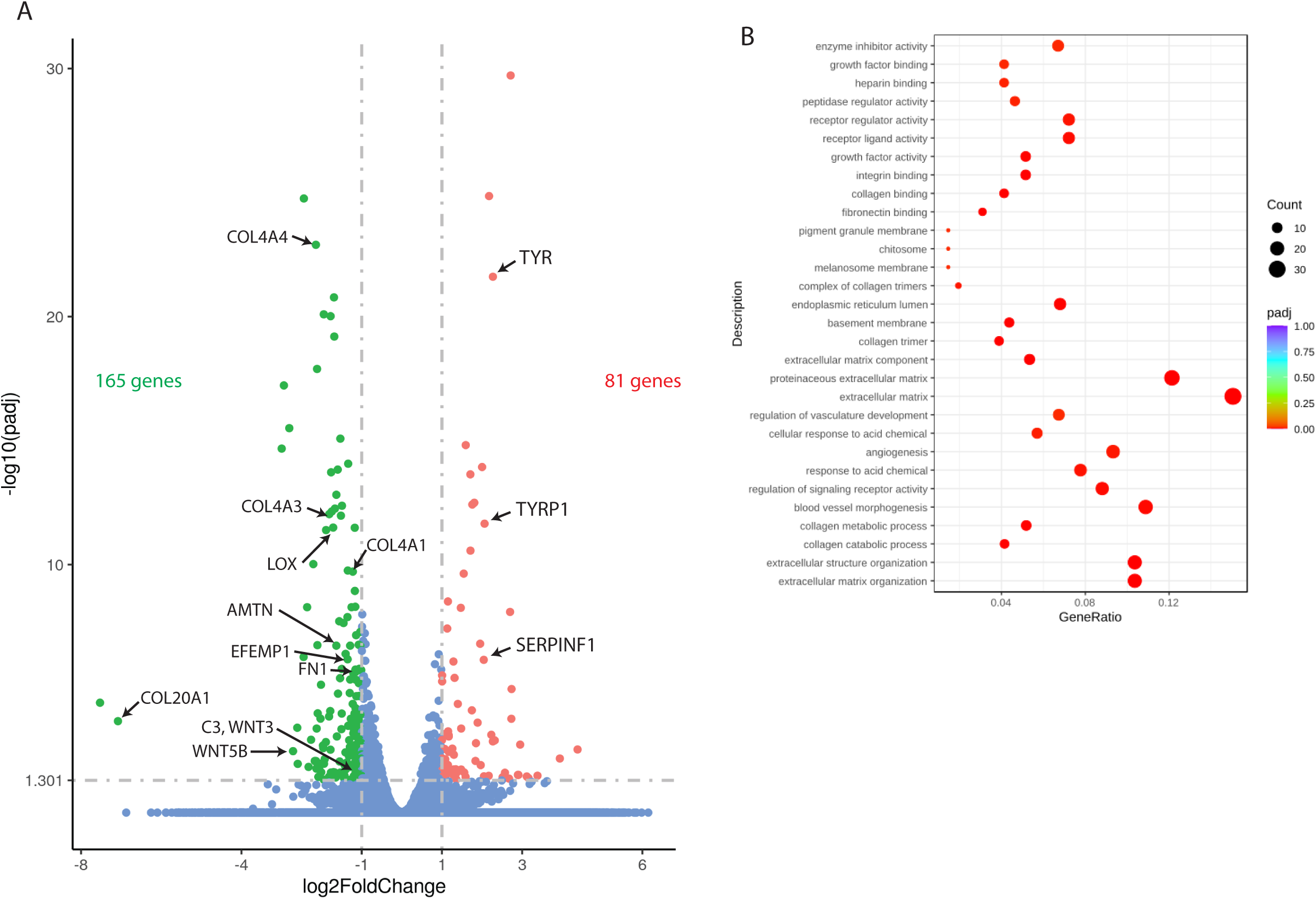
Low level, one-week CHIR99021 treatment reduces extracellular matrix (ECM) and collagen-associated transcripts while upregulating RPE differentiation-associated genes. (A) Bulk RNAseq analysis of ARPE-19 cells treated with CHIR99021 indicates significant transcript reduction (≥ 2-fold) of genes implicated in ECM formation and sub-RPE deposit formation while upregulating (≥ 2-fold) RPE differentiation genes. n = 3 independent experiments combined to produce these data, * padj < 0.05 using a negative binomial distribution model. (B) Pathway enrichment of significantly altered genes from RNAseq data set using Gene Ontology enrichment analysis.

### CHIR treatment reduces R345W F3-related pathology in vitro

Given the ability of CHIR to reduce several key phenomena which are associated with introduction of the R345W F3 ML/DHRD mutation in cells, humans, or mice (i.e., buildup of F3 (20, 34, 57), collagen IV deposition (58), increased MMP2 (4, 24), and activated C3 (4, 59, 60)), we decided to test whether this compound could serve as a multipronged approach to alleviate R345W F3-dependent pathology. To test this idea, initially we generated homozygous R345W knockin (KI) ARPE-19 cells using CRISPR in a similar manner as described previously (4), isolated clonal edited cells, and then tagged R345W F3 in those cells with HiBiT, as described in Figure 1A (Supplemental Figure 10, A-C). HiBiT edited WT and R345W cells were seeded on transwells as described previously (4, 9) to promote cell polarization and ECM deposition in serum-free media. Consistent with previous observations (4), under untreated conditions, HiBiT R345W F3 cells demonstrated significantly increased apical (1.81 ± 0.18 fold) and basal (1.46 ± 0.15 fold) MMP2 levels based on zymography (Figure 6A). CHIR treatment (1 μM, 1 week) significantly reduced both apically and basally secreted HiBiT WT F3 and HiBiT R345W F3 (Figure 6B) to a similar extent to cells plated on plastic culture dishes (cf. Figure 4C). CHIR treatment also significantly reduced MMP2 levels in the apical and basal media of HiBiT WT F3 and HiBiT R345W F3 cells by 16-17% in the apical media, and 29-31% in the basal media (Figure 6C).

**Figure 6:**
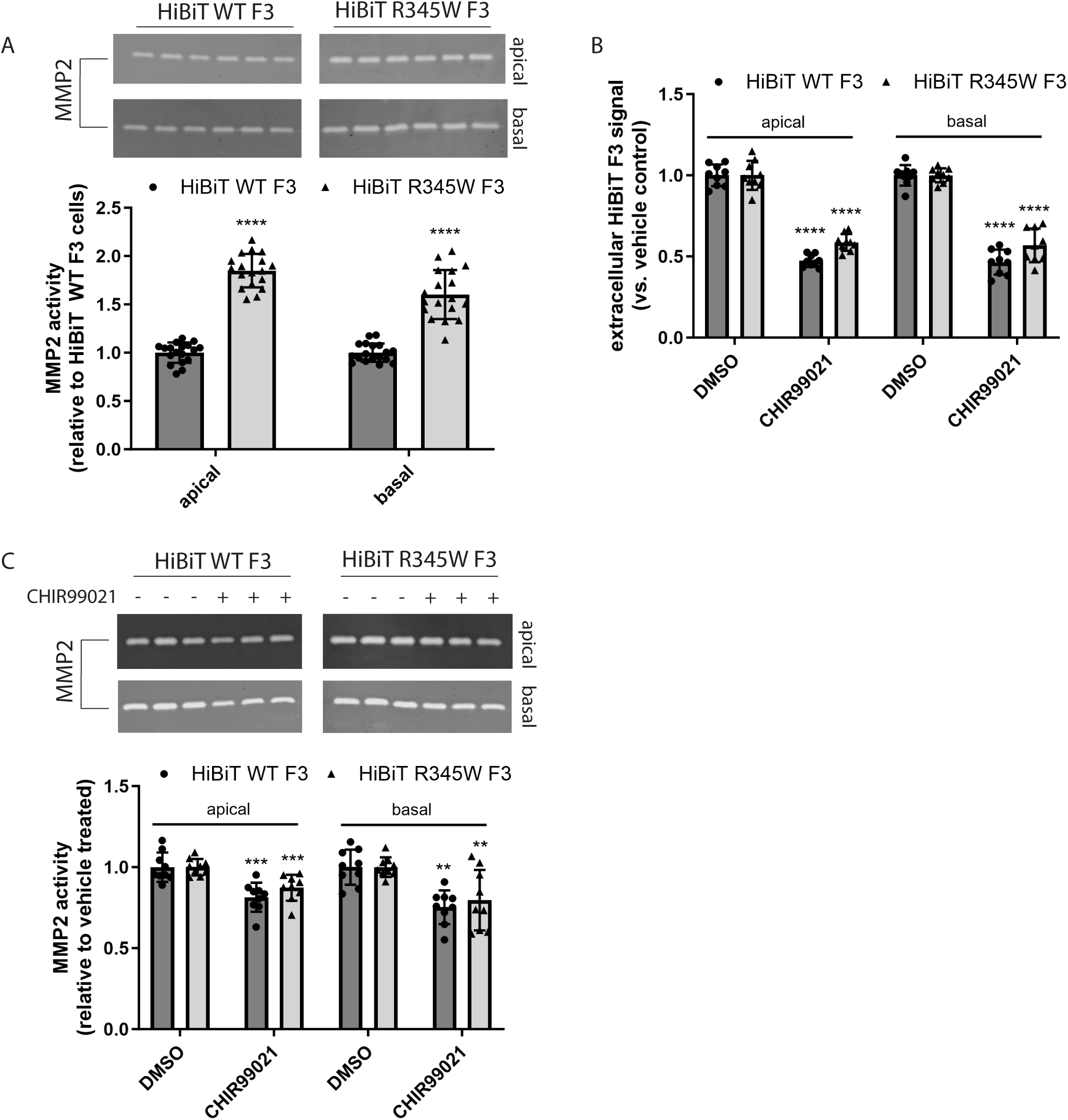
CHIR99021 reduces R345W F3-associated matrix metalloproteinase 2 (MMP2) alterations in culture. (A) ARPE-19 cells expressing HiBiT R345W F3 demonstrate significantly elevated apical and basal MMP2 activity levels compared to HiBiT WT F3 cells after one week on transwells. n = 3 independent experiments, performed in sextuplet biological replicates. **** p ≤ 0.0001, t-test vs. WT cells. (B) Treatment (one-week, 1 μM) with CHIR99021 significantly decreases HiBiT WT and R345W F3 both apically and basally in transwell format (C) while simultaneously significantly decreasing MMP2 activity. n = 3 independent experiments, performed in triplicate biological replicates. ** p ≤ 0.01, *** p ≤ 0.001, **** p ≤ 0.0001, t-test vs. each respective vehicle-treated control.

### CHIR enters the retina and demonstrates favorable pharmacokinetic (PK) properties

Previous studies using CHIR in preclinical neurological disorders (i.e., bipolar disorder and Huntington’s disease) have demonstrated that it can cross the blood-brain barrier (61) and provide beneficial effects in the brain (62). However, no studies have validated whether it can also efficiently cross into the retina. Thus, we next performed PK studies to test CHIR penetrance into the mouse eye. CHIR trihydrochloride (25 mg/kg i.p.) showed maximal distribution 30 min post i.p. dosing in plasma, liver, and retina (Table 3). In plasma and liver, CHIR was quickly metabolized/excreted (Table 3), yielding a plasma terminal half-life (T½) of 118 and 120 min, respectively, and a mean residence time (MRT) of 49 min and 94 min, respectively (Table 4). Yet, levels of CHIR in the retina persisted, recording a substantially higher terminal T½ of 405 min and MRT of 363 min (Table 4). Based on the vitreous volume of the mouse eye (∼4.4 μL (33)), we estimate that the peak concentration of CHIR in the eye is ∼1.3 μM after 30 min, falling to ∼1.1 μM after 3 h, 364 nM after 6 h, and 151 nM after 24 h. Thus, based on our cell culture observations, the concentration of CHIR falls within the range of concentrations capable to reduce F3 production, but not trigger detectable Wnt activation.

**Table 3:** Pharmacokinetic results of CHIR99021 in plasma, liver, and retina when administered via i.p.

| CHIR99021, 25 mpk IP |  |  |  |  |  |  |
| --- | --- | --- | --- | --- | --- | --- |
|  | Plasma |  | Liver* |  | Retina |  |
| Time (min) | Ave Conc (ng/mL) | Std Dev | Ave Conc (ng/g) | Std Dev | Ave Conc (ng/g) | Std Dev |
| 30 | 9250 | 2236 | 33337 | 3820 | 527 | 244 |
| 180 | 490 | 128 | 4681 | 1084 | 434 | 98 |
| 360 | 41 | 44 | 1165 | 692 | 148 | 24 |
| 960 | 0.3 | 0.0 | 6.1 | 3.7 | 41 | 14 |
| 1440 | 0.4 | 0.2 | 5.2 | 5.7 | 61 | 31 |

**Table 4:** CHIR99021 metabolism/excretion parameters post i.p. administration

| CHIR99021, 25mpk IP | Plasma | Liver* | Retina |
| --- | --- | --- | --- |
| Terminal T½ (min): | 118 | 120 | 405 |
| Tmax (min): | 30 | 30 | 30 |
| Cmax (ng/mL or ng/g): | 9250 | 33337 | 527 |
| AUClast (min*ng/mL or min*ng/g): | 929,394 ± 117,255 | 4,231,688 ± 272,680 | 213,348 ± 17,725 |
| Vz_F (mL/kg): | 4586 |  |  |
| CL_F (mL/min/kg): | 27 |  |  |
| MRT (min): | 49 | 94 | 362 |

### Prolonged in vivo administration of CHIR has no detrimental effect on retinal structure or function

Eight-month-old homozygous R345W (R345W^+/+^) F3 KI mice (25) were treated with CHIR (25 mg/kg i.p.) or vehicle every weekday for 1 mo. We observed no physical changes (e.g., size, coat condition, or behavior) in the mice after CHIR treatment, consistent with previous studies (61, 62). Mice were then evaluated for electroretinogram (ERG) functional changes in their outer retina (photoreceptor cells, a wave) or inner retina (ON bipolar cells, Muller cells, b wave) at various light intensities. CHIR-treated mice showed no difference in ERG readings (a- or b-wave) when compared to vehicle-treated mice (Figure 7, A and B). Consistent with these functional observations, histology demonstrated no structural changes across all retinal cell layers (Figure 7, C and D), reaffirming the ocular safety of prolonged systemic CHIR treatment.

**Figure 7:**
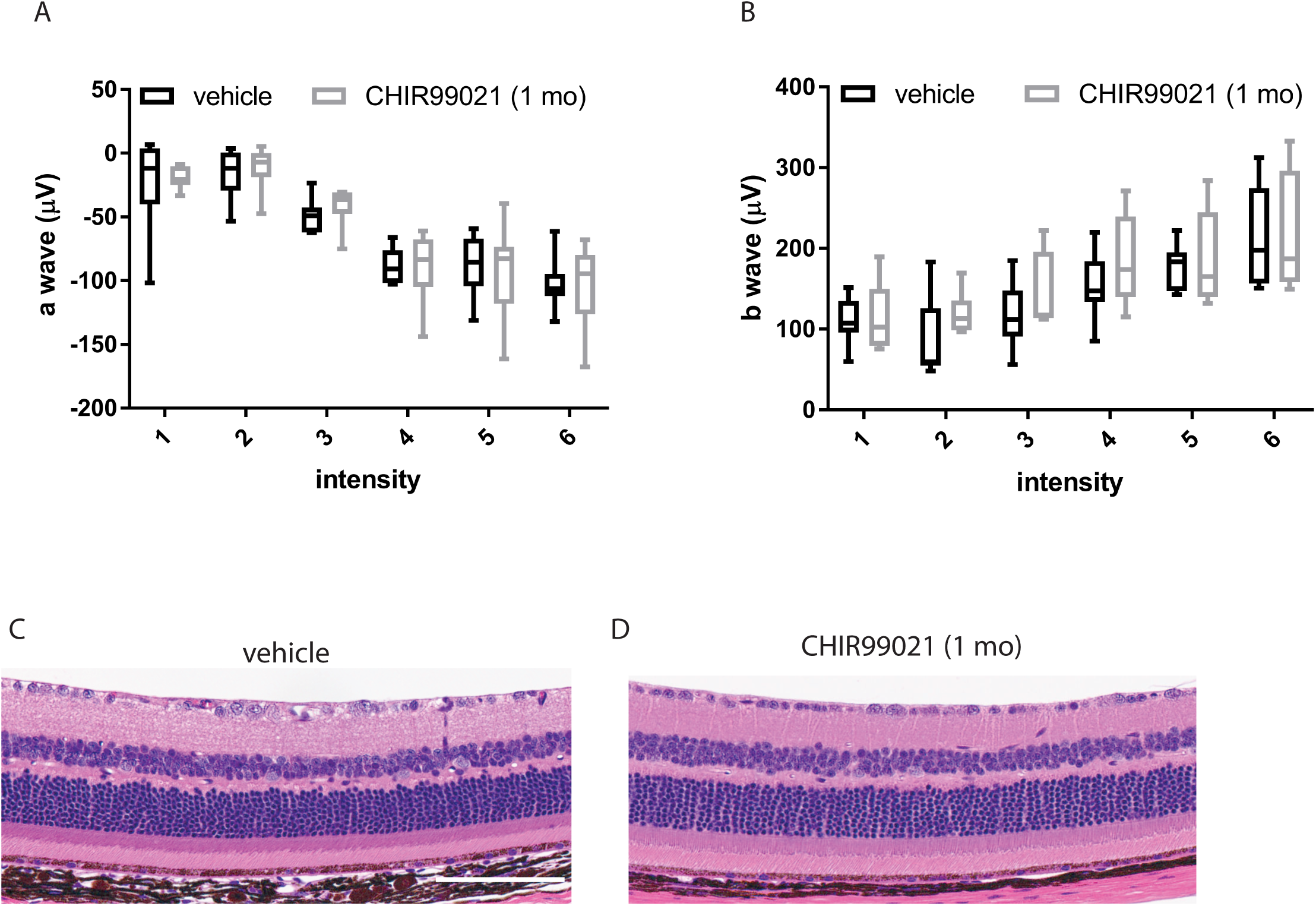
One-month CHIR99021 treatment does not affect retinal function or gross structure. (A,B) Eight-month-old R345W^+/+^ C57BL/6 mice were injected intraperitonially (i.p.) with vehicle (PBS) or CHIR99021 for one month (every weekday). Scotopic electroretinogram (ERG) readings demonstrated no difference between groups in either a-wave (outer retina, A) or b-wave (inner retina, B), values not significant by an ANOVA test. (C) After completion of ERG evaluation, mice were sacrificed and their eye prepared for H&E histology, which demonstrate no observable differences between the two groups. n = 4 mice/treatment group, 2 male, 2 female. Scale bar = 2 μm.

CHIR reduces basal laminar deposit (BLamD) formation in R345W^+/+^ mice. The canonical pathologic feature of the ML/DHRD macular dystrophy mouse model is the formation of sub-RPE BLamDs that increase with R345W F3 gene dosage and age (25, 57). Given the ability of CHIR to reduce the production of proteins associated with sub-RPE deposits (e.g., F3, FN1, AMTN, C3, and collagens) from cultured cells, we next asked whether CHIR treatment could prevent or slow the formation of BLamDs in vivo. Eight-month-old R345W^+/+^ mice were treated for 1 mo as described above. At this age, R345W^+/+^ mice form a few isolated BLamD, some of which begin to coalesce to form continuous BLamD across transmission electron microscopy (TEM) fields of view (FOV) (25, 57). Thus, the BLamD that form in these mice represent an early, but symptomatic stage of retinal disease progression (2). One month of CHIR treatment significantly reduced the number of BLamDs formed, decreasing number of FOVs containing deposits from 14.205% (125/880 FOV) in untreated mice to 4.167% (30/720 FOV) in treated mice (Figure 8, A-C). Example FOVs where BLamDs were not observed can be found in the Supplemental Material (Supplemental Figure 11, A and B). Additionally, in FOVs where BLamDs were observed, their average size was also significantly reduced from 0.817 µm^2^ in untreated mice to 0.416 µm^2^ in CHIR-treated mice (Figure 8D). These results are the first demonstration of small molecule-mediated reduction in BLamD formation in mice, paving the way for future studies in additional preclinical models of macular dystrophy/macular degeneration characterized by sub-RPE deposit formation.

**Figure 8:**
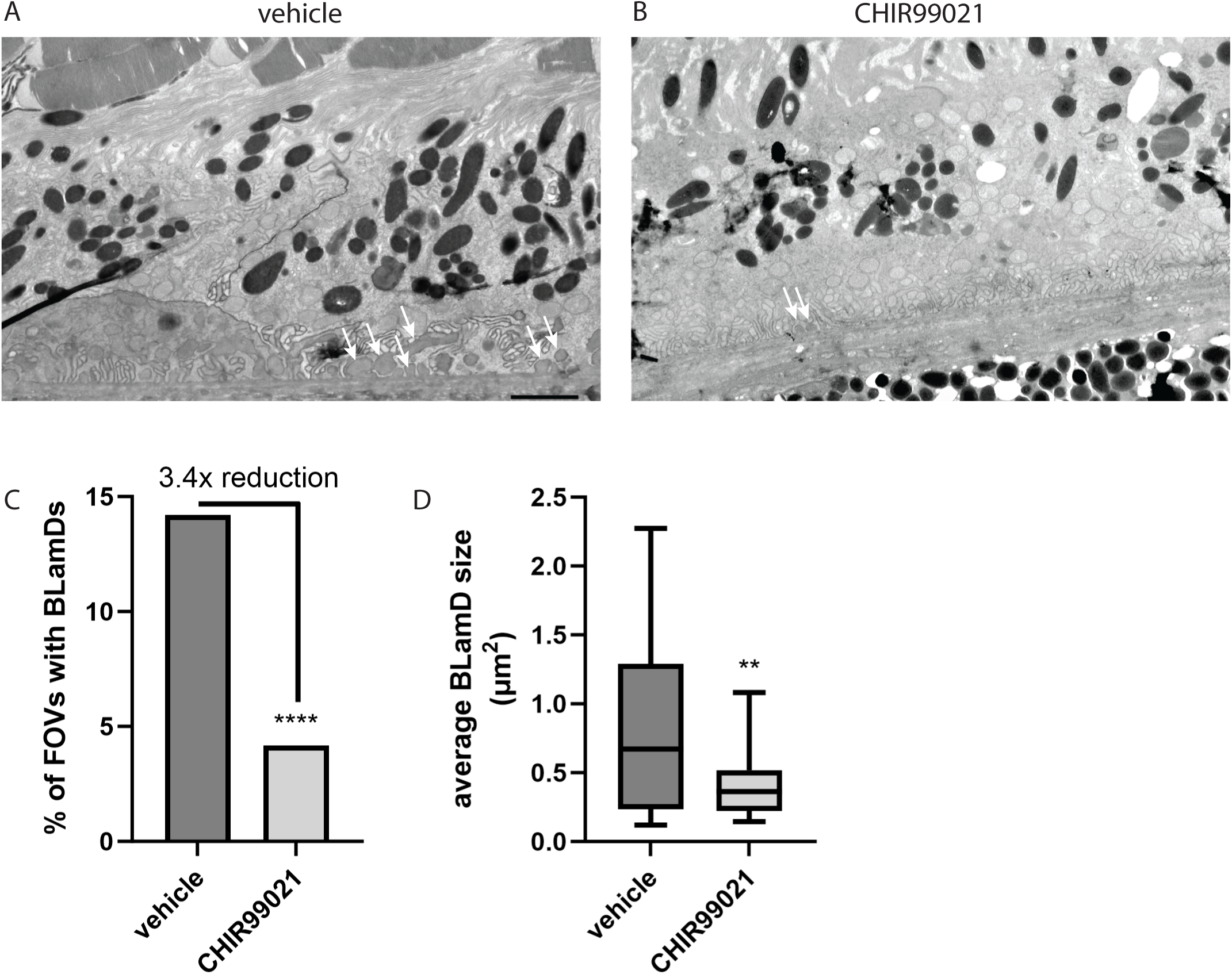
CHIR99021 significantly reduces the formation of basal laminar deposits (BLamD) *in vivo*. (A, B) After one month of vehicle (A) or CHIR99021 (B) treatment, nine-month-old mice were evaluated for BLamD (arrows) by transmission microscopy (TEM). Representative fields of view (FOV) of twenty-two fields are presented for vehicle or CHIR99021 treatment. (C, D) A masked observer systematically quantified number of FOV containing any BLamD (C), **** p ≤ 0.0001, chi-square test, as well as the average size of the BLamD, if at all present (D), ** p ≤ 0.01, t-test vs. vehicle-treated samples.

## DISCUSSION

Herein we have highlighted the first small molecule-based treatment that can prevent AMD-like pathology associated with ML/DHRD, an aggressive and currently incurable retinal disease that can affect individuals as young as 12 years of age (25). Natural history studies have indicated that ML/DHRD onset can vary substantially, even among siblings (63), which suggests that disease pathology is likely modifiable. Reaffirming this idea, at least one 62-year-old male harboring the R345W mutation was identified as asymptomatic (63). These observations are consistent with our findings that small molecules such as CHIR can substantially affect R345W-dependent pathology both in vitro and in vivo. While the prevalence of ML/DHRD is indeed rare with an unknown number of patients worldwide, like many inherited retinal degenerations (IRDs), ML/DHRD is likely underdiagnosed. In fact, in some European populations, it may comprise up to 0.9% of all IRDs (64), amounting to over three thousand affected individuals in Europe alone, representing a significant unmet patient population that could benefit from new therapeutic approaches.

More broadly, our observations that CHIR reduces BLamD formation in the ML/DHRD mouse model suggest that it may be possible that the main pathogenic feature of early and intermediate AMD (e.g., sub-RPE drusen deposits) is manipulatable using a single small molecule. Interestingly, there is precedence in translating pathology reduction in the ML/DHRD mice to meaningful benefit in humans with AMD. For example, a previous study indicated that genetic elimination of C3 also prevented BLamD formation in ML/DHRD mice (25). Just this year, pegcetacoplan (Syfovre) (65), a C3 peptide inhibitor, was approved as the only Food and Drug Administration (FDA)-approved drug to minimize progression of geographic atrophy in AMD. Thus, we are hopeful that our own encouraging findings in the ML/DHRD mouse model will contribute toward the development of therapeutic advances aimed at early and intermediate stages of AMD. We hypothesize that therapies such as CHIR have the potential to delay the advancement of early and intermediate stages of AMD (affecting ∼150 million individuals worldwide (66, 67)) to later stages such as geographic atrophy, and thus may be able to prevent eventual vision loss associated with dry AMD.

Our chemogenetic data strongly suggest that CHIR is primarily working through GSK3 inhibition. Yet, even after the identification of GSK3 40 years ago (68), its role(s) in regulating fundamental cellular processes including cellular architecture, differentiation, cell fate, gene expression, and energy metabolism, and others (69, 70) is still being defined. Given its wide range of targets and biological effects, perhaps it is not surprising that GSK3 overactivity has been implicated in diverse diseases including neurodegenerative diseases (e.g., Alzheimer’s disease (71, 72), Parkinson’s disease (73), and myotonic dystrophy (74)) as well as psychiatric disorders (e.g., unipolar and bipolar affective disorders (75, 76)). Accordingly, a plethora of GSK3 inhibitors have been developed (77) and safely used in multiple preclinical systems (77), even in models of retinitis pigmentosa (78, 79), optic nerve regeneration (80), and steroid-induced glaucoma (81). However, translating these observations into humans has been met with varying success (61, 82). As a result, LiCl, a commonly used antipsychotic (75, 76, 83), remains the only FDA-approved GSK3 inhibitor. Yet, its narrow therapeutic window all but prevents its use for neurodegenerative diseases (75, 76). Thus, there is a substantial opportunity to either develop more efficacious GSK3 inhibitors or to apply existing inhibitors for new diseases, such as ML/DHRD or AMD.

Given the wide number of GSK3 phosphorylation targets, it is likely that CHIR-mediated prevention of BLamD formation could originate from not just a single phenomenon such as reduction in F3, as we had envisioned, but rather a combination of reduced F3 burden, ECM compositional changes, gene expression alterations (49, 84–86), and possibly metabolic remodeling (87, 88). Moreover, it is not clear whether broad GSK3 inhibition across tissues (or the body, for that matter) is necessary for the BLamD prevention we observe, or if GSK3 inhibition in a particular retinal cell layer (i.e., in the neural retina vs. RPE vs. choroid) is sufficient to achieve BLamD reduction. Elucidation of these critical questions will enable more precise and targeted GSK3-based therapeutics while avoiding potential systemic side effects of broad GSK3 inhibition (82, 89). Ultimately, testing CHIR in non-human primates, or humans to reverse existing sub-RPE deposit formation will serve as a challenging, but necessary next endeavor.

The authors declare no conflicts of interest

## ACKNOWLEDGEMENTS

We thank Jessica Kilgore and Noelle Williams, Ph.D. (UT Southwestern Preclinical Pharmacology Core) for developing a quantitative CHIR99021 LC/MS assay and for their PK analysis of this compound in mouse tissue. We thank Andy Lemoff and the UT Southwestern Mass Spectrometry Core for running and analyzing our secreted proteome samples. JDH is the Larson Endowed Chair for Macular Degeneration Research (UMN). JDH is supported by a Macular Degeneration Research Grant from the Fichtenbaum Charitable Trust, the Roger and Dorothy Hirl Endowed Research Fund, the Edward N. and Della L. Thome Memorial Foundation Award in Age-Related Macular Degeneration Research, R01-EY027785, and P30-EY030413 (to the UT Southwestern Department of Ophthalmology). JDH is also a member of the Promega Advanced Academic Access Program, which provided reagent support for high-throughput screening.

## REFERENCES

1. Daniel S, Renwick M, Chau VQ, Datta S, Maddineni P, Zode G, et al. Fibulin-3 knockout mice demonstrate corneal dysfunction but maintain normal retinal integrity. Journal of molecular medicine. 2020;98(11):1639–56.

2. Sura AA, Chen L, Messinger JD, Swain TA, McGwin G, Jr., Freund KB, and Curcio CA. Measuring the Contributions of Basal Laminar Deposit and Bruch’s Membrane in Age-Related Macular Degeneration. Investigative ophthalmology & visual science. 2020;61(13):19.

3. Fernandez-Godino R, Pierce EA, and Garland DL. Extracellular Matrix Alterations and Deposit Formation in AMD. Adv Exp Med Biol. 2016;854:53–8.

4. Fernandez-Godino R, Bujakowska KM, and Pierce EA. Changes in extracellular matrix cause RPE cells to make basal deposits and activate the alternative complement pathway. Human molecular genetics. 2018;27(1):147–59.

5. Moore DJ, and Clover GM. The effect of age on the macromolecular permeability of human Bruch’s membrane. Investigative ophthalmology & visual science. 2001;42(12):2970–5.

6. Booij JC, Baas DC, Beisekeeva J, Gorgels TG, and Bergen AA. The dynamic nature of Bruch’s membrane. Progress in retinal and eye research. 2010;29(1):1–18.

7. Driver SGW, Jackson MR, Richter K, Tomlinson P, Brockway B, Halliday BJ, et al. Biallelic variants in EFEMP1 in a man with a pronounced connective tissue phenotype. European journal of human genetics : EJHG. 2020;28(4):445–52.

8. Bizzari S, El-Bazzal L, Nair P, Younan A, Stora S, Mehawej C, et al. Recessive marfanoid syndrome with herniation associated with a homozygous mutation in Fibulin-3. European journal of medical genetics. 2020;63(5):103869.

9. Woodard DR, Daniel S, Nakahara E, Abbas A, DiCesare SM, Collier GE, and Hulleman JD. A loss-of-function cysteine mutant in fibulin-3 (EFEMP1) forms aberrant extracellular disulfide-linked homodimers and alters extracellular matrix composition. Hum Mutat. 2022.

10. Meyer KJ, Davis LK, Schindler EI, Beck JS, Rudd DS, Grundstad AJ, et al. Genome-wide analysis of copy number variants in age-related macular degeneration. Hum Genet. 2011;129(1):91–100.

11. Cheng L, Chen C, Guo W, Liu K, Zhao Q, Lu P, et al. EFEMP1 Overexpression Contributes to Neovascularization in Age-Related Macular Degeneration. Front Pharmacol. 2020;11:547436.

12. Collantes ERA, Delfin MS, Fan B, Torregosa JMR, Siguan-Bell C, Florcruz NVG, et al. EFEMP1 rare variants cause familial juvenile-onset open-angle glaucoma. Hum Mutat. 2021.

13. Gupta V, Somarajan BI, Gupta S, Mahalingam K, Kumar M, and Singh A. Association of EFEMP1 with juvenile-onset open angle glaucoma in a patient with concomitant COL11A1-related Stickler syndrome. Ophthalmic genetics. 2022:1–5.

14. Mackay DS, Bennett TM, and Shiels A. Exome Sequencing Identifies a Missense Variant in EFEMP1 Co-Segregating in a Family with Autosomal Dominant Primary Open-Angle Glaucoma. PLoS One. 2015;10(7):e0132529.

15. Woodard DR, Nakahara E, and Hulleman JD. Clinically-identified C-terminal mutations in fibulin-3 are prone to misfolding and destabilization. Scientific reports. 2021;11(1):2998.

16. Stone EM, Lotery AJ, Munier FL, Heon E, Piguet B, Guymer RH, et al. A single EFEMP1 mutation associated with both Malattia Leventinese and Doyne honeycomb retinal dystrophy. Nature genetics. 1999;22(2):199–202.

17. Hulleman JD. Malattia Leventinese/Doyne Honeycomb Retinal Dystrophy: Similarities to Age-Related Macular Degeneration and Potential Therapies. Adv Exp Med Biol. 2016;854:153–8.

18. Roybal CN, Marmorstein LY, Vander Jagt DL, and Abcouwer SF. Aberrant accumulation of fibulin-3 in the endoplasmic reticulum leads to activation of the unfolded protein response and VEGF expression. Investigative ophthalmology & visual science. 2005;46(11):3973–9.

19. Hulleman JD, Balch WE, and Kelly JW. Translational attenuation differentially alters the fate of disease-associated fibulin proteins. FASEB J. 2012;26(11):4548–60.

20. Hulleman JD, Kaushal S, Balch WE, and Kelly JW. Compromised mutant EFEMP1 secretion associated with macular dystrophy remedied by proteostasis network alteration. Mol Biol Cell. 2011;22(24):4765–75.

21. Hulleman JD, and Kelly JW. Genetic ablation of N-linked glycosylation reveals two key folding pathways for R345W fibulin-3, a secreted protein associated with retinal degeneration. FASEB J. 2015;29(2):565–75.

22. Zhou M, Weber SR, Zhao Y, Chen H, Barber AJ, Grillo SL, et al. Expression of R345W-Fibulin-3 Induces Epithelial-Mesenchymal Transition in Retinal Pigment Epithelial Cells. Front Cell Dev Biol. 2020;8:469.

23. Zhou M, Zhao Y, Weber SR, Gates C, Carruthers NJ, Chen H, et al. Extracellular vesicles from retinal pigment epithelial cells expressing R345W-Fibulin-3 induce epithelial-mesenchymal transition in recipient cells. J Extracell Vesicles. 2023;12(10):e12373.

24. Fernandez-Godino R, Garland DL, and Pierce EA. A local complement response by RPE causes early-stage macular degeneration. Human molecular genetics. 2015;24(19):5555–69.

25. Fu L, Garland D, Yang Z, Shukla D, Rajendran A, Pearson E, et al. The R345W mutation in EFEMP1 is pathogenic and causes AMD-like deposits in mice. Human molecular genetics. 2007;16(20):2411–22.

26. Stanton JB, Marmorstein AD, Zhang Y, and Marmorstein LY. Deletion of Efemp1 Is Protective Against the Development of Sub-RPE Deposits in Mouse Eyes. Investigative ophthalmology & visual science. 2017;58(3):1455–61.

27. Nguyen A, and Hulleman JD. Evidence of Alternative Cystatin C Signal Sequence Cleavage Which Is Influenced by the A25T Polymorphism. PLoS One. 2016;11(2):e0147684.

28. Zhang JH, Chung TD, and Oldenburg KR. A Simple Statistical Parameter for Use in Evaluation and Validation of High Throughput Screening Assays. J Biomol Screen. 1999;4(2):67–73.

29. Vu KT, and Hulleman JD. An inducible form of Nrf2 confers enhanced protection against acute oxidative stresses in RPE cells. Experimental eye research. 2017;164:31–6.

30. Vu KT, Zhang F, and Hulleman JD. Conditional, Genetically Encoded, Small Molecule-Regulated Inhibition of NFkappaB Signaling in RPE Cells. Investigative ophthalmology & visual science. 2017;58(10):4126–37.

31. Fuerer C, and Nusse R. Lentiviral vectors to probe and manipulate the Wnt signaling pathway. PLoS One. 2010;5(2):e9370.

32. Sun N, Shibata B, Hess JF, and FitzGerald PG. An alternative means of retaining ocular structure and improving immunoreactivity for light microscopy studies. Molecular vision. 2015;21:428–42.

33. Datta S, Renwick M, Chau VQ, Zhang F, Nettesheim ER, Lipinski DM, and Hulleman JD. A Destabilizing Domain Allows for Fast, Noninvasive, Conditional Control of Protein Abundance in the Mouse Eye - Implications for Ocular Gene Therapy. Investigative ophthalmology & visual science. 2018;59(12):4909–20.

34. Marmorstein LY, Munier FL, Arsenijevic Y, Schorderet DF, McLaughlin PJ, Chung D, et al. Aberrant accumulation of EFEMP1 underlies drusen formation in Malattia Leventinese and age-related macular degeneration. Proceedings of the National Academy of Sciences of the United States of America. 2002;99(20):13067–72.

35. Dixon AS, Schwinn MK, Hall MP, Zimmerman K, Otto P, Lubben TH, et al. NanoLuc Complementation Reporter Optimized for Accurate Measurement of Protein Interactions in Cells. ACS Chem Biol. 2016;11(2):400–8.

36. Hu B, Thirtamara-Rajamani KK, Sim H, and Viapiano MS. Fibulin-3 is uniquely upregulated in malignant gliomas and promotes tumor cell motility and invasion. Mol Cancer Res. 2009;7(11):1756–70.

37. Bharti K, den Hollander AI, Lakkaraju A, Sinha D, Williams DS, Finnemann SC, et al. Cell culture models to study retinal pigment epithelium-related pathogenesis in age-related macular degeneration. Experimental eye research. 2022;222:109170.

38. Sisask G, Marsell R, Sundgren-Andersson A, Larsson S, Nilsson O, Ljunggren O, and Jonsson KB. Rats treated with AZD2858, a GSK3 inhibitor, heal fractures rapidly without endochondral bone formation. Bone. 2013;54(1):126–32.

39. Ring DB, Johnson KW, Henriksen EJ, Nuss JM, Goff D, Kinnick TR, et al. Selective glycogen synthase kinase 3 inhibitors potentiate insulin activation of glucose transport and utilization in vitro and in vivo. Diabetes. 2003;52(3):588–95.

40. Beurel E, Grieco SF, and Jope RS. Glycogen synthase kinase-3 (GSK3): regulation, actions, and diseases. Pharmacol Ther. 2015;148:114–31.

41. De Simone A, Tumiatti V, Andrisano V, and Milelli A. Glycogen Synthase Kinase 3beta: A New Gold Rush in Anti-Alzheimer’s Disease Multitarget Drug Discovery? J Med Chem. 2021;64(1):26–41.

42. Kisseleff E, Vigouroux RJ, Hottin C, Lourdel S, Thomas L, Shah P, et al. Glycogen Synthase Kinase 3 Regulates the Genesis of Displaced Retinal Ganglion Cells3. eNeuro. 2021;8(5).

43. Timpl R, Sasaki T, Kostka G, and Chu ML. Fibulins: a versatile family of extracellular matrix proteins. Nature reviews. 2003;4(6):479–89.

44. Vijay GV, Zhao N, Den Hollander P, Toneff MJ, Joseph R, Pietila M, et al. GSK3beta regulates epithelial-mesenchymal transition and cancer stem cell properties in triple-negative breast cancer. Breast Cancer Res. 2019;21(1):37.

45. Bachelder RE, Yoon SO, Franci C, de Herreros AG, and Mercurio AM. Glycogen synthase kinase-3 is an endogenous inhibitor of Snail transcription: implications for the epithelial-mesenchymal transition. The Journal of cell biology. 2005;168(1):29–33.

46. Wu D, and Pan W. GSK3: a multifaceted kinase in Wnt signaling. Trends in biochemical sciences. 2010;35(3):161–8.

47. Liu C, Li Y, Semenov M, Han C, Baeg GH, Tan Y, et al. Control of beta-catenin phosphorylation/degradation by a dual-kinase mechanism. Cell. 2002;108(6):837–47.

48. Shu DY, Butcher E, and Saint-Geniez M. EMT and EndMT: Emerging Roles in Age-Related Macular Degeneration. International journal of molecular sciences. 2020;21(12).

49. Rajapakse D, Peterson K, Mishra S, Fan J, Lerner J, Campos M, and Wistow G. Amelotin is expressed in retinal pigment epithelium and localizes to hydroxyapatite deposits in dry age-related macular degeneration. Transl Res. 2020;219:45–62.

50. Miller CG, Budoff G, Prenner JL, and Schwarzbauer JE. Minireview: Fibronectin in retinal disease. Exp Biol Med (Maywood). 2017;242(1):1–7.

51. Xiao Q, and Ge G. Lysyl oxidase, extracellular matrix remodeling and cancer metastasis. Cancer Microenviron. 2012;5(3):261–73.

52. Wu Y, Ginther C, Kim J, Mosher N, Chung S, Slamon D, and Vadgama JV. Expression of Wnt3 activates Wnt/beta-catenin pathway and promotes EMT-like phenotype in trastuzumab-resistant HER2-overexpressing breast cancer cells. Mol Cancer Res. 2012;10(12):1597–606.

53. Wang B, Tang Z, Gong H, Zhu L, and Liu X. Wnt5a promotes epithelial-to-mesenchymal transition and metastasis in non-small-cell lung cancer. Biosci Rep. 2017;37(6).

54. Maruotti J, Sripathi SR, Bharti K, Fuller J, Wahlin KJ, Ranganathan V, et al. Small-molecule-directed, efficient generation of retinal pigment epithelium from human pluripotent stem cells. Proceedings of the National Academy of Sciences of the United States of America. 2015;112(35):10950–5.

55. Zhu D, Deng X, Spee C, Sonoda S, Hsieh CL, Barron E, et al. Polarized secretion of PEDF from human embryonic stem cell-derived RPE promotes retinal progenitor cell survival. Investigative ophthalmology & visual science. 2011;52(3):1573–85.

56. Leach LL, Buchholz DE, Nadar VP, Lowenstein SE, and Clegg DO. Canonical/beta-catenin Wnt pathway activation improves retinal pigmented epithelium derivation from human embryonic stem cells. Investigative ophthalmology & visual science. 2015;56(2):1002–13.

57. Marmorstein LY, McLaughlin PJ, Peachey NS, Sasaki T, and Marmorstein AD. Formation and progression of sub-retinal pigment epithelium deposits in Efemp1 mutation knock-in mice: a model for the early pathogenic course of macular degeneration. Human molecular genetics. 2007;16(20):2423–32.

58. Sohn EH, Wang K, Thompson S, Riker MJ, Hoffmann JM, Stone EM, and Mullins RF. Comparison of drusen and modifying genes in autosomal dominant radial drusen and age-related macular degeneration. Retina. 2015;35(1):48–57.

59. Fernandez-Godino R, and Pierce EA. C3a triggers formation of sub-retinal pigment epithelium deposits via the ubiquitin proteasome pathway. Scientific reports. 2018;8(1):9679.

60. Fernandez-Godino R. Alterations in Extracellular Matrix/Bruch’s Membrane Can Cause the Activation of the Alternative Complement Pathway via Tick-Over. Adv Exp Med Biol. 2018;1074:29–35.

61. Pan JQ, Lewis MC, Ketterman JK, Clore EL, Riley M, Richards KR, et al. AKT kinase activity is required for lithium to modulate mood-related behaviors in mice. Neuropsychopharmacology : official publication of the American College of Neuropsychopharmacology. 2011;36(7):1397–411.

62. Hu D, Sun X, Magpusao A, Fedorov Y, Thompson M, Wang B, et al. Small-molecule suppression of calpastatin degradation reduces neuropathology in models of Huntington’s disease. Nature communications. 2021;12(1):5305.

63. Michaelides M, Jenkins SA, Brantley MA, Jr., Andrews RM, Waseem N, Luong V, et al. Maculopathy due to the R345W substitution in fibulin-3: distinct clinical features, disease variability, and extent of retinal dysfunction. Investigative ophthalmology & visual science. 2006;47(7):3085–97.

64. Pontikos N, Arno G, Jurkute N, Schiff E, Ba-Abbad R, Malka S, et al. Genetic Basis of Inherited Retinal Disease in a Molecularly Characterized Cohort of More Than 3000 Families from the United Kingdom. Ophthalmology. 2020;127(10):1384–94.

65. Kolev M, Barbour T, Baver S, Francois C, and Deschatelets P. With complements: C3 inhibition in the clinic. Immunol Rev. 2023;313(1):358–75.

66. Colijn JM, Buitendijk GHS, Prokofyeva E, Alves D, Cachulo ML, Khawaja AP, et al. Prevalence of Age-Related Macular Degeneration in Europe: The Past and the Future. Ophthalmology. 2017;124(12):1753–63.

67. Wong WL, Su X, Li X, Cheung CM, Klein R, Cheng CY, and Wong TY. Global prevalence of age-related macular degeneration and disease burden projection for 2020 and 2040: a systematic review and meta-analysis. The Lancet Global health. 2014;2(2):e106–16.

68. Embi N, Rylatt DB, and Cohen P. Glycogen synthase kinase-3 from rabbit skeletal muscle. Separation from cyclic-AMP-dependent protein kinase and phosphorylase kinase. European journal of biochemistry / FEBS. 1980;107(2):519–27.

69. Kaidanovich-Beilin O, and Woodgett JR. GSK-3: Functional Insights from Cell Biology and Animal Models. Front Mol Neurosci. 2011;4:40.

70. Cole AR. GSK3 as a Sensor Determining Cell Fate in the Brain. Front Mol Neurosci. 2012;5:4.

71. Hooper C, Killick R, and Lovestone S. The GSK3 hypothesis of Alzheimer’s disease. Journal of neurochemistry. 2008;104(6):1433–9.

72. Griebel G, Stemmelin J, Lopez-Grancha M, Boulay D, Boquet G, Slowinski F, et al. The selective GSK3 inhibitor, SAR502250, displays neuroprotective activity and attenuates behavioral impairments in models of neuropsychiatric symptoms of Alzheimer’s disease in rodents. Scientific reports. 2019;9(1):18045.

73. Li J, Ma S, Chen J, Hu K, Li Y, Zhang Z, et al. GSK-3beta Contributes to Parkinsonian Dopaminergic Neuron Death: Evidence From Conditional Knockout Mice and Tideglusib. Front Mol Neurosci. 2020;13:81.

74. Jones K, Wei C, Iakova P, Bugiardini E, Schneider-Gold C, Meola G, et al. GSK3beta mediates muscle pathology in myotonic dystrophy. The Journal of clinical investigation. 2012;122(12):4461–72.

75. Schou M. Forty years of lithium treatment. Arch Gen Psychiatry. 1997;54(1):9–13; discussion 4-5.

76. Moncrieff J. Forty years of lithium treatment. Arch Gen Psychiatry. 1998;55(1):92–3.

77. Arciniegas Ruiz SM, and Eldar-Finkelman H. Glycogen Synthase Kinase-3 Inhibitors: Preclinical and Clinical Focus on CNS-A Decade Onward. Front Mol Neurosci. 2021;14:792364.

78. Marchena M, Villarejo-Zori B, Zaldivar-Diez J, Palomo V, Gil C, Hernandez-Sanchez C, et al. Small molecules targeting glycogen synthase kinase 3 as potential drug candidates for the treatment of retinitis pigmentosa. J Enzyme Inhib Med Chem. 2017;32(1):522–6.

79. Sanchez-Cruz A, Villarejo-Zori B, Marchena M, Zaldivar-Diez J, Palomo V, Gil C, et al. Modulation of GSK-3 provides cellular and functional neuroprotection in the rd10 mouse model of retinitis pigmentosa. Mol Neurodegener. 2018;13(1):19.

80. Leibinger M, Andreadaki A, Golla R, Levin E, Hilla AM, Diekmann H, and Fischer D. Boosting CNS axon regeneration by harnessing antagonistic effects of GSK3 activity. Proceedings of the National Academy of Sciences of the United States of America. 2017;114(27):E5454–E63.

81. Sugali CK, Rayana NP, Dai J, Peng M, Harris SL, Webber HC, et al. The Canonical Wnt Signaling Pathway Inhibits the Glucocorticoid Receptor Signaling Pathway in the Trabecular Meshwork. Am J Pathol. 2021;191(6):1020–35.

82. Hottin C, Perron M, and Roger JE. GSK3 Is a Central Player in Retinal Degenerative Diseases but a Challenging Therapeutic Target. Cells. 2022;11(18).

83. Cohen P, and Goedert M. GSK3 inhibitors: development and therapeutic potential. Nat Rev Drug Discov. 2004;3(6):479–87.

84. Winkler TW, Grassmann F, Brandl C, Kiel C, Gunther F, Strunz T, et al. Genome-wide association meta-analysis for early age-related macular degeneration highlights novel loci and insights for advanced disease. BMC Med Genomics. 2020;13(1):120.

85. Holliday EG, Smith AV, Cornes BK, Buitendijk GH, Jensen RA, Sim X, et al. Insights into the genetic architecture of early stage age-related macular degeneration: a genome-wide association study meta-analysis. PLoS One. 2013;8(1):e53830.

86. Thompson RB, Reffatto V, Bundy JG, Kortvely E, Flinn JM, Lanzirotti A, et al. Identification of hydroxyapatite spherules provides new insight into subretinal pigment epithelial deposit formation in the aging eye. Proceedings of the National Academy of Sciences of the United States of America. 2015;112(5):1565–70.

87. Jellusova J, Cato MH, Apgar JR, Ramezani-Rad P, Leung CR, Chen C, et al. Gsk3 is a metabolic checkpoint regulator in B cells. Nat Immunol. 2017;18(3):303–12.

88. He L, Gomes AP, Wang X, Yoon SO, Lee G, Nagiec MJ, et al. mTORC1 Promotes Metabolic Reprogramming by the Suppression of GSK3-Dependent Foxk1 Phosphorylation. Mol Cell. 2018;70(5):949–60 e4.

89. Gitlin M. Lithium side effects and toxicity: prevalence and management strategies. Int J Bipolar Disord. 2016;4(1):27.

