## Supplemental Data for "GSK3 inhibition reduces ECM production and prevents age-related macular degeneration-like pathology"

The authors declare no conflicts of interest

**SUPPLEMENTAL METHODS**

HiBiT mouse F3 cell line generation. To add HiBiT to mouse F3, NIH-3T3 fibroblasts (ATCC CRL-1658) were used. Similar to the procedure described above, a 2xFLAG VS HiBiT sequence was inserted immediately after the Ser17 residue of mouse F3 (Supplemental Figure 5, A and B). Unlike in *EFEMP1*, no mutation of the PAM site was required in *Efemp1* due to significant disruption of the gRNA sequence after HiBiT insertion (Supplemental Figure 6A). RNP, ssODN, and electroporation enhancer were introduced into the NIH-3T3 cells (1400 V, 20 ms, 2 pulses, Neon Transfection System). After electroporation, cells were incubated in antibiotic-free media containing HDR Enhancer V3 (1 μM, IDT) for 48 h to promote homology-directed repair. Heterogenous cultures were expanded and assayed for secreted HiBiT via a luciferase assay.

HiBiT fibulin-5 (F5) cell line generation. As a control cell line for HiBiT F3 cells, we appended HiBiT onto endogenous fibulin-5 (FBLN5, DANCE, or F5), a highly homologous protein to F3 (19), in a similar manner, immediately after the F5 signal sequence cleavage after residue Ala23 (Supplemental Figure 8A). No mutation of the PAM site was required in *FBLN5* due to significant disruption of the gRNA sequence after HiBiT insertion (Table S1). RNP, ssODN, and electroporation enhancer were introduced into ARPE-19 cells using electroporation (1400V, 20 ms, 2 pulses, Neon Transfection System). After electroporation, cells were incubated in antibiotic-free media containing HDR Enhancer V3 (1 μM, IDT) for 48 h to promote homology-directed repair. Heterogenous cultures were expanded and assayed for secreted HiBiT via a luciferase assay.

### SUPPLEMENTAL FIGURE LEGENDS

**Supplemental Figure 1: Introduction of the 2xFLAG-VS-HiBiT tag is not predicted to alter human F3 signal sequence cleavage.** (A) SignalP5.0 predicted signal sequence cleavage of human wild-type (WT) F3 and (B) human F3 after introduction of the 2xFLAG-VS-HiBiT tag.

**Supplemental Figure 2: Example data from HTS plate indicating assay uniformity and identification of potential hit compounds.** (A) Example plate data from a whole-well HiBiT assay mock screen performed with DMSO and BFA (as a positive control) to identify an assay Z'-score. (B, C) Representative screening data identifying a potential F3 reducer (B) or a potential F3 enhancer (C).

**Supplemental Figure 3: Select hit compounds identified in the primary screen were confirmed in dose-response.** Seven hit compounds were tested at 0.2 nM – 50 μM for 24 h. All hits reproduced and demonstrated varying levels of dose-responsiveness.

**Supplemental Figure 4: Verification of CHIR99021 activity toward F3 in a human non-RPE cell line.** F3 in primary human dermal fibroblasts was edited with a HiBiT tag followed by treatment with CHIR99021 for 72 h. $n \geq 3$ independent experiments with mean of each experiment presented as a single data point in this graph. ** $p < 0.01$, one sample t-test vs. hypothetical mean of 1 (i.e., unchanged).

**Supplemental Figure 5: Introduction of the 2xFLAG-VS-HiBiT tag is not predicted to alter mouse F3 signal sequence cleavage.** (A) SignalP5.0 predicted signal sequence cleavage of mouse (mus) wild-type (WT) F3 and (B) human F3 after introduction of the 2xFLAG-VS-HiBiT tag.

**Supplemental Figure 6: HiBiT editing of mouse fibroblast NIH-3T3 cells followed by treatment with CHIR99021 demonstrates cross species activity of the compound.** (A) Schematic demonstrating the 2xFLAG-VS-HiBiT F3 design for genomic insertion into the mouse genome. The upward arrow indicates the predicted signal sequence cleavage site based on SignalP5.0 prediction. (B) The origins of the HiBiT signal were confirmed to be from mouse F3 as demonstrated by siRNA knockdown experiments. n = 3 independent experiments, *** $p < 0.001$, t-test vs. non-targeting siRNA. (C) Seventy-two hour treatment with CHIR99021 reduces F3 production in mouse cells. n = 3 independent experiments, ** $p < 0.01$, *** $p < 0.001$, t-test vs. vehicle-treated samples.

**Supplemental Figure 7: Introduction of the 2xFLAG-VS-HiBiT tag is not predicted to alter human fibulin-5 (F5) signal sequence cleavage.** (A) SignalP5.0 predicted signal sequence cleavage of human wild-type (WT) F5 and (B) human F5 after introduction of the 2xFLAG-VS-HiBiT tag.

**Supplemental Figure 8: Production of HiBiT-tagged human F5 is also reduced by CHIR99021 treatment.** (A) Schematic demonstrating the design of 2xFLAG-VS-HiBiT genomic insertion onto the N-terminus of human F5 in ARPE-19 cells. The origins of the resulting HiBiT signal were confirmed to be F5 gene expression-dependent using siRNA. Representative data of n = 3 independent experiments, average ± S.D. of technical triplicates. *** $p < 0.001$, t-test vs. non-targeting siRNA. (C) HiBiT F5-expressing ARPE-19 cells were treated with CHIR99021 for 72 h, followed by a HiBiT assay on conditioned media. n = 3 independent experiments performed in biological duplicates each time, ** $p < 0.01$, *** $p < 0.001$, t-test vs. vehicle-treated samples. (D) One-week CHIR99021 treatment also leads to reduced extracellular and intracellular F5 levels. n = 3 independent experiments performed in biological triplicate, *** $p < 0.001$, t-test vs. vehicle-treated samples.

**Supplemental Figure 9: Low level, 1-week CHIR99021 treatment does not substantially affect the secreted proteome from ARPE-19 cells.** Total concentrated secreted protein after 1 week of CHIR99021 treatment silver stained to visualize differences in overall protein abundance. Representative data of three independent experiments shown in biological duplicate.

**Supplemental Figure 10: Design and validation of R345W$^{+/+}$ ARPE-19 knockin cells and their subsequent HiBiT editing.** (A) Design of the R345W knockin strategy using CRISPR/Cas9 editing and homology-directed repair. (B) Genomic DNA from single colonies was isolated and validated to be homozygous ($^{+/+}$) for the R345W mutation. (C) The validated R345W$^{+/+}$ clone was then edited to include the 2xFLAG-VS-HiBiT tag.

**Supplemental Figure 11: Not all TEM fields of view show BLamD.** Some TEM fields were found to contain no BLamDs in both (A) vehicle (A) and CHIR99021-treated mice (B). Note the healthy basal infoldings in these images also. Scale bar = 2 $\mu$m.

**Supplemental Figure 12: Full gel and blot images for the indicated figure.**

**Supplemental Table 1: CRISPR reagent sequences used for guide RNAs, PAM sites, and ssODN design for insertion of the 2xFLAG-VS-HiBiT tag.**

**Supplemental Table 2: Primer sequences for exon 2 amplification and gDNA verification of 2xFLAG-VS-HiBiT insertion into the F3 gene.**

**Supplemental Table 3: siRNA sense sequences used for confirming the origins of the HiBiT signal in generated cell lines.**

**Supplemental Table 4: Processed data received from HTS with eight hit compounds of interest and additional information found about MOA, SMILES, HiBiT readout, etc.**

**Supplemental Table 5: Top 50 increased proteins between DMSO and CHIR treated cells.**

**Supplemental Table 6: GO analysis of top proteins and their involvement in cellular composition.**

**Supplemental Table 7: RNA-seq genes identified to have ≥ 2-fold decrease between CHIR treated and DMSO treated cells.**

**Supplemental Table 8: RNA-seq genes identified to have ≥ 2-fold increase between CHIR treated and DMSO treated cells.**

Supplemental Figure 1.

A

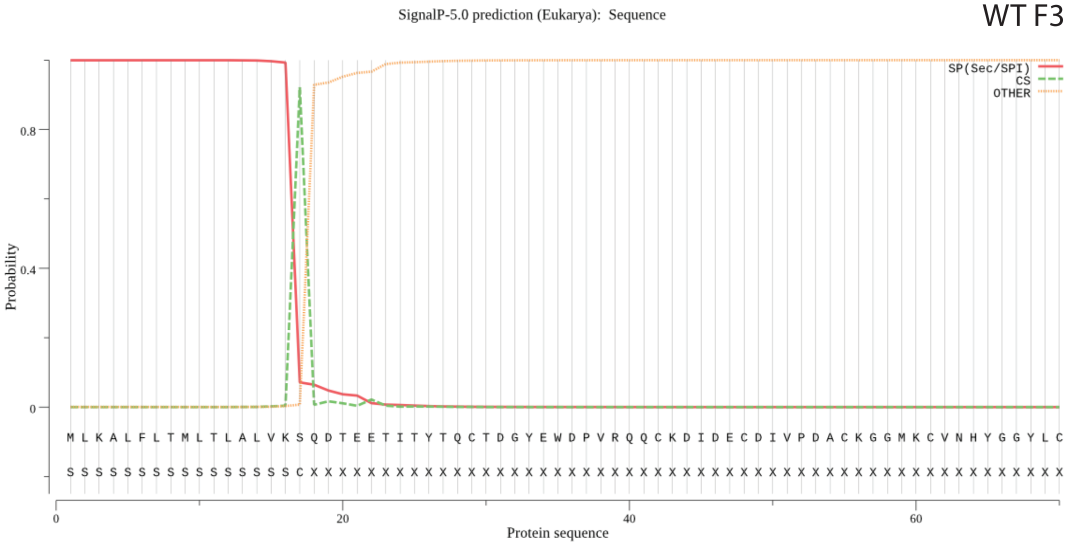


B

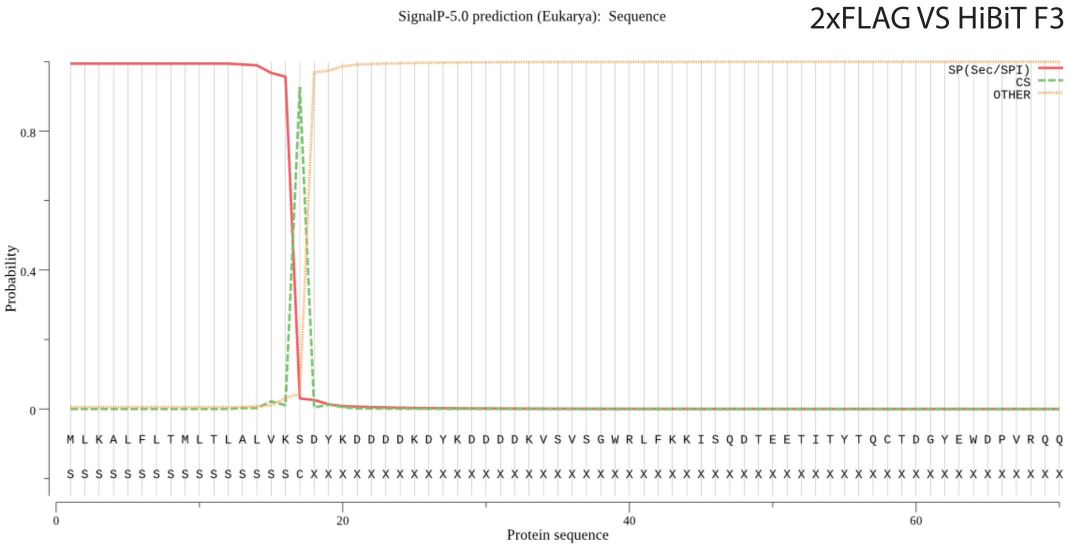

Supplemental Figure 2.

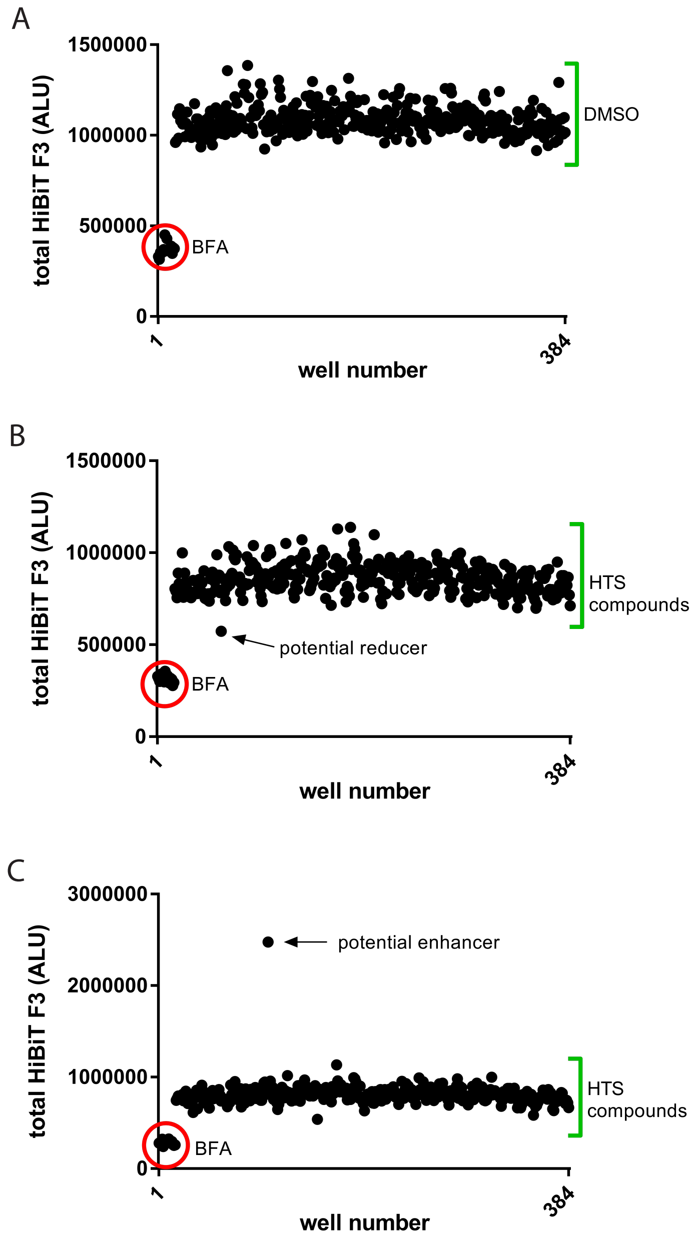

Supplemental Figure 3.

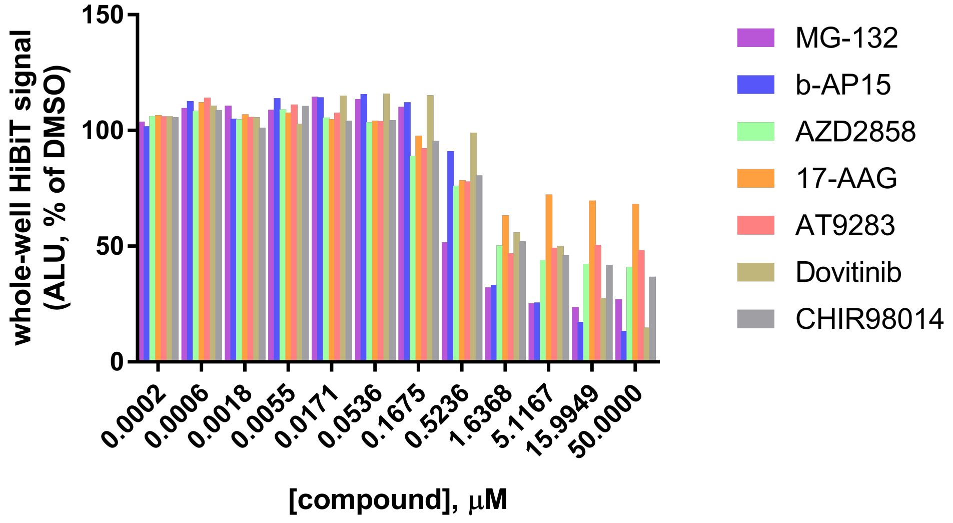

Supplemental Figure 4.

A

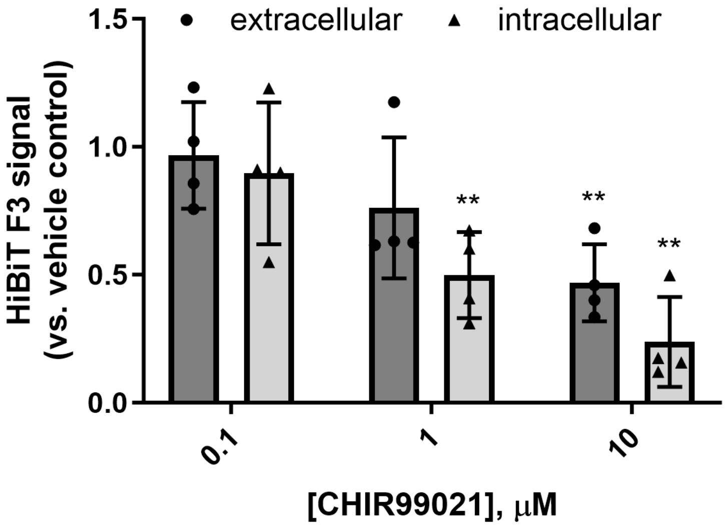

Supplemental Figure 5.

A

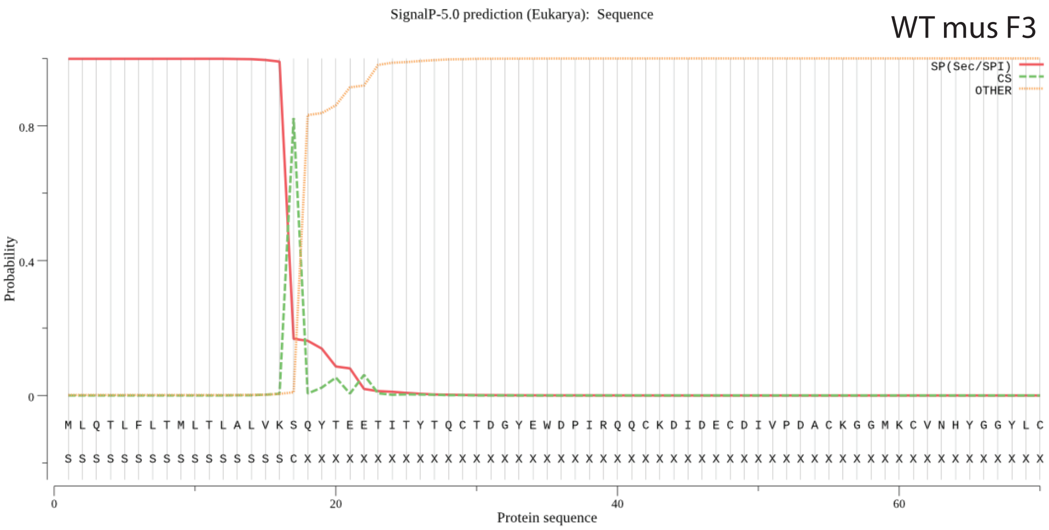


B

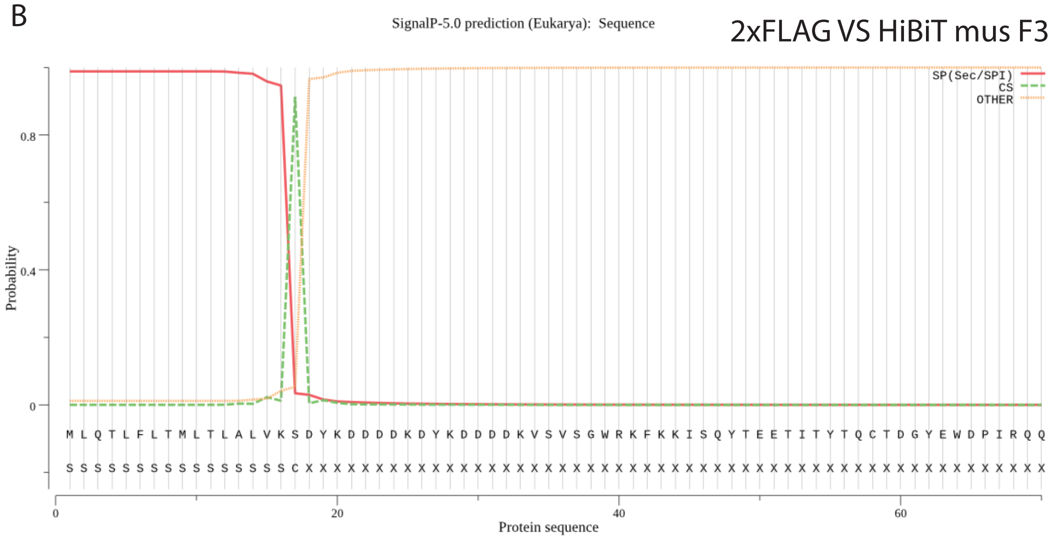

Supplemental Figure 6.

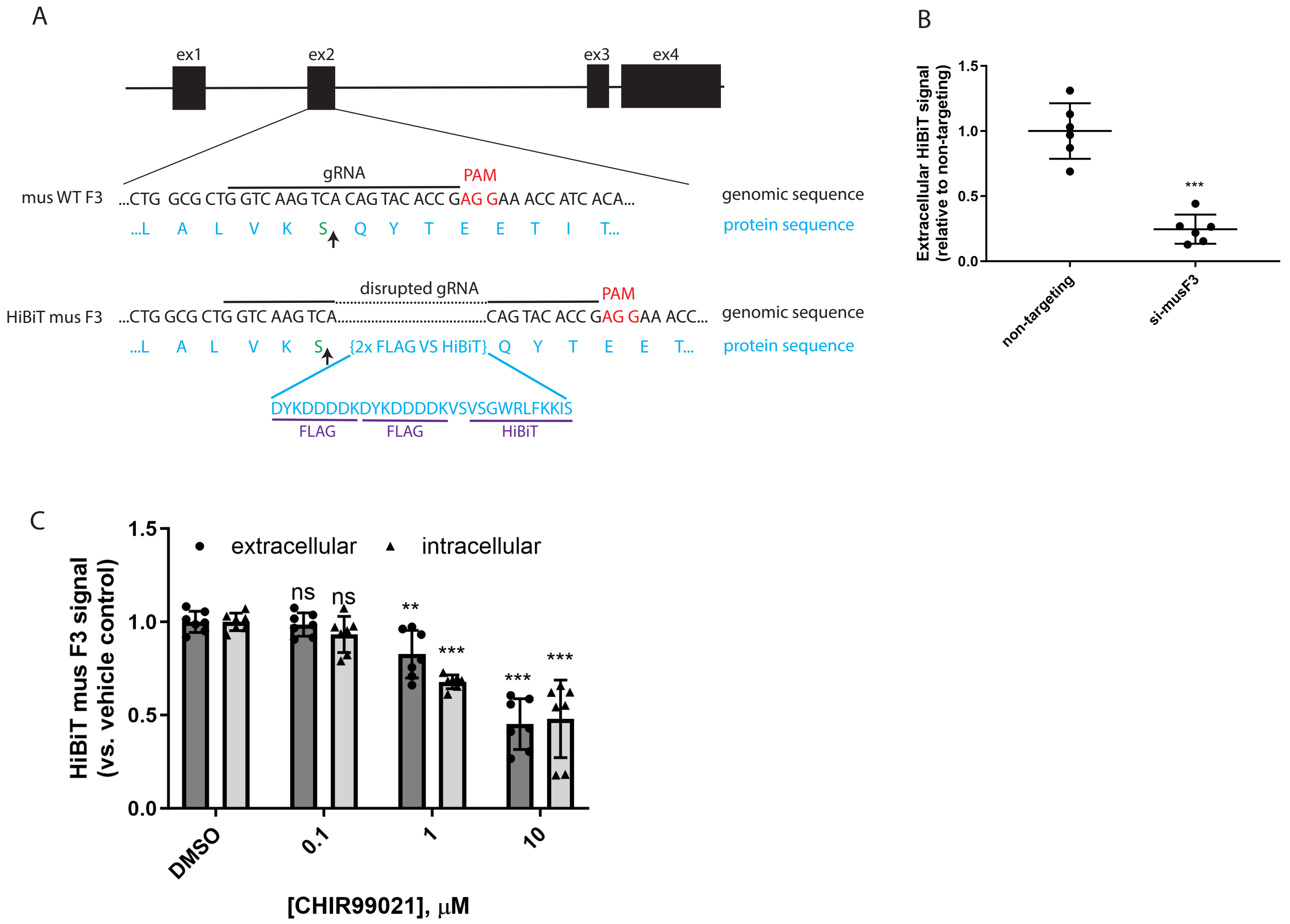

Supplemental Figure 7.

A

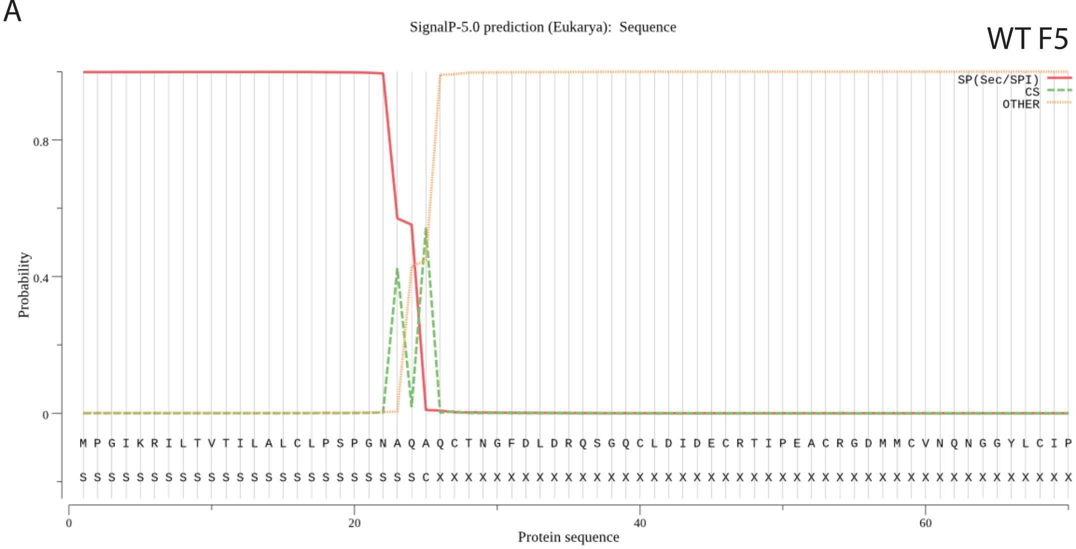


B

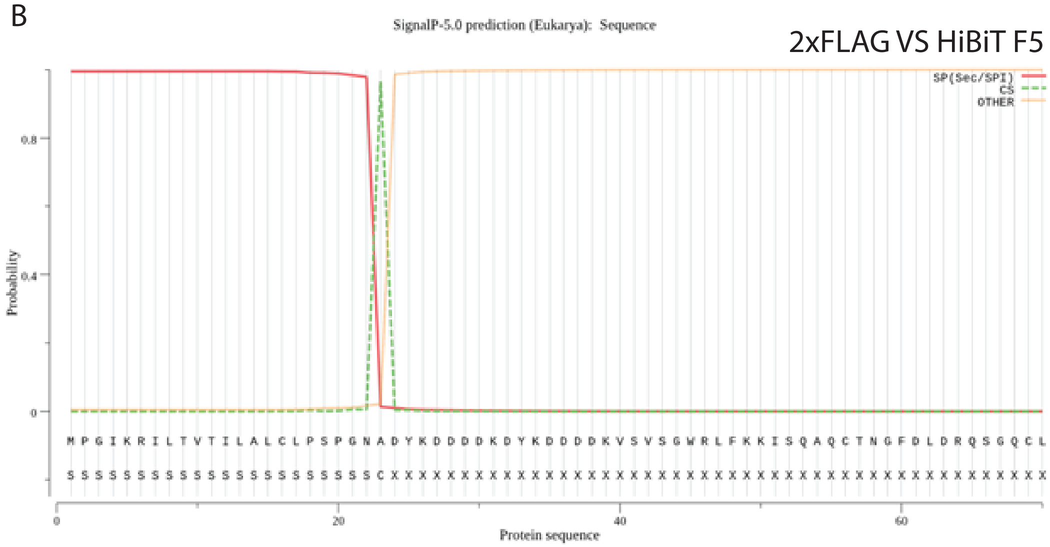

Supplemental Figure 8.

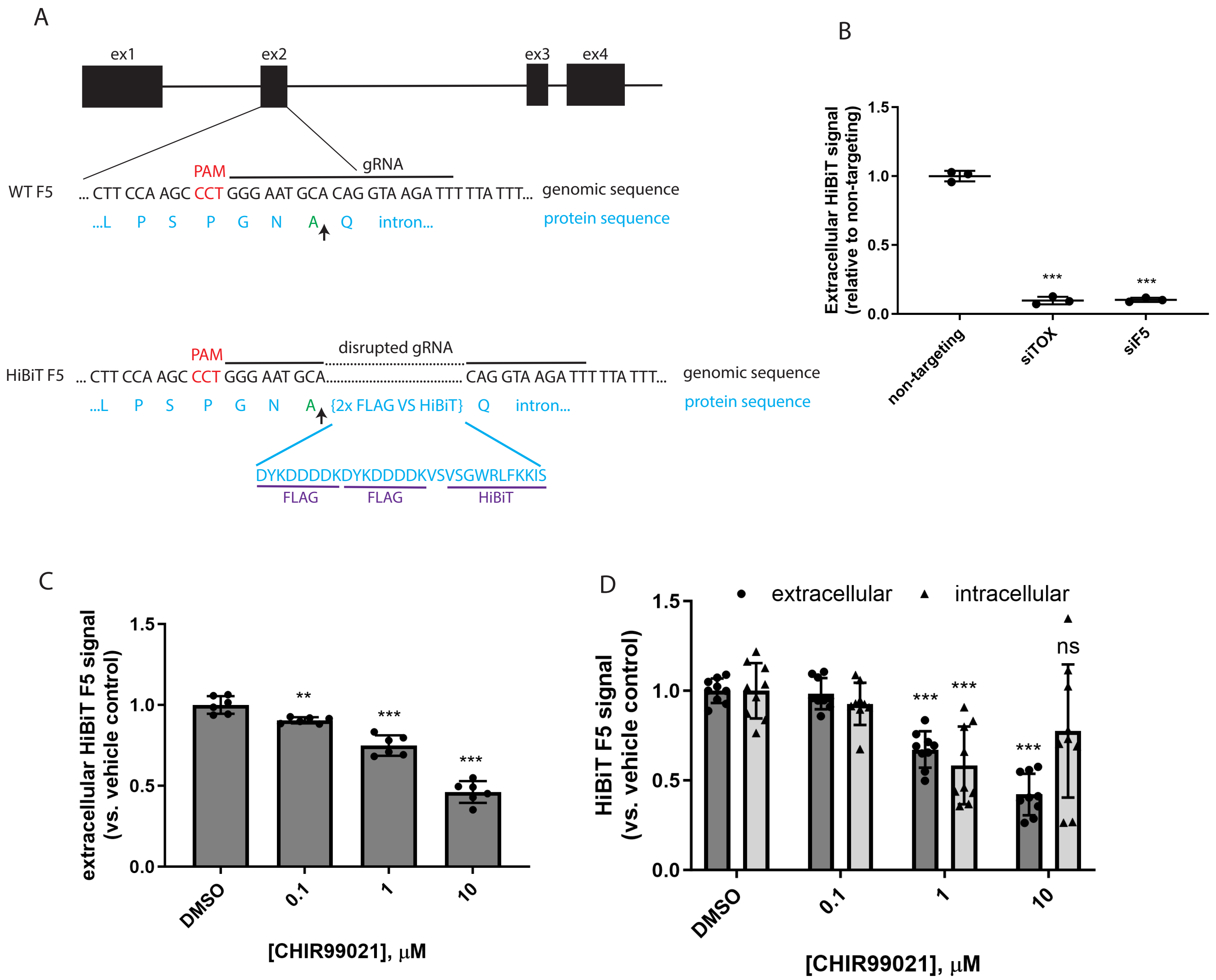

Supplemntal Figure 9.

A

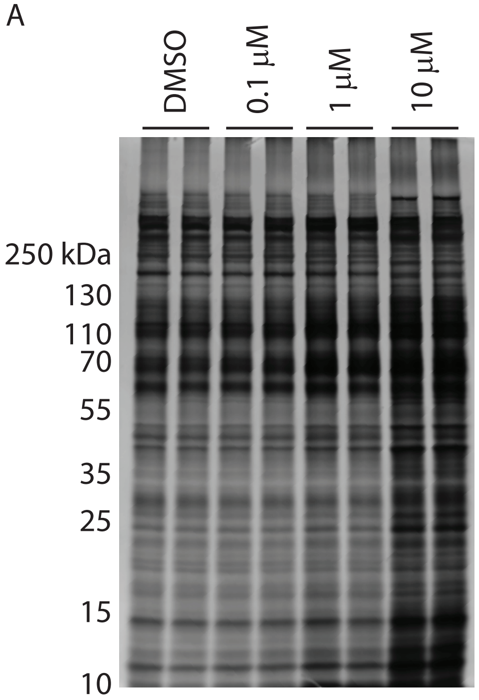

Supplemental Figure 10.

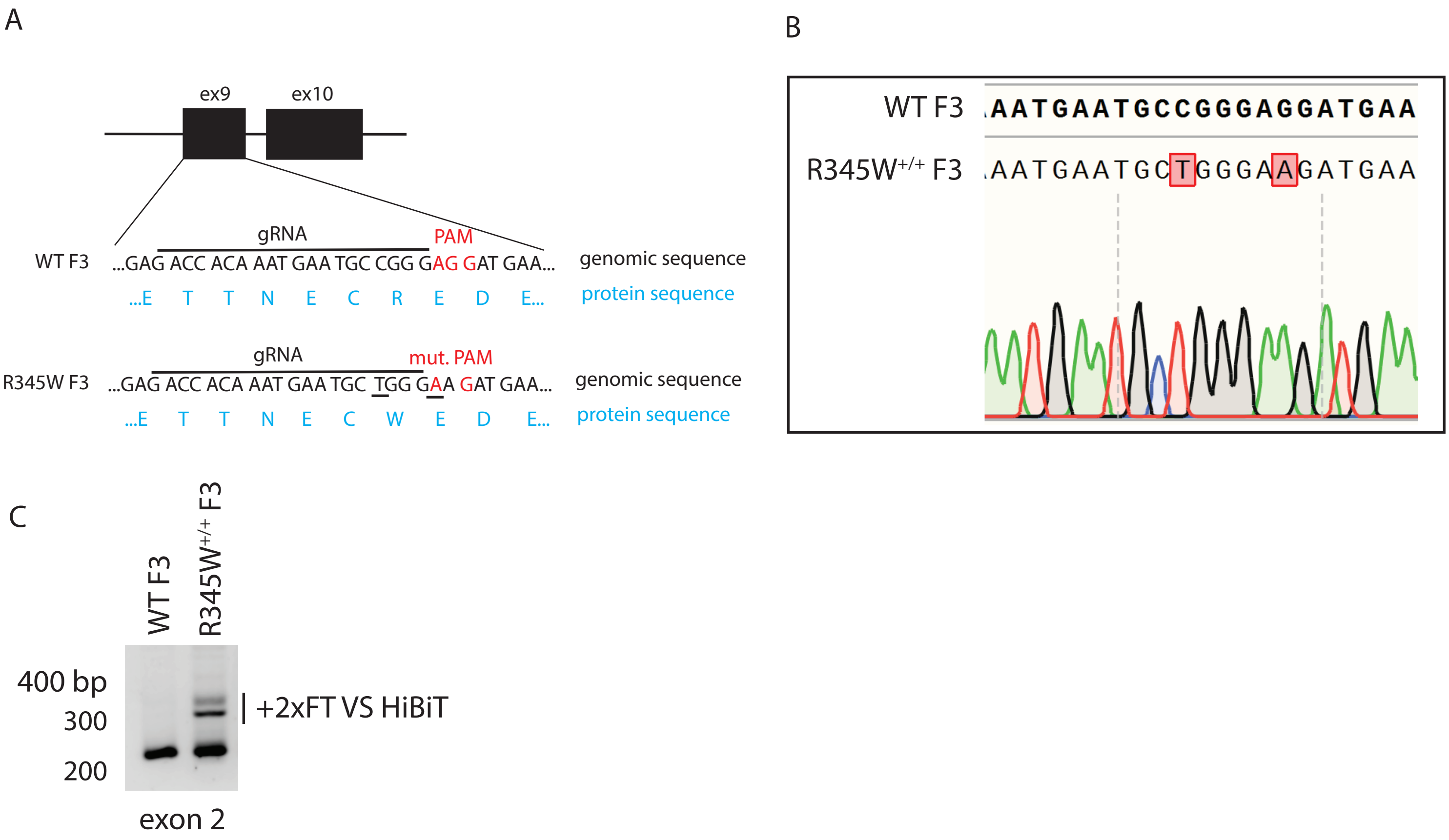

Supplemental Figure 11.

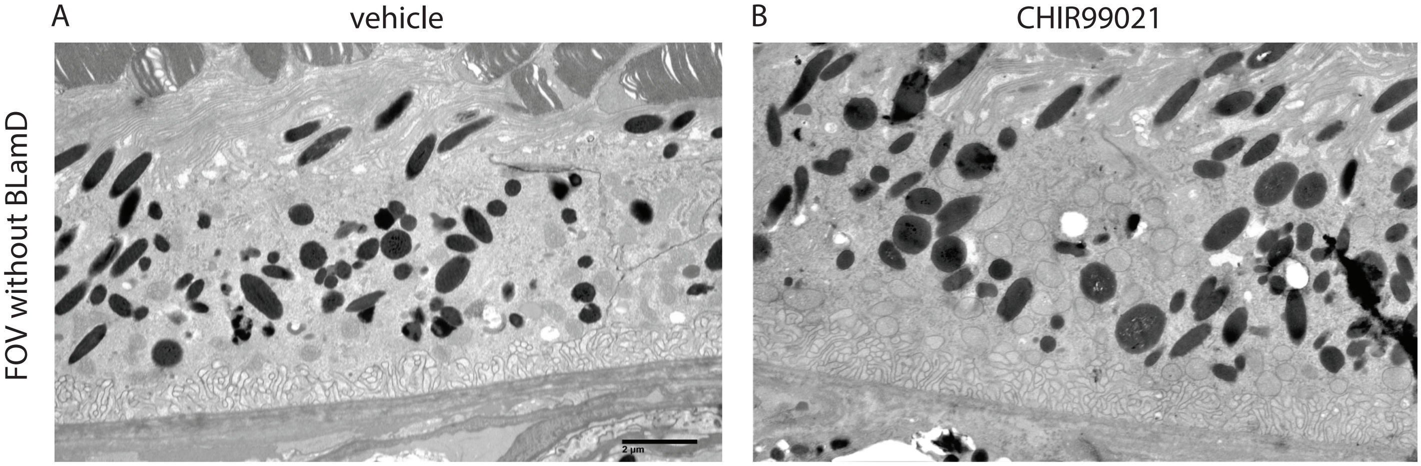

Supplemental Figure 12.

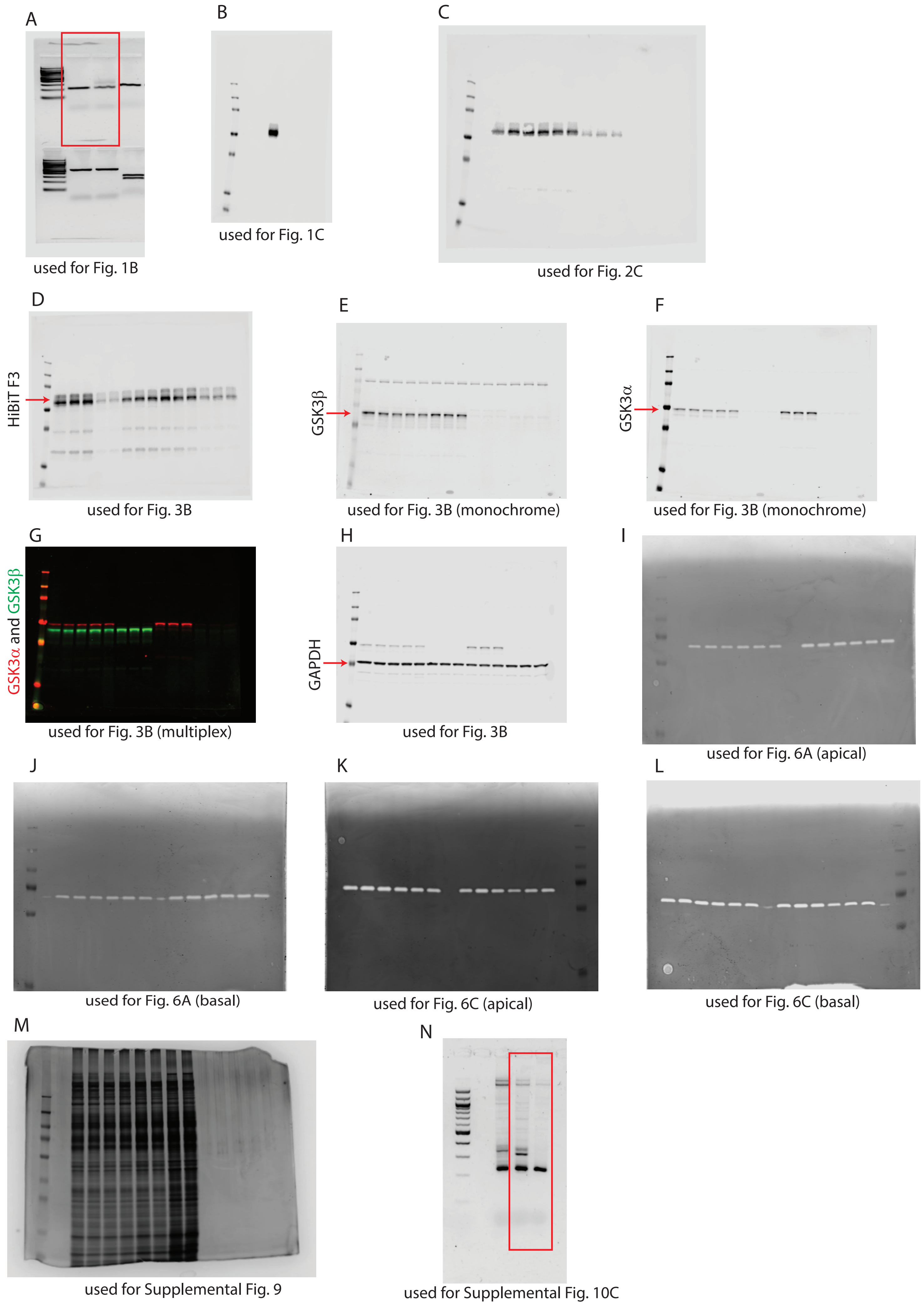

Supplemental Table 1:

| HiBiT edit | gRNA | PAM | ssODN |
|---|---|---|---|
| 2xFLAG VS HiBiT F3 (human) | CGTGTACGTGATGGTTTCTT | CGG | TGTTGAAAGCCCTTTTCCTAACTATGCTGACTCTGGCGCTGGTCAAGTCAG ATTACAAGGATGACGACGATAAGGATTACAAGGATGACGACGATAAGGT GAGCGTGAGCGGCTGGCGGCTGTTCAAGAAGATTAGCCAGGACACtGAA GAAACCATCACGTACACGGTAAGGGGTGATGGAATTTG |
| 2xFLAG VS HiBiT F3 (mouse) | GGTCAAGTCACAGTACACCG | AGG | TGTTGCAAACACTTTTCCTAACTATGCTGACTCTGGCGCTGGTCAAGTCAG ATTACAAGGATGACGACGATAAGGATTACAAGGATGACGACGATAAGGT GAGCGTGAGCGGCTGGCGGCTGTTCAAGAAGATTAGCCAGTACACCGAG GAAACCATCACATACACGGTAAGGCATGATGACATTTG |
| 2xFLAG VS HiBiT F5 (human) | AATCTTACCTGTGCATTCCC | AGG | ACAGGATACTCACTGTTACCATTCTGGCTCTCTGTCTTCCAAGCCCTGGGA ATGCAGATTACAAGGATGACGACGATAAGGATTACAAGGATGACGACGA TAAGGTGAGCGTGAGCGGCTGGCGGCTGTTCAAGAAGATTAGCCAGGTA AGATTTTTATTTTTAAAAAATATCTACAAGCGTATTAAAATCCGGGGTGA |

Supplemental Table 2:

| HiBiT edit | exon | forward primer | reverse primer |
|---|---|---|---|
| 2xFLAG VS HiBiT F3 (human) | 2 | TAAAGTTTTGTTACTTTCCCCGCAGCAGA | GTGTGTAAAGTCTTTTTTTGTCACAGAATCCCG |

Supplemental Table 3:

| Target | sense sequence |
|---|---|
| siTOX transfection control | proprietary |
| non-targeting siRNA control | proprietary |
| siF3#1 (human) | GCAAUGCACUGACGGAUAUtt |
| siF3#2 (human) | CAUCCGAUCUGAUAGGUCUtt |
| siF3 (mouse) | CAUUAAUACUUUUCGGAUUtt |
| siF5 (human) | GCAGCUCUGUGCGAAUGUUtt |
| siGSK3α (human) | GAAAGACGAGCUUUACCUAtt |
| siGSK3β (human) | CUCAAGAACUGUCAAGUAAtt |

### Supplemental Table 4 - Information on selected hit compounds based on primary, confirmatory, and counterscreening

| Structure, SMILES | common name | Target | Pathway | whole well HiBiT F3 (1.66 μM) | whole well HiBiT F3 (5 μM) | media only HiBiT F3 (1.66 μM, counterscreen) | media only HiBiT F3 (5 μM, counterscreen) | CTG (1.66 μM, counterscreen, 48 h Rx) | CTG ( counte , 48 h |
|---|---|---|---|---|---|---|---|---|---|
| 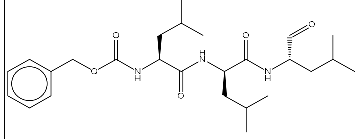 CC(C)C[C@H](NC(=O)[C@@H](CC(C)C)NC(=O)[C@H](CC(C)C)NC(=O)OCc1ccccc1)C=O | MG132 | Proteasome, Cysteine Protease | Proteases/Proteasome; Ubiquitination | -55.52 | -69.49 | -1.61 | -1.32 | -2.25 | -26 |
| 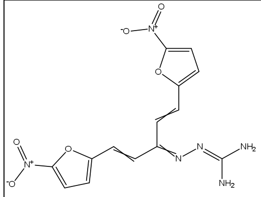 NC(N)=NN=C(C=CC1=CC=C(O1)[N+]([O-])=O)C=CC1=CC=C(O1)[N+]([O-])=O | | unknown | unknown | -47.01 | -69.62 | -5.07 | -7.72 | 7.42 | -40 |
| 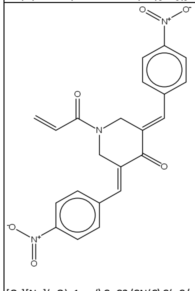 [O-][N+](=O)c1ccc(\C=C2/CN(C\C(=C/c3ccc(cc3)[N+]([O-])=O)C2=O)C(=O)C=C)cc1 | B-AP15 | DUB inhibitor | Ubiquitination | -38.36 | -63.01 | -5.24 | -5.82 | -5.54 | -7. |
| 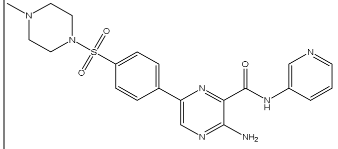 CN1CCN(CC1)S(=O)(=O)C1=CC=C(C=C1)C1=CN=C(N)C(=N1)C(=O)NC1=CN=CC=C1 | AZD2858 | GSK-3 inhibitor | PI3K/Akt/mTOR signaling; Stem Cells | -37.85 | -46.41 | -3.60 | -6.55 | 9.55 | 16. |
| 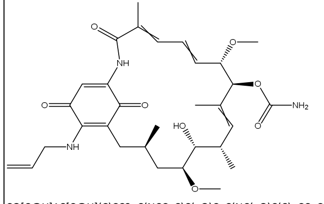 CO[C@H]1C[C@H](C)CC2=C(NCC=C)C(=O)C=C(NC(=O)C(C)=CC=C[C@H](OC)[C@@H](OC(N)=O)C(C)=C[C@H](C)[C@H]1O)C2= | Tanespimycin (17-AAG) | HSP (e.g. HSP90) | Cytoskeletal Signaling | -36.00 | -34.14 | -0.30 | -5.02 | -9.89 | -12 |
| 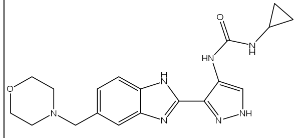 O=C(NC1CC1)NC1=CNN=C1C1=NC2=CC(CN3CCOCC3)=CC=C2N1 | AT9283 | Aurora Kinase inhibitor; Bcr-Abl inhibitor; JAK inhibitor | Angiogenesis; Cell Cycle/Checkpoint; Chromatin/Epigenetic; Cytoskeletal Signaling; JAK/STAT signaling; Stem Cells | -29.84 | -37.86 | -0.68 | -4.14 | 10.02 | 7. |
| 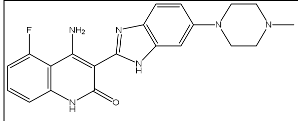 CN1CCN(CC1)C1=CC2=C(C=C1)N=C(N2)C1=C(N)C2=C(NC1=O)C=CC=C2F | Dovitinib (TKI-258, CHIR-258) | c-Kit,FGFR,FLT3,PDGFR,VEGFR | Angiogenesis | -26.64 | -44.03 | 1.94 | 2.43 | -5.14 | -21 |
| 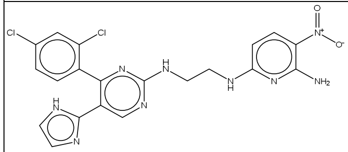 Nc1nc(NCCNc2ncc(-c3ncc[nH]3)c(n2)-c2ccc(Cl)cc2Cl)ccc1[N+]([O-])=O | CHIR98014 | GSK-3 inhibitor; S6 Kinase inhibitor | PI3K/Akt/mTOR signaling; Stem Cells | -26.41 | -37.93 | -4.02 | -3.26 | 32.09 | 37. |

Supplemental Table 5.

| accession # | gene name | description | fold change vs. DMSO |
|---|---|---|---|
| Q99470 | SDF2 | Stromal cell-derived factor 2 | 13.910 |
| O43660 | PLRG1 | Pleiotropic regulator 1 | 9.797 |
| Q9NRV9 | HEBP1 | Heme-binding protein 1 | 9.605 |
| P02452 | COL1A1 | Collagen alpha-1(I) | 8.871 |
| P48307 | TFPI2 | Tissue factor pathway inhibitor 2 | 4.974 |
| P36955 | SERPINF1 | Pigment epithelium-derived factor | 4.923 |
| P52803 | EFNA5 | Ephrin-A5 | 4.225 |
| Q14376 | GALE | UDP-glucose 4-epimerase | 3.823 |
| P81605 | DCD | Dermcidin | 3.809 |
| Q5D862 | FLG2 | Filaggrin-2 | 3.777 |
| P07311 | ACYP1 | Acylphosphatase-1 | 3.681 |
| P61081 | UBE2M | NEDD8-conjugating enzyme Ubc12 | 3.608 |
| O60245 | PCDH7 | Protocadherin-7 | 3.484 |
| P39748 | FEN1 | Flap endonuclease 1 | 3.471 |
| P49770 | EIF2B2 | Translation initiation factor eIF-2B subunit beta | 3.456 |
| P62266 | RPS23 | 40S ribosomal protein S23 | 3.443 |
| P20930 | FLG | Filaggrin | 3.417 |
| Q14112 | NID2 | Nidogen-2 | 3.395 |
| O14817 | TSPAN4 | Tetraspanin-4 | 3.372 |
| Q92520 | FAM3C | Protein FAM3C | 3.330 |
| P83916 | CBX1 | Chromobox protein homolog 1 | 3.301 |
| Q9UHJ6 | SHPK | Sedoheptulokinase | 3.228 |
| P61769 | B2M | Beta-2-microglobulin | 3.169 |
| Q6FI81 | CIAPIN1 | Anamorsin | 3.092 |
| O43488 | AKR7A2 | Aflatoxin B1 aldehyde reductase member 2 | 3.060 |
| Q9BXJ0 | C1QTNF5 | Complement C1q tumor necrosis factor-related protein 5 | 3.031 |
| Q9NRX4 | PHPT1 | 14 kDa phosphohistidine phosphatase | 3.014 |
| P15104 | GLUL | Glutamine synthetase | 2.966 |
| P21926 | CD9 | CD9 antigen | 2.966 |
| Q9UJU6 | DBNL | Drebrin-like protein | 2.962 |
| O75663 | TIPRL | TIP41-like protein | 2.873 |
| P10909 | CLU | Clusterin | 2.861 |
| P53611 | RABGGTB | Geranylgeranyl transferase type-2 subunit beta | 2.844 |
| P61204 | ARF3 | ADP-ribosylation factor 3 | 2.834 |
| P53990 | IST1 | IST1 homolog | 2.804 |
| P01009 | SERPINA1 | Alpha-1-antitrypsin | 2.801 |
| P27986 | PIK3R1 | Phosphatidylinositol 3-kinase regulatory subunit alpha | 2.793 |
| P52758 | RIDA | 2-iminobutanoate/2-iminopropanoate deaminase | 2.774 |
| P45877 | PPIC | Peptidyl-prolyl cis-trans isomerase C | 2.748 |
| P33908 | MAN1A1 | Mannosyl-oligosaccharide 1,2-alpha-mannosidase IA | 2.728 |
| O75144 | ICOSLG | ICOS ligand | 2.727 |
| P17900 | GM2A | Ganglioside GM2 activator | 2.710 |
| Q13045 | FLII | Protein flightless-1 homolog | 2.706 |
| Q14956 | GPNMB | Transmembrane glycoprotein NMB | 2.684 |
| O15400 | STX7 | Syntaxin-7 | 2.684 |
| Q8NBJ4 | GOLM1 | Golgi membrane protein 1 | 2.665 |
| Q969H8 | MYDGF | Myeloid-derived growth factor | 2.653 |
| Q9UII2 | ATP5IF1 | ATPase inhibitor, mitochondrial | 2.645 |
| P55196 | AFDN | Afadin | 2.638 |
| Q92824 | PCSK5 | Proprotein convertase subtilisin/kexin type 5 | 2.622 |

Supplemental Table 6.

| GO cellular component complete | # of genes in H. sapiens reference list | # of genes in dataset | expected frequency | fold enrichment | raw p value | FDR |
|---|---|---|---|---|---|---|
| keratohyalin granule (GO:0036457) | 4 | 2 | 0.03 | 66.85 | 8.06E-04 | 2.94E-02 |
| retromer complex (GO:0030904) | 12 | 3 | 0.09 | 33.42 | 1.71E-04 | 9.71E-03 |
| platelet alpha granule membrane (GO:0031092) | 17 | 3 | 0.13 | 23.59 | 4.17E-04 | 1.74E-02 |
| cytosolic small ribosomal subunit (GO:0022627) | 45 | 5 | 0.34 | 14.85 | 3.42E-05 | 2.59E-03 |
| chaperone complex (GO:0101031) | 42 | 4 | 0.31 | 12.73 | 3.74E-04 | 1.70E-02 |
| tertiary granule lumen (GO:1904724) | 55 | 5 | 0.41 | 12.15 | 8.31E-05 | 5.47E-03 |
| lysosomal lumen (GO:0043202) | 98 | 7 | 0.73 | 9.55 | 1.34E-05 | 1.10E-03 |
| specific granule lumen (GO:0035580) | 61 | 4 | 0.46 | 8.77 | 1.39E-03 | 4.59E-02 |
| small ribosomal subunit (GO:0015935) | 78 | 5 | 0.58 | 8.57 | 3.85E-04 | 1.71E-02 |
| platelet alpha granule (GO:0031091) | 91 | 5 | 0.68 | 7.35 | 7.50E-04 | 2.79E-02 |

### Supplemental Table 7 - List of down regulated genes by RNAseq

| gene_id | CH_0323 | CH_0407 | CH_0603 | DMSO_0323 | DMSO_0407 | DMSO_0603 | CHIR | DMSO | log2FoldChange | pvalue | padj | gene_name | gene_chr | gene_start | gene_end | gene_strand | gene_length | gene_biotype | gene_description |
|---|---|---|---|---|---|---|---|---|---|---|---|---|---|---|---|---|---|---|---|
| ENSG00000109193 | 0 | 0 | 0 | 35.89291575 | 49.51437596 | 10.73089607 | 0 | 32.04606259 | -7.527989742 | 2.41E-07 | 3.71E-05 | SULT1E1 | 4 | 69841212 | 69860152 | - | 2091 | protein_coding | sulfotransferase family 1E member 1 [Source:HGNC Symbol;Acc:HGNC:11377] |
| ENSG00000101203 | 0 | 0 | 0 | 27.68882072 | 18.18895443 | 24.68106096 | 0 | 23.51961204 | -7.080422905 | 1.80E-06 | 0.000204984 | COL20A1 | 20 | 63293186 | 63334851 | + | 10974 | protein_coding | collagen type XX alpha 1 chain [Source:HGNC Symbol;Acc:HGNC:14670] |
| ENSG00000198959 | 40.78680295 | 66.11933289 | 103.386971 | 681.9653992 | 685.1172837 | 321.9268821 | 70.09770229 | 563.0031883 | -2.996849739 | 2.01E-18 | 2.11E-15 | TGM2 | 20 | 38127387 | 38166578 | - | 8036 | protein_coding | transglutaminase 2 [Source:HGNC Symbol;Acc:HGNC:11778] |
| ENSG00000188869 | 121.2870719 | 55.42355845 | 54.92432837 | 431.7405008 | 645.7078824 | 692.1427966 | 77.21165292 | 589.8637266 | -2.942125725 | 4.17E-21 | 5.97E-18 | TMC3 | 15 | 81331217 | 81374213 | - | 4848 | protein_coding | transmembrane channel like 3 [Source:HGNC Symbol;Acc:HGNC:22995] |
| ENSG00000159251 | 55.81351983 | 79.73213672 | 116.3103424 | 380.4649069 | 790.2090204 | 600.93018 | 83.95199966 | 590.5347024 | -2.807025293 | 2.40E-19 | 3.15E-16 | ACTC1 | 15 | 34788095 | 34795726 | - | 4602 | protein_coding | actin, alpha, cardiac muscle 1 [Source:HGNC Symbol;Acc:HGNC:143] |
| ENSG00000111186 | 4.293347679 | 5.834058784 | 8.884817824 | 48.19905829 | 37.38840634 | 40.77740507 | 6.337408096 | 42.12162323 | -2.71346216 | 5.71E-05 | 0.003320259 | WNT5B | 12 | 1529891 | 1647243 | + | 4305 | protein_coding | Wnt family member 5B [Source:HGNC Symbol;Acc:HGNC:16265] |
| ENSG00000130720 | 10.7333692 | 19.44686261 | 8.884817824 | 69.73480773 | 122.2701937 | 45.0697635 | 13.02168321 | 79.02492164 | -2.607611205 | 3.65E-06 | 0.000378607 | FIBCD1 | 9 | 130902438 | 130939286 | - | 3831 | protein_coding | fibrinogen C domain containing 1 [Source:HGNC Symbol;Acc:HGNC:25922] |
| ENSG00000156466 | 13.95337996 | 25.2809214 | 6.46168569 | 75.88787901 | 184.9210367 | 13.95016489 | 15.23199568 | 91.58636021 | -2.596109033 | 0.000259883 | 0.010823946 | GDF6 | 8 | 96142330 | 96160792 | - | 3701 | protein_coding | growth differentiation factor 6 [Source:HGNC Symbol;Acc:HGNC:4221] |
| ENSG00000179772 | 31.12677067 | 56.39590158 | 23.42361063 | 210.2299351 | 272.8343165 | 121.2591256 | 36.98209429 | 201.4411257 | -2.450598997 | 2.19E-09 | 5.30E-07 | FOXS1 | 20 | 31844301 | 31845619 | - | 1319 | protein_coding | forkhead box S1 [Source:HGNC Symbol;Acc:HGNC:3735] |
| ENSG00000172575 | 157.7805272 | 140.0174108 | 196.2737028 | 1041.920069 | 775.0515584 | 876.714209 | 164.690547 | 897.8952786 | -2.443875171 | 3.30E-29 | 1.73E-25 | RASGRP1 | 15 | 38488103 | 38565575 | - | 6097 | protein_coding | RAS guanyl releasing protein 1 [Source:HGNC Symbol;Acc:HGNC:9878] |
| ENSG00000259207 | 69.76689978 | 36.94903897 | 128.4260031 | 425.5874296 | 349.6321241 | 439.9667389 | 78.38064728 | 405.0620975 | -2.361751506 | 1.36E-11 | 5.22E-09 | ITGB3 | 17 | 47253846 | 47311816 | + | 4394 | protein_coding | integrin subunit beta 3 [Source:HGNC Symbol;Acc:HGNC:6156] |
| ENSG00000186994 | 10.7333692 | 6.806401915 | 7.269396402 | 36.91842762 | 54.5668633 | 32.19268821 | 8.269722505 | 41.22599305 | -2.326471352 | 0.000386502 | 0.01439377 | KANK3 | 19 | 8322584 | 8343262 | - | 3594 | protein_coding | KN motif and ankyrin repeat domains 3 [Source:HGNC Symbol;Acc:HGNC:24796] |
| ENSG00000173597 | 16.1000538 | 35.00435271 | 10.50023925 | 136.3930798 | 97.00775698 | 61.16610761 | 20.53488192 | 98.18898147 | -2.264741002 | 1.45E-05 | 0.001156829 | SULT1B1 | 4 | 69721162 | 69787961 | - | 7356 | protein_coding | sulfotransferase family 1B member 1 [Source:HGNC Symbol;Acc:HGNC:17845] |
| ENSG00000171385 | 89.08696434 | 47.64481341 | 110.6563674 | 396.873097 | 403.1884899 | 346.6079431 | 82.46271506 | 382.2231767 | -2.208572081 | 2.06E-13 | 9.55E-11 | KCND3 | 1 | 111770662 | 111989155 | - | 7863 | protein_coding | potassium voltage-gated channel subfamily D member 3 [Source:HGNC Symbol;Acc:HGNC:6239] |
| ENSG00000238271 | 10.7333692 | 18.47451948 | 8.077107113 | 48.19905829 | 72.75581773 | 46.14285311 | 12.42833193 | 55.69924304 | -2.171203285 | 0.000184593 | 0.008284583 | IFNWP19 | 9 | 21455484 | 21456049 | + | 566 | unprocessed_pseudogene | interferon omega 1 pseudogene 19 [Source:HGNC Symbol;Acc:HGNC:5451] |
| ENSG00000081052 | 2553.468532 | 1604.366166 | 1811.695125 | 9845.939545 | 9411.773422 | 7119.949543 | 1989.843274 | 8792.55417 | -2.14401394 | 3.24E-27 | 1.28E-23 | COL4A4 | 2 | 227002711 | 227164453 | - | 10076 | protein_coding | collagen type IV alpha 4 chain [Source:HGNC Symbol;Acc:HGNC:2206] |
| ENSG00000168398 | 16.1000538 | 18.47451948 | 7.269396402 | 73.83685525 | 39.40940127 | 69.75082446 | 13.94798989 | 60.999027 | -2.141118718 | 0.000185631 | 0.008307528 | BDKRB2 | 14 | 96204679 | 96244166 | + | 6845 | protein_coding | bradykinin receptor B2 [Source:HGNC Symbol;Acc:HGNC:1030] |
| ENSG00000166448 | 12.88004304 | 9.723431307 | 25.84674276 | 25.63779696 | 93.97626457 | 94.43188543 | 16.15007237 | 71.34864899 | -2.130761546 | 0.000475409 | 0.016849972 | TMEM130 | 7 | 98846488 | 98870771 | - | 4663 | protein_coding | transmembrane protein 130 [Source:HGNC Symbol;Acc:HGNC:25429] |
| ENSG00000113578 | 51.52017215 | 20.41920575 | 15.34650351 | 113.8318185 | 145.5116355 | 116.9667672 | 29.0952938 | 125.4367404 | -2.124047646 | 3.97E-06 | 0.000406559 | FGF1 | 5 | 142592178 | 142698070 | - | 5275 | protein_coding | fibroblast growth factor 1 [Source:HGNC Symbol;Acc:HGNC:3665] |
| ENSG00000165879 | 26.83342299 | 4.861715654 | 16.96192494 | 60.50520083 | 83.87128989 | 66.53155564 | 16.21902119 | 70.30268212 | -2.122518655 | 0.000216576 | 0.009450763 | FRAT1 | 10 | 97319267 | 97321915 | + | 2649 | protein_coding | FRAT1, WNT signaling pathway regulator [Source:HGNC Symbol;Acc:HGNC:3944] |
| ENSG00000146674 | 10175.234 | 11419.19773 | 7134.508713 | 42943.30991 | 52576.18329 | 28831.77156 | 9576.31348 | 41450.42159 | -2.113920173 | 8.20E-22 | 1.29E-18 | IGFBP3 | 7 | 45912245 | 45921874 | - | 4571 | protein_coding | insulin like growth factor binding protein 3 [Source:HGNC Symbol;Acc:HGNC:5472] |
| ENSG00000169891 | 47.22682447 | 33.05966644 | 58.96288192 | 237.9187558 | 200.0784988 | 163.1096203 | 46.41645761 | 200.3689583 | -2.10525471 | 6.53E-10 | 1.77E-07 | REPS2 | X | 16946691 | 17153280 | + | 11311 | protein_coding | RALBP1 associated Eps domain containing 2 [Source:HGNC Symbol;Acc:HGNC:9963] |
| ENSG00000120937 | 1270.830913 | 2004.971536 | 931.2904501 | 3894.894114 | 7738.389614 | 6373.079177 | 1402.3643 | 6002.120968 | -2.097734356 | 7.13E-07 | 9.94E-05 | NPPB | 1 | 11857464 | 11858931 | - | 694 | protein_coding | natriuretic peptide B [Source:HGNC Symbol;Acc:HGNC:7940] |
| ENSG00000236333 | 9.660032278 | 3.889372523 | 29.07758561 | 49.22457017 | 55.57736077 | 78.33554132 | 14.2089968 | 61.04582409 | -2.08164626 | 0.001331583 | 0.036369598 | TRHDE-AS1 | 12 | 72253508 | 72274907 | - | 8781 | processed_transcript | TRHDE antisense RNA 1 [Source:HGNC Symbol;Acc:HGNC:27471] |
| ENSG00000160963 | 16.1000538 | 10.69577444 | 9.692528536 | 54.35212956 | 62.65084305 | 34.33886743 | 12.16278559 | 50.44728001 | -2.062556551 | 0.000459204 | 0.016403251 | COL26A1 | 7 | 101362820 | 101559024 | + | 3156 | protein_coding | collagen type XXVI alpha 1 chain [Source:HGNC Symbol;Acc:HGNC:18038] |
| ENSG00000139946 | 69.76689978 | 56.39590158 | 33.11613916 | 336.3678961 | 206.1414836 | 104.0896919 | 53.09298018 | 215.5330239 | -2.028186736 | 1.34E-06 | 0.000162561 | PELI2 | 14 | 56117814 | 56301526 | + | 6278 | protein_coding | pellino E3 ubiquitin protein ligase family member 2 [Source:HGNC Symbol;Acc:HGNC:8828] |
| ENSG00000164849 | 13.95337996 | 13.61280383 | 17.76963565 | 55.37764144 | 90.94477217 | 38.63122586 | 15.11193981 | 61.65121315 | -2.024561623 | 0.000250924 | 0.010625811 | GPR146 | 7 | 1044576 | 1059261 | + | 3160 | protein_coding | G protein-coupled receptor 146 [Source:HGNC Symbol;Acc:HGNC:21718] |
| ENSG00000072657 | 68.69356286 | 53.47887219 | 128.4260031 | 244.0718271 | 259.6978494 | 515.0830114 | 83.53281272 | 339.6175626 | -2.017013697 | 4.04E-08 | 7.00E-06 | TRHDE | 12 | 72087266 | 72670757 | + | 11950 | protein_coding | thyrotropin releasing hormone degrading enzyme [Source:HGNC Symbol;Acc:HGNC:30748] |
| ENSG00000101605 | 8.586695358 | 13.61280383 | 12.11566067 | 34.86740387 | 52.54586836 | 51.50830114 | 11.43838662 | 46.30719112 | -2.012208742 | 0.000738219 | 0.023493267 | MYOM1 | 18 | 3066807 | 3220108 | - | 6574 | protein_coding | myomesin 1 [Source:HGNC Symbol;Acc:HGNC:7613] |
| ENSG00000170379 | 47.22682447 | 25.2809214 | 29.07758561 | 202.0258401 | 133.3856658 | 60.093018 | 33.86177716 | 131.8348413 | -1.966933613 | 2.78E-05 | 0.001960099 | TCAF2 | 7 | 143620952 | 143730409 | + | 6409 | protein_coding | TRPM8 channel associated factor 2 [Source:HGNC Symbol;Acc:HGNC:26878] |
| ENSG00000141052 | 45.08015063 | 14.58514696 | 59.77059264 | 83.06646215 | 189.9735241 | 193.1561293 | 39.81196341 | 155.3987052 | -1.959580871 | 3.86E-05 | 0.002503575 | MYOCD | 17 | 12665890 | 12768949 | + | 8875 | protein_coding | myocardin [Source:HGNC Symbol;Acc:HGNC:16067] |
| ENSG00000138623 | 252.2341761 | 240.1687533 | 206.7739421 | 854.2513948 | 964.014585 | 874.5680298 | 233.0589572 | 897.6113365 | -1.947439929 | 3.61E-24 | 8.12E-21 | SEMA7A | 15 | 74409289 | 74433958 | - | 3800 | protein_coding | semaphorin 7A (John Milton Hagen blood group) [Source:HGNC Symbol;Acc:HGNC:10741] |
| ENSG00000010310 | 12.88004304 | 4.861715654 | 26.65445347 | 38.96945138 | 70.7348228 | 63.31228682 | 14.79873739 | 57.672187 | -1.946877622 | 0.001890538 | 0.046900229 | GIPR | 19 | 45668244 | 45683724 | + | 4736 | protein_coding | gastric inhibitory polypeptide receptor [Source:HGNC Symbol;Acc:HGNC:4271] |
| ENSG00000233818 | 16.1000538 | 32.08732331 | 12.92337138 | 109.729771 | 75.78731014 | 48.28903232 | 20.3702495 | 77.93537115 | -1.940866785 | 0.000252985 | 0.01068437 | AP000695.2 | 21 | 36445731 | 36532408 | + | 1178 | antisense | novel transcript |
| ENSG00000176678 | 15.02671688 | 12.6404607 | 37.15469272 | 72.81134337 | 86.90278229 | 89.06643739 | 21.6072901 | 82.92685435 | -1.92369429 | 0.000146571 | 0.006975633 | FOXL1 | 16 | 86576368 | 86582160 | + | 3742 | protein_coding | forkhead box L1 [Source:HGNC Symbol;Acc:HGNC:3817] |
| ENSG00000186479 | 1028.256769 | 425.8862913 | 1159.872581 | 3808.751117 | 2687.923266 | 3225.707359 | 871.3385473 | 3240.793914 | -1.894898438 | 1.88E-05 | 0.001434154 | RGS7BP | 5 | 64506257 | 64612312 | + | 4379 | protein_coding | regulator of G protein signaling 7 binding protein [Source:HGNC Symbol;Acc:HGNC:23271] |
| ENSG00000149591 | 971.3699124 | 1516.855284 | 858.5964861 | 3300.097225 | 5703.247712 | 3397.401696 | 1115.607227 | 4133.582211 | -1.8898532 | 8.24E-15 | 4.06E-12 | TAGLN | 11 | 117199321 | 117204782 | + | 3917 | protein_coding | transgelin [Source:HGNC Symbol;Acc:HGNC:11553] |
| ENSG00000172061 | 7.513358438 | 14.58514696 | 31.50071774 | 42.04598702 | 86.90278229 | 63.31228682 | 17.86640771 | 64.08701871 | -1.82476002 | 0.001984651 | 0.048850328 | LRRC15 | 3 | 194355247 | 194369743 | - | 5940 | protein_coding | leucine rich repeat containing 15 [Source:HGNC Symbol;Acc:HGNC:20818] |
| ENSG00000235750 | 9.660032278 | 14.58514696 | 19.38505707 | 45.12252265 | 64.67183799 | 45.0697635 | 14.5434121 | 51.62137471 | -1.816521157 | 0.001321055 | 0.036192298 | KIAA0040 | 1 | 175156987 | 175192999 | - | 4783 | protein_coding | KIAA0040 [Source:HGNC Symbol;Acc:HGNC:28950] |
| ENSG00000198542 | 51.52017215 | 56.39590158 | 64.6168569 | 227.663637 | 272.8343165 | 107.3089607 | 57.51097688 | 202.6023047 | -1.814981423 | 1.06E-06 | 0.000133348 | ITGBL1 | 13 | 101452593 | 101720856 | + | 6174 | protein_coding | integrin subunit beta like 1 [Source:HGNC Symbol;Acc:HGNC:6164] |
| ENSG00000169031 | 1150.617178 | 1079.300875 | 641.3223048 | 3563.653778 | 3927.80366 | 2572.195788 | 957.0801193 | 3354.551075 | -1.810464455 | 1.59E-15 | 9.25E-13 | COL4A3 | 2 | 227164565 | 227314792 | + | 9124 | protein_coding | collagen type IV alpha 3 chain [Source:HGNC Symbol;Acc:HGNC:2204] |
| ENSG00000100027 | 147.047158 | 47.64481341 | 84.00191397 | 406.1027039 | 303.1492406 | 248.9567889 | 92.89796179 | 319.4029111 | -1.78584993 | 5.71E-07 | 8.03E-05 | YPEL1 | 22 | 21697544 | 21735834 | - | 4713 | protein_coding | yippee like 1 [Source:HGNC Symbol;Acc:HGNC:12845] |
| ENSG00000152822 | 27.90675991 | 37.9213821 | 29.07758561 | 114.8573304 | 147.5326304 | 63.31228682 | 31.63524254 | 108.5674159 | -1.78007471 | 4.81E-05 | 0.002904996 | GRM1 | 6 | 146027646 | 146437598 | + | 7272 | protein_coding | glutamate metabotropic receptor 1 [Source:HGNC Symbol;Acc:HGNC:4593] |
| ENSG00000106366 | 5799.239377 | 6297.866458 | 7290.39688 | 18092.08056 | 26781.21441 | 21594.85525 | 6462.500905 | 22156.05007 | -1.777435712 | 4.90E-24 | 9.65E-21 | SERPINE1 | 7 | 101127089 | 101139266 | + | 3190 | protein_coding | serpin family E member 1 [Source:HGNC Symbol;Acc:HGNC:8583] |
| ENSG00000213694 | 22.54007531 | 11.66811757 | 17.76963565 | 49.22457017 | 69.72432533 | 57.94683879 | 17.32594284 | 58.96524476 | -1.770262374 | 0.000775055 | 0.024370148 | S1PR3 | 9 | 88990863 | 89005010 | + | 13578 | protein_coding | sphingosine-1-phosphate receptor 3 [Source:HGNC Symbol;Acc:HGNC:3167] |
| ENSG00000135318 | 716.9890624 | 535.761065 | 921.5979216 | 2801.698452 | 2140.233638 | 2449.863573 | 724.782683 | 2463.931888 | -1.764367706 | 2.29E-17 | 1.90E-14 | NT5E | 6 | 85449584 | 85495791 | + | 4533 | protein_coding | 5'-nucleotidase ecto [Source:HGNC Symbol;Acc:HGNC:8021] |
| ENSG00000165072 | 847.9361666 | 1108.471169 | 828.7111898 | 2834.514832 | 4182.449022 | 2331.823716 | 928.3728418 | 3116.262524 | -1.747253279 | 1.19E-15 | 7.23E-13 | MAMDC2 | 9 | 70043581 | 70226970 | + | 3854 | protein_coding | MAM domain containing 2 [Source:HGNC Symbol;Acc:HGNC:23673] |
| ENSG00000126561 | 45.08015063 | 7.778745046 | 33.11613916 | 73.83685525 | 73.7663152 | 140.5747385 | 28.65834495 | 96.059303 | -1.746612643 | 0.001374506 | 0.037012981 | STAT5A | 17 | 42287547 | 42311943 | + | 5754 | protein_coding | signal transducer and activator of transcription 5A [Source:HGNC Symbol;Acc:HGNC:11366] |
| ENSG00000113083 | 9751.265916 | 8842.488431 | 6430.992683 | 32886.11492 | 30966.69492 | 18184.57648 | 8341.582343 | 27345.79544 | -1.713026163 | 6.12E-15 | 3.21E-12 | LOX | 5 | 122063195 | 122078360 | - | 5547 | protein_coding | lysyl oxidase [Source:HGNC Symbol;Acc:HGNC:6664] |
| ENSG00000005187 | 34.34678143 | 35.97669584 | 29.07758561 | 148.6992224 | 139.4486507 | 35.41195704 | 33.13368762 | 107.8532767 | -1.705443103 | 0.000734519 | 0.023422817 | ACSM3 | 16 | 20610243 | 20797581 | + | 7822 | protein_coding | acyl-CoA synthetase medium chain family member 3 [Source:HGNC Symbol;Acc:HGNC:10522] |
| ENSG00000118523 | 11214.22414 | 12467.38362 | 10999.40447 | 33537.31496 | 45229.86669 | 33349.47881 | 11560.33741 | 37372.22015 | -1.69279902 | 6.41E-25 | 1.68E-21 | CTGF | 6 | 131948176 | 131951373 | - | 2339 | protein_coding | connective tissue growth factor [Source:HGNC Symbol;Acc:HGNC:2500] |
| ENSG00000111859 | 668.688901 | 588.2675941 | 772.9791507 | 2302.274167 | 2068.488318 | 2160.129379 | 676.6452153 | 2176.963955 | -1.685014264 | 3.65E-23 | 6.39E-20 | NEDD9 | 6 | 11183298 | 11382348 | - | 10359 | protein_coding | neural precursor cell expressed, developmentally down-regulated 9 [Source:HGNC Symbol;Acc:HGNC:7733] |
| ENSG00000249456 | 31.12677067 | 13.61280383 | 42.8086677 | 56.40315331 | 115.1967114 | 109.4551399 | 29.1827474 | 93.68500155 | -1.676335011 | 0.000758411 | 0.023990451 | AL731577.2 | 10 | 124917143 | 124942881 | + | 555 | sense_overlapping | novel transcript, sense overlapping ZRANB1 |
| ENSG00000131738 | 1355.62453 | 1385.588961 | 2075.008817 | 3976.935065 | 6000.333968 | 5338.620796 | 1605.407436 | 5105.296609 | -1.66844345 | 8.76E-16 | 5.52E-13 | KRT33B | 17 | 41363494 | 41369800 | - | 1607 | protein_coding | keratin 33B [Source:HGNC Symbol;Acc:HGNC:6451] |
| ENSG00000165959 | 68.69356286 | 7.778745046 | 55.73203908 | 137.4185917 | 147.5326304 | 129.8438425 | 44.06811566 | 138.2650215 | -1.650321147 | 0.001429585 | 0.038234727 | CLMN | 14 | 95181940 | 95319906 | - | 15701 | protein_coding | calmin [Source:HGNC Symbol;Acc:HGNC:19972] |
| ENSG00000187689 | 595.7019904 | 1207.650168 | 656.6688083 | 2079.73809 | 3493.289749 | 2062.478225 | 820.006989 | 2545.168688 | -1.634169926 | 7.06E-10 | 1.86E-07 | AMTN | 4 | 70518540 | 70532742 | + | 1037 | protein_coding | amelotin [Source:HGNC Symbol;Acc:HGNC:33188] |
| ENSG00000169991 | 642.9288149 | 445.3331539 | 530.6659373 | 1388.543083 | 1914.892703 | 1716.943371 | 539.6426354 | 1673.459719 | -1.633440365 | 2.04E-16 | 1.53E-13 | IFFO2 | 1 | 18904281 | 18956686 | - | 6188 | protein_coding | intermediate filament family orphan 2 [Source:HGNC Symbol;Acc:HGNC:27006] |
| ENSG00000140015 | 33.27344451 | 47.64481341 | 76.73251757 | 175.3625312 | 148.5431279 | 164.1827099 | 52.5502585 | 162.696123 | -1.620348178 | 9.98E-06 | 0.000850083 | KCNH5 | 14 | 62699454 | 63102037 | - | 12320 | protein_coding | potassium voltage-gated channel subfamily H member 5 [Source:HGNC Symbol;Acc:HGNC:6254] |
| ENSG00000260604 | 500.1750046 | 397.6883405 | 518.5502767 | 1627.487351 | 1385.392029 | 1279.122812 | 472.1378739 | 1430.667397 | -1.599063337 | 1.69E-17 | 1.48E-14 | AL590004.3 | 6 | 3904920 | 3911979 | - | 7060 | lincRNA | novel transcript |
| ENSG00000079841 | 129.8737673 | 213.9154888 | 334.3922345 | 502.5008204 | 776.0620558 | 772.6245171 | 226.0604968 | 683.7291311 | -1.592726917 | 9.63E-08 | 1.60E-05 | RIMS1 | 6 | 71886703 | 72403143 | + | 12788 | protein_coding | regulating synaptic membrane exocytosis 1 [Source:HGNC Symbol;Acc:HGNC:17282] |
| ENSG00000261051 | 31.12677067 | 27.22560766 | 19.38505707 | 102.5511878 | 68.71382756 | 57.94683879 | 25.91247847 | 76.4039515 | -1.56720895 | 0.000876285 | 0.026641313 | AC107021.2 | 3 | 146059585 | 146061679 | - | 2095 | sense_overlapping | novel transcript, overlapping to PLOD2 |
| ENSG00000106688 | 170.6605702 | 146.8238127 | 124.3874495 | 467.6334166 | 458.7658507 | 374.5082729 | 147.2906108 | 433.6358467 | -1.56121042 | 5.75E-11 | 1.93E-08 | SLC1A1 | 9 | 4490444 | 4587469 | + | 3998 | protein_coding | solute carrier family 1 member 1 [Source:HGNC Symbol;Acc:HGNC:10939] |
| ENSG00000164694 | 98.74699661 | 97.23431307 | 43.61637841 | 243.0463152 | 350.6426216 | 106.2358711 | 79.86589603 | 233.3082693 | -1.55181048 | 0.000195497 | 0.008675108 | FNDC1 | 6 | 159169397 | 159272109 | + | 6854 | protein_coding | fibronectin type III domain containing 1 [Source:HGNC Symbol;Acc:HGNC:21184] |
| ENSG00000139211 | 157.7805272 | 144.8791265 | 270.5830883 | 543.5212956 | 682.0857913 | 446.4052766 | 191.080914 | 557.3374545 | -1.540690469 | 2.00E-08 | 3.80E-06 | AMIGO2 | 12 | 47075707 | 47079951 | - | 3956 | protein_coding | adhesion molecule with Ig like domain 2 [Source:HGNC Symbol;Acc:HGNC:24073] |
| ENSG00000146005 | 24.68674915 | 22.36389201 | 30.69300703 | 92.29606906 | 58.60885318 | 75.1162725 | 25.9145494 | 75.34039825 | -1.535424291 | 0.00078188 | 0.024414153 | PSD2 | 5 | 139795821 | 139844466 | + | 4513 | protein_coding | pleckstrin and Sec7 domain containing 2 [Source:HGNC Symbol;Acc:HGNC:19092] |
| ENSG00000206538 | 6245.747536 | 4304.56304 | 5523.933555 | 17357.81405 | 14030.75735 | 15154.17143 | 5358.081377 | 15514.24761 | -1.53385329 | 6.89E-19 | 8.35E-16 | VGLL3 | 3 | 86876388 | 86991119 | - | 12714 | protein_coding | vestigial like family member 3 [Source:HGNC Symbol;Acc:HGNC:24327] |
| ENSG00000166106 | 718.0623993 | 483.254536 | 555.7049694 | 1559.803567 | 1844.15788 | 1631.096203 | 585.6739682 | 1678.35255 | -1.519859595 | 1.89E-15 | 1.06E-12 | ADAMTS15 | 11 | 130448974 | 130476641 | + | 5673 | protein_coding | ADAM metallopeptidase with thrombospondin type 1 motif 15 [Source:HGNC Symbol;Acc:HGNC:16305] |
| ENSG00000171401 | 85.86695358 | 57.36824471 | 80.77107113 | 172.2859956 | 285.9707836 | 178.1328748 | 74.66875647 | 212.1298847 | -1.506851334 | 4.33E-06 | 0.000431712 | KRT13 | 17 | 41500981 | 41505705 | - | 2913 | protein_coding | keratin 13 [Source:HGNC Symbol;Acc:HGNC:6415] |
| ENSG00000186352 | 68.69356286 | 135.1556952 | 105.0023925 | 352.7760862 | 275.8658089 | 245.73752 | 102.9505502 | 291.459805 | -1.499065883 | 7.35E-07 | 0.000101338 | ANKRD37 | 4 | 185396021 | 185400628 | + | 2580 | protein_coding | ankyrin repeat domain 37 [Source:HGNC Symbol;Acc:HGNC:29593] |
| ENSG00000011028 | 287.6542945 | 140.0174108 | 270.5830883 | 614.2816152 | 658.8443495 | 699.6544239 | 232.7515979 | 657.5934628 | -1.49821201 | 7.36E-09 | 1.61E-06 | MRC2 | 17 | 62627401 | 62693597 | + | 6860 | protein_coding | mannose receptor C type 2 [Source:HGNC Symbol;Acc:HGNC:16875] |
| ENSG00000082497 | 310.1943698 | 268.3667041 | 303.6992274 | 837.8432047 | 825.5764318 | 819.8404599 | 294.0867671 | 827.7533654 | -1.492966695 | 6.46E-16 | 4.24E-13 | SERTAD4 | 1 | 210232799 | 210246631 | + | 5467 | protein_coding | SERTA domain containing 4 [Source:HGNC Symbol;Acc:HGNC:25236] |
| ENSG00000118515 | 232.9141116 | 358.7946152 | 316.6225988 | 692.220518 | 835.6814065 | 963.6344672 | 302.7771085 | 830.5121306 | -1.454300028 | 6.93E-11 | 2.46E-08 | SGK1 | 6 | 134169246 | 134318112 | - | 9992 | protein_coding | serum/glucocorticoid regulated kinase 1 [Source:HGNC Symbol;Acc:HGNC:10810] |
| ENSG00000108821 | 1792.472656 | 2918.001735 | 1432.878802 | 4397.394935 | 8115.30517 | 4202.218902 | 2047.784398 | 5571.639669 | -1.44410746 | 0.001532744 | 0.040175245 | COL1A1 | 17 | 50183289 | 50201632 | - | 9819 | protein_coding | collagen type I alpha 1 chain [Source:HGNC Symbol;Acc:HGNC:2197] |
| ENSG00000049192 | 27.90675991 | 24.30857827 | 31.50071774 | 64.60724834 | 78.81880255 | 82.62789975 | 27.90535197 | 75.35131688 | -1.43039434 | 0.001154711 | 0.032598867 | ADAMTS6 | 5 | 65148736 | 65481920 | - | 8567 | protein_coding | ADAM metallopeptidase with thrombospondin type 1 motif 6 [Source:HGNC Symbol;Acc:HGNC:222] |
| ENSG00000171219 | 37.56679219 | 42.78309775 | 81.57878184 | 115.8828423 | 134.3961633 | 188.8637709 | 53.97622393 | 146.3809255 | -1.43010457 | 0.00027673 | 0.011411844 | CDC42BPG | 11 | 64823387 | 64844569 | - | 6112 | protein_coding | CDC42 binding protein kinase gamma [Source:HGNC Symbol;Acc:HGNC:29829] |
| ENSG00000103647 | 64.40021518 | 53.47887219 | 37.15469272 | 179.4645787 | 148.5431279 | 85.84716857 | 51.6779267 | 137.9516251 | -1.422864007 | 0.000306309 | 0.012154356 | CORO2B | 15 | 68578969 | 68727806 | + | 3998 | protein_coding | coronin 2B [Source:HGNC Symbol;Acc:HGNC:2256] |
| ENSG00000166482 | 90.16030126 | 78.75979359 | 49.27035339 | 149.7247343 | 262.7293418 | 163.1096203 | 72.73014941 | 191.8545655 | -1.405302866 | 6.61E-05 | 0.003734219 | MFAP4 | 17 | 19383442 | 19387240 | - | 2152 | protein_coding | microfibril associated protein 4 [Source:HGNC Symbol;Acc:HGNC:7035] |
| ENSG00000123843 | 31.12677067 | 90.42791116 | 46.84722126 | 143.571663 | 178.8580519 | 123.4053048 | 56.13396769 | 148.6116732 | -1.403785946 | 0.000536644 | 0.018407569 | C4BPB | 1 | 207088842 | 207099993 | + | 3219 | protein_coding | complement component 4 binding protein beta [Source:HGNC Symbol;Acc:HGNC:1328] |
| ENSG00000138356 | 297.3143268 | 244.0581258 | 441.8177591 | 950.6495113 | 847.8073761 | 804.8172054 | 327.7300706 | 867.7580309 | -1.402359644 | 1.59E-09 | 3.97E-07 | AOX1 | 2 | 200585868 | 200677064 | + | 5556 | protein_coding | aldehyde oxidase 1 [Source:HGNC Symbol;Acc:HGNC:553] |
| ENSG00000159263 | 79.42693206 | 48.61715654 | 72.69396402 | 232.7911964 | 191.994519 | 99.79733346 | 66.9126842 | 174.8610163 | -1.386744014 | 0.000229401 | 0.009846741 | SIM2 | 21 | 36699133 | 36749917 | + | 7652 | protein_coding | SIM bHLH transcription factor 2 [Source:HGNC Symbol;Acc:HGNC:10883] |
| ENSG00000166145 | 48.30016139 | 60.2852741 | 55.73203908 | 103.5766997 | 122.2701937 | 198.5215773 | 54.77249152 | 141.4561569 | -1.366827787 | 0.000219911 | 0.009569777 | SPINT1 | 15 | 40844018 | 40858207 | + | 3542 | protein_coding | serine peptidase inhibitor, Kunitz type 1 [Source:HGNC Symbol;Acc:HGNC:11246] |
| ENSG00000113389 | 1628.252107 | 1080.273218 | 1870.658007 | 4008.725933 | 3347.778113 | 4363.182343 | 1526.394444 | 3906.56213 | -1.35542502 | 3.81E-11 | 1.30E-08 | NPR3 | 5 | 32689070 | 32791724 | + | 9866 | protein_coding | natriuretic peptide receptor 3 [Source:HGNC Symbol;Acc:HGNC:7945] |
| ENSG00000115380 | 11189.53739 | 10246.55191 | 7366.321687 | 30725.36139 | 27095.47912 | 15691.78933 | 9600.803662 | 24504.20995 | -1.35188662 | 2.75E-09 | 6.57E-07 | EFEMP1 | 2 | 55865967 | 55924139 | - | 3962 | protein_coding | EGF containing fibulin extracellular matrix protein 1 [Source:HGNC Symbol;Acc:HGNC:3218] |
| ENSG00000101333 | 5509.438409 | 3896.178925 | 5698.399068 | 15208.34116 | 11337.7816 | 11901.63683 | 5034.672134 | 12815.91986 | -1.347923124 | 3.86E-13 | 1.74E-10 | PLCB4 | 20 | 9068763 | 9481242 | + | 6900 | protein_coding | phospholipase C beta 4 [Source:HGNC Symbol;Acc:HGNC:9059] |
| ENSG00000079931 | 97.6736597 | 98.2066562 | 134.8876888 | 328.1638011 | 300.1177482 | 210.325563 | 110.2560016 | 279.5357041 | -1.339133599 | 3.01E-06 | 0.000316285 | MOXD1 | 6 | 132296055 | 132401545 | - | 5293 | protein_coding | monooxygenase DBH like 1 [Source:HGNC Symbol;Acc:HGNC:21063] |
| ENSG00000119681 | 4977.063297 | 3911.736415 | 4549.026726 | 10741.21141 | 12504.90617 | 10745.91933 | 4479.275479 | 11330.67897 | -1.338959653 | 8.70E-18 | 8.56E-15 | LTBP2 | 14 | 74498170 | 74612378 | - | 9091 | protein_coding | latent transforming growth factor beta binding protein 2 [Source:HGNC Symbol;Acc:HGNC:6715] |
| ENSG00000146094 | 111.6270397 | 90.42791116 | 79.96336042 | 280.9902547 | 291.0232709 | 131.9900217 | 94.00610374 | 234.6678491 | -1.323013241 | 9.85E-05 | 0.005135705 | DOK3 | 5 | 177501907 | 177511274 | + | 4823 | protein_coding | docking protein 3 [Source:HGNC Symbol;Acc:HGNC:24583] |
| ENSG00000243753 | 63.32687826 | 72.9257348 | 35.5392713 | 152.8012699 | 199.0680013 | 74.04318289 | 57.26396145 | 141.970818 | -1.315299593 | 0.001966987 | 0.048491303 | HLA-L | 6 | 30259584 | 30293014 | + | 5505 | transcribed_unprocessed_pseudogene | major histocompatibility complex, class I, L (pseudogene) [Source:HGNC Symbol;Acc:HGNC:4970] |
| ENSG00000027075 | 75.13358438 | 67.09167602 | 75.92480686 | 186.6431619 | 203.1099912 | 152.3787242 | 72.71668909 | 180.7106258 | -1.31332388 | 1.52E-05 | 0.001201646 | PRKCH | 14 | 61187559 | 61550976 | + | 8833 | protein_coding | protein kinase C eta [Source:HGNC Symbol;Acc:HGNC:9403] |
| ENSG00000175274 | 991.7633138 | 788.570279 | 482.2032946 | 1778.237597 | 2268.566817 | 1513.056346 | 754.1789625 | 1853.28692 | -1.298541401 | 9.00E-08 | 1.51E-05 | TP53I11 | 11 | 44885903 | 44951306 | - | 7345 | protein_coding | tumor protein p53 inducible protein 11 [Source:HGNC Symbol;Acc:HGNC:16842] |
| ENSG00000162373 | 114.8470504 | 81.67682298 | 65.42456762 | 236.8932439 | 226.351433 | 177.0597852 | 87.316147 | 213.4348207 | -1.295314878 | 3.31E-05 | 0.002248249 | BEND5 | 1 | 48727523 | 48776969 | - | 2138 | protein_coding | BEN domain containing 5 [Source:HGNC Symbol;Acc:HGNC:25668] |
| ENSG00000074590 | 472.2682447 | 290.7305961 | 547.6278623 | 949.6239994 | 1140.851642 | 1113.867012 | 436.8755677 | 1068.114218 | -1.288802278 | 2.32E-08 | 4.30E-06 | NUAK1 | 12 | 106063340 | 106140033 | - | 7008 | protein_coding | NUAK family kinase 1 [Source:HGNC Symbol;Acc:HGNC:14311] |
| ENSG00000153162 | 2153.113861 | 1736.604831 | 1340.799781 | 4845.543626 | 4712.960193 | 3214.976463 | 1743.506158 | 4257.826761 | -1.288711946 | 7.19E-10 | 1.86E-07 | BMP6 | 6 | 7726797 | 7881422 | + | 2780 | protein_coding | bone morphogenetic protein 6 [Source:HGNC Symbol;Acc:HGNC:1073] |
| ENSG00000004799 | 59.03353058 | 75.8427642 | 63.80914619 | 236.8932439 | 123.2806912 | 124.4783944 | 66.22848032 | 161.5507765 | -1.286366043 | 0.000387476 | 0.014396019 | PDK4 | 7 | 95583499 | 95596491 | - | 4519 | protein_coding | pyruvate dehydrogenase kinase 4 [Source:HGNC Symbol;Acc:HGNC:8812] |
| ENSG00000175161 | 172.8072441 | 246.0028121 | 160.7344315 | 598.898937 | 434.5139115 | 366.9966456 | 193.1814959 | 466.8031647 | -1.273895983 | 1.38E-06 | 0.000164452 | CADM2 | 3 | 84958981 | 86074429 | + | 9990 | protein_coding | cell adhesion molecule 2 [Source:HGNC Symbol;Acc:HGNC:29849] |
| ENSG00000141738 | 201.7873409 | 151.6855284 | 114.694921 | 251.2504102 | 552.7421153 | 314.4152549 | 156.0559301 | 372.8025935 | -1.259635868 | 7.85E-05 | 0.004279865 | GRB7 | 17 | 39737927 | 39747291 | + | 3597 | protein_coding | growth factor receptor bound protein 7 [Source:HGNC Symbol;Acc:HGNC:4567] |
| ENSG00000198948 | 326.2944236 | 316.0115175 | 358.6235558 | 741.4450881 | 816.4819546 | 831.6444455 | 333.6431656 | 796.5238294 | -1.254542314 | 1.39E-11 | 5.22E-09 | MFAP3L | 4 | 169986597 | 170033031 | - | 8741 | protein_coding | microfibril associated protein 3 like [Source:HGNC Symbol;Acc:HGNC:29083] |
| ENSG00000177181 | 728.7957685 | 286.8412236 | 455.5488412 | 1592.619947 | 1076.179804 | 784.4285028 | 490.3952777 | 1151.076085 | -1.231968955 | 4.00E-05 | 0.002552983 | RIMKLA | 1 | 42380795 | 42422578 | + | 8981 | protein_coding | ribosomal modification protein rimK like family member A [Source:HGNC Symbol;Acc:HGNC:28725] |
| ENSG00000157680 | 38.64012911 | 47.64481341 | 84.00191397 | 102.5511878 | 162.6900924 | 136.2823801 | 56.7622855 | 133.8412201 | -1.229424144 | 0.001703509 | 0.043169597 | DGKI | 7 | 137381037 | 137847092 | - | 15549 | protein_coding | diacylglycerol kinase iota [Source:HGNC Symbol;Acc:HGNC:2855] |
| ENSG00000075618 | 825.3960913 | 1175.562845 | 1155.834028 | 1600.824042 | 2479.760788 | 3296.531273 | 1052.264321 | 2459.038701 | -1.224098213 | 2.73E-07 | 4.14E-05 | FSCN1 | 7 | 5592823 | 5606655 | + | 3692 | protein_coding | fascin actin-bundling protein 1 [Source:HGNC Symbol;Acc:HGNC:11148] |
| ENSG00000187498 | 11348.39125 | 8946.529146 | 9572.98735 | 20878.39633 | 27637.10576 | 21216.05462 | 9955.969249 | 23243.85224 | -1.2232707 | 4.40E-13 | 1.93E-10 | COL4A1 | 13 | 110148963 | 110307233 | - | 14738 | protein_coding | collagen type IV alpha 1 chain [Source:HGNC Symbol;Acc:HGNC:2202] |
| ENSG00000257671 | 136.3137888 | 129.3216364 | 127.6182924 | 306.6280517 | 364.7895861 | 245.73752 | 131.0845725 | 305.7183859 | -1.222734888 | 2.25E-06 | 0.000247918 | KRT7-AS | 12 | 52245048 | 52247448 | + | 1693 | antisense | KRT7 antisense RNA 1 [Source:HGNC Symbol;Acc:HGNC:52643] |
| ENSG00000144583 | 186.760624 | 188.6345674 | 250.3903205 | 316.8831704 | 587.0990292 | 546.20261 | 208.5951706 | 483.3949366 | -1.210547865 | 6.67E-06 | 0.000636951 | 4-Mar | 2 | 216257865 | 216372027 | - | 4447 | protein_coding | membrane associated ring-CH-type finger 4 [Source:HGNC Symbol;Acc:HGNC:29269] |
| ENSG00000115414 | 8505.121752 | 4717.80887 | 8098.107591 | 19262.18961 | 16107.32965 | 13969.48051 | 7107.012738 | 16446.33326 | -1.210450708 | 1.20E-08 | 2.42E-06 | FN1 | 2 | 215360440 | 215436172 | - | 17315 | protein_coding | fibronectin 1 [Source:HGNC Symbol;Acc:HGNC:3778] |
| ENSG00000058335 | 48.30016139 | 67.09167602 | 63.00143548 | 149.7247343 | 127.322681 | 136.2823801 | 59.4644243 | 137.7765985 | -1.209506455 | 0.000304785 | 0.012154356 | RASGRF1 | 15 | 78959947 | 79090773 | - | 9929 | protein_coding | Ras protein specific guanine nucleotide releasing factor 1 [Source:HGNC Symbol;Acc:HGNC:9875] |
| ENSG00000108846 | 64.40021518 | 59.31293097 | 90.46359967 | 123.0614254 | 244.5403874 | 125.551484 | 71.39224861 | 164.3844323 | -1.20024909 | 0.001223801 | 0.034155058 | ABCC3 | 17 | 50634777 | 50692252 | + | 9614 | protein_coding | ATP binding cassette subfamily C member 3 [Source:HGNC Symbol;Acc:HGNC:54] |
| ENSG00000185100 | 134.167115 | 224.6112632 | 180.1194886 | 410.2047514 | 447.6503786 | 377.7275417 | 179.6326223 | 411.8608906 | -1.195768963 | 1.62E-06 | 0.000187538 | ADSSL1 | 14 | 104724186 | 104747325 | + | 5775 | protein_coding | adenylosuccinate synthase like 1 [Source:HGNC Symbol;Acc:HGNC:20093] |
| ENSG00000079308 | 774.949256 | 317.9562037 | 460.3951054 | 1237.792837 | 1215.628455 | 1099.916847 | 517.7668551 | 1184.446046 | -1.195149851 | 4.66E-06 | 0.000458463 | TNS1 | 2 | 217799588 | 218033982 | - | 16281 | protein_coding | tensin 1 [Source:HGNC Symbol;Acc:HGNC:11973] |
| ENSG00000020181 | 84.79361666 | 64.17464663 | 102.5792603 | 179.4645787 | 190.9840216 | 200.6677565 | 83.84917454 | 190.3721189 | -1.179858211 | 0.000103935 | 0.005383214 | ADGRA2 | 8 | 37784191 | 37844896 | + | 7061 | protein_coding | adhesion G protein-coupled receptor A2 [Source:HGNC Symbol;Acc:HGNC:17849] |
| ENSG00000073464 | 169.5872333 | 142.9344402 | 206.7739421 | 414.3067989 | 403.1884899 | 360.558108 | 173.0985385 | 392.6844656 | -1.179647212 | 7.40E-07 | 0.000101338 | CLCN4 | X | 10156945 | 10237660 | + | 6789 | protein_coding | chloride voltage-gated channel 4 [Source:HGNC Symbol;Acc:HGNC:2022] |
| ENSG00000136490 | 49.37349831 | 81.67682298 | 51.69348552 | 95.3726047 | 181.8895443 | 135.2092905 | 60.91460227 | 137.4904798 | -1.175158083 | 0.001962981 | 0.048468412 | LIMD2 | 17 | 63695902 | 63701172 | - | 4745 | protein_coding | LIM domain containing 2 [Source:HGNC Symbol;Acc:HGNC:28142] |
| ENSG00000214517 | 3160.977229 | 3618.088789 | 3577.35074 | 7084.236056 | 8574.071021 | 7705.856469 | 3452.138919 | 7788.054515 | -1.173657304 | 6.43E-15 | 3.27E-12 | PPME1 | 11 | 74171099 | 74254703 | + | 7917 | protein_coding | protein phosphatase methylesterase 1 [Source:HGNC Symbol;Acc:HGNC:30178] |
| ENSG00000198053 | 2294.794334 | 1920.377683 | 2408.593341 | 4212.802797 | 5352.605091 | 5347.205512 | 2207.921786 | 4970.871133 | -1.170679729 | 2.78E-12 | 1.15E-09 | SIRPA | 20 | 1894167 | 1940592 | + | 5068 | protein_coding | signal regulatory protein alpha [Source:HGNC Symbol;Acc:HGNC:9662] |
| ENSG00000181274 | 65.4735521 | 102.0960287 | 82.38649255 | 175.3625312 | 222.3094431 | 162.0365307 | 83.31869113 | 186.5695017 | -1.161838892 | 0.000203564 | 0.008982479 | FRAT2 | 10 | 97332497 | 97334709 | - | 2213 | protein_coding | FRAT2, WNT signaling pathway regulator [Source:HGNC Symbol;Acc:HGNC:16048] |
| ENSG00000112139 | 5478.311638 | 4825.738958 | 4252.596895 | 9041.938232 | 12606.96642 | 10903.6635 | 4852.21583 | 10850.85605 | -1.161253741 | 1.28E-11 | 5.02E-09 | MDGA1 | 6 | 37630679 | 37699306 | - | 14187 | protein_coding | MAM domain containing glycosylphosphatidylinositol anchor 1 [Source:HGNC Symbol;Acc:HGNC:19267] |
| ENSG00000261068 | 46.15348755 | 70.98104854 | 52.50119623 | 139.4696155 | 94.98676204 | 144.867097 | 56.54524411 | 126.4411582 | -1.160036425 | 0.001516947 | 0.039893922 | AL512274.1 | 6 | 42092233 | 42094259 | - | 2027 | lincRNA | novel transcript |
| ENSG00000144642 | 829.6894389 | 567.8483883 | 702.7083188 | 1865.406107 | 1446.021877 | 1364.96998 | 700.0820487 | 1558.799322 | -1.155313054 | 8.00E-09 | 1.70E-06 | RBMS3 | 3 | 28574791 | 30010391 | + | 18339 | protein_coding | RNA binding motif single stranded interacting protein 3 [Source:HGNC Symbol;Acc:HGNC:13427] |
| ENSG00000235609 | 134.167115 | 125.4322639 | 151.041903 | 355.8526218 | 240.4983975 | 318.7076133 | 136.8804273 | 305.0195442 | -1.154477366 | 9.68E-06 | 0.000833403 | AF127577.4 | 21 | 14818843 | 15014430 | - | 5929 | lincRNA | novel transcript |
| ENSG00000125730 | 137.3871257 | 60.2852741 | 165.5806958 | 247.1483627 | 233.4249152 | 328.3654198 | 121.0843652 | 269.6462326 | -1.152602425 | 0.00076951 | 0.024244135 | C3 | 19 | 6677704 | 6730562 | - | 8328 | protein_coding | complement C3 [Source:HGNC Symbol;Acc:HGNC:1318] |
| ENSG00000105856 | 968.1499016 | 519.2312318 | 877.9815432 | 2053.074781 | 1694.604255 | 1505.544719 | 788.4542255 | 1751.074585 | -1.151254728 | 4.22E-07 | 6.16E-05 | HBP1 | 7 | 107168961 | 107202529 | + | 6351 | protein_coding | HMG-box transcription factor 1 [Source:HGNC Symbol;Acc:HGNC:23200] |
| ENSG00000133424 | 431.4814417 | 288.7859098 | 493.5112446 | 808.1033602 | 1053.94886 | 833.7906248 | 404.5928654 | 898.6142816 | -1.150386474 | 4.38E-07 | 6.33E-05 | LARGE1 | 22 | 33162226 | 33922841 | - | 10687 | protein_coding | LARGE xylosyl- and glucuronyltransferase 1 [Source:HGNC Symbol;Acc:HGNC:6511] |
| ENSG00000164176 | 1421.098082 | 1062.771042 | 1109.794517 | 3338.041164 | 2520.180687 | 2111.840347 | 1197.88788 | 2656.687399 | -1.149616446 | 1.02E-08 | 2.09E-06 | EDIL3 | 5 | 83940554 | 84384793 | - | 5738 | protein_coding | EGF like repeats and discoidin domains 3 [Source:HGNC Symbol;Acc:HGNC:3173] |
| ENSG00000054277 | 238.2807962 | 174.0494204 | 189.0043064 | 452.2507384 | 501.2067444 | 378.8006313 | 200.444841 | 444.086038 | -1.149551869 | 9.28E-07 | 0.000120873 | OPN3 | 1 | 241590102 | 241677376 | - | 3716 | protein_coding | opsin 3 [Source:HGNC Symbol;Acc:HGNC:14007] |
| ENSG00000185499 | 368.1545635 | 246.9751552 | 233.4283956 | 733.2409931 | 747.7681267 | 398.1162443 | 282.8527047 | 626.3751214 | -1.149115796 | 2.58E-05 | 0.001853962 | MUC1 | 1 | 155185824 | 155192916 | - | 4717 | protein_coding | mucin 1, cell surface associated [Source:HGNC Symbol;Acc:HGNC:7508] |
| ENSG00000213713 | 104.1136812 | 53.47887219 | 99.34841749 | 266.6330884 | 141.4696456 | 160.9634411 | 85.6469903 | 189.688725 | -1.146944454 | 0.001488031 | 0.039330447 | PIGCP1 | 11 | 33075566 | 33076460 | - | 895 | processed_pseudogene | phosphatidylinositol glycan anchor biosynthesis class C pseudogene 1 [Source:HGNC Symbol;Acc:HGNC:8961] |
| ENSG00000164197 | 447.5814955 | 346.1541545 | 421.6249913 | 1078.838496 | 815.4714571 | 793.0132197 | 405.1202138 | 895.774391 | -1.145001197 | 2.41E-08 | 4.41E-06 | RNF180 | 5 | 64165844 | 64372869 | + | 5434 | protein_coding | ring finger protein 180 [Source:HGNC Symbol;Acc:HGNC:27752] |
| ENSG00000001617 | 155.6338534 | 103.0683719 | 96.11757464 | 232.7911964 | 284.9602861 | 263.9800434 | 118.2732666 | 260.5771753 | -1.14388702 | 5.36E-05 | 0.003160929 | SEMA3F | 3 | 50155045 | 50189075 | + | 4826 | protein_coding | semaphorin 3F [Source:HGNC Symbol;Acc:HGNC:10728] |
| ENSG00000114796 | 1245.070827 | 1129.862718 | 1303.645088 | 3276.510452 | 2240.272888 | 2595.80376 | 1226.192878 | 2704.1957 | -1.140860585 | 2.36E-10 | 7.02E-08 | KLHL24 | 3 | 183635568 | 183684477 | + | 8650 | protein_coding | kelch like family member 24 [Source:HGNC Symbol;Acc:HGNC:25947] |
| ENSG00000133216 | 792.1226468 | 543.5398101 | 657.476519 | 1396.747178 | 1836.0739 | 1148.20588 | 664.3796586 | 1460.342319 | -1.136817909 | 1.27E-07 | 2.03E-05 | EPHB2 | 1 | 22710839 | 22921500 | + | 12516 | protein_coding | EPH receptor B2 [Source:HGNC Symbol;Acc:HGNC:3393] |
| ENSG00000152409 | 807.1493636 | 457.9736146 | 854.5579326 | 1755.676336 | 1342.951136 | 1535.591228 | 706.5603036 | 1544.739566 | -1.128101963 | 8.28E-07 | 0.000109619 | JMY | 5 | 79236189 | 79327215 | + | 9042 | protein_coding | junction mediating and regulatory protein, p53 cofactor [Source:HGNC Symbol;Acc:HGNC:28916] |
| ENSG00000165312 | 805.0026898 | 612.5761724 | 763.2866222 | 1122.935507 | 1473.305309 | 2151.544662 | 726.9551614 | 1582.595159 | -1.122251601 | 1.04E-06 | 0.000133022 | OTUD1 | 10 | 23439458 | 23442390 | + | 2933 | protein_coding | OTU deubiquitinase 1 [Source:HGNC Symbol;Acc:HGNC:27346] |
| ENSG00000108379 | 64.40021518 | 68.06401915 | 134.0799781 | 170.2349718 | 174.8160621 | 236.0797136 | 88.8480708 | 193.7102492 | -1.117466144 | 0.001369055 | 0.036929322 | WNT3 | 17 | 46762506 | 46833154 | - | 4334 | protein_coding | Wnt family member 3 [Source:HGNC Symbol;Acc:HGNC:12782] |
| ENSG00000170442 | 562.4285459 | 484.2268791 | 240.697792 | 820.4095028 | 1110.536718 | 841.302252 | 429.117739 | 924.0828242 | -1.10891946 | 4.00E-05 | 0.002552983 | KRT86 | 12 | 52249300 | 52309163 | + | 2394 | protein_coding | keratin 86 [Source:HGNC Symbol;Acc:HGNC:6463] |
| ENSG00000168453 | 370.3012373 | 260.587959 | 277.044774 | 351.7505743 | 735.6421571 | 868.1294922 | 302.6446568 | 651.8407412 | -1.107661965 | 0.000141977 | 0.006818793 | HR | 8 | 22114415 | 22133384 | - | 8784 | protein_coding | HR, lysine demethylase and nuclear receptor corepressor [Source:HGNC Symbol;Acc:HGNC:5172] |
| ENSG00000278948 | 62.25354134 | 139.0450677 | 185.7734636 | 244.0718271 | 320.3276975 | 271.4916706 | 129.0240242 | 278.6303984 | -1.105433193 | 0.001254424 | 0.034801769 | AL031587.5 | 22 | 37943050 | 37944898 | + | 1849 | TEC | TEC |
| ENSG00000133107 | 485.1482877 | 315.0391744 | 479.7801625 | 1128.063066 | 748.7786242 | 852.0331481 | 426.6558749 | 909.6249462 | -1.092002709 | 2.12E-06 | 0.00023634 | TRPC4 | 13 | 37636636 | 37870425 | - | 4205 | protein_coding | transient receptor potential cation channel subfamily C member 4 [Source:HGNC Symbol;Acc:HGNC:12336] |
| ENSG00000102452 | 183.5406133 | 179.8834792 | 195.4659921 | 481.9905829 | 410.2619722 | 298.3189108 | 186.2966949 | 396.8571553 | -1.09081472 | 1.05E-05 | 0.000887127 | NALCN | 13 | 101053776 | 101416492 | - | 9140 | protein_coding | sodium leak channel, non-selective [Source:HGNC Symbol;Acc:HGNC:19082] |
| ENSG00000154736 | 3217.864085 | 2346.263974 | 4669.375622 | 6398.16861 | 6503.561708 | 8684.514191 | 3411.167894 | 7195.414836 | -1.076532279 | 1.51E-06 | 0.000177623 | ADAMTS5 | 21 | 26917912 | 26966513 | - | 9056 | protein_coding | ADAM metallopeptidase with thrombospondin type 1 motif 5 [Source:HGNC Symbol;Acc:HGNC:221] |
| ENSG00000173801 | 2054.366864 | 1566.444784 | 1454.686991 | 3673.383549 | 3731.767151 | 3289.019646 | 1691.83288 | 3564.723449 | -1.075810326 | 6.16E-10 | 1.70E-07 | JUP | 17 | 41754604 | 41786931 | - | 4942 | protein_coding | junction plakoglobin [Source:HGNC Symbol;Acc:HGNC:6207] |
| ENSG00000151414 | 2188.533979 | 1212.511884 | 2090.355321 | 4880.41103 | 3475.100794 | 3216.049553 | 1830.467061 | 3857.187125 | -1.075333112 | 2.37E-06 | 0.000257643 | NEK7 | 1 | 198156963 | 198322420 | + | 5485 | protein_coding | NIMA related kinase 7 [Source:HGNC Symbol;Acc:HGNC:13386] |
| ENSG00000073910 | 225.4007531 | 156.547244 | 268.1599562 | 517.8834986 | 408.2409773 | 445.332187 | 216.7026511 | 457.152221 | -1.07522956 | 1.61E-05 | 0.001247406 | FRY | 13 | 31846713 | 32299122 | + | 18849 | protein_coding | FRY microtubule binding protein [Source:HGNC Symbol;Acc:HGNC:20367] |
| ENSG00000011201 | 1721.632419 | 1203.760796 | 1459.533255 | 3670.307013 | 3082.017279 | 2472.398455 | 1461.642157 | 3074.907582 | -1.073224863 | 3.40E-08 | 6.02E-06 | ANOS1 | X | 8528874 | 8732186 | - | 7131 | protein_coding | anosmin 1 [Source:HGNC Symbol;Acc:HGNC:6211] |
| ENSG00000091592 | 4218.214095 | 4394.018608 | 3739.700593 | 5959.249526 | 9073.25677 | 10909.02895 | 4117.311099 | 8647.178414 | -1.070574562 | 1.23E-07 | 1.99E-05 | NLRP1 | 17 | 5499427 | 5619424 | - | 12808 | protein_coding | NLR family pyrin domain containing 1 [Source:HGNC Symbol;Acc:HGNC:14374] |
| ENSG00000127329 | 630.0487719 | 427.8309775 | 932.0981608 | 1600.824042 | 1277.2688 | 1293.072977 | 663.3259701 | 1390.388606 | -1.066519387 | 1.34E-05 | 0.001090702 | PTPRB | 12 | 70515866 | 70637440 | - | 14542 | protein_coding | protein tyrosine phosphatase, receptor type B [Source:HGNC Symbol;Acc:HGNC:9665] |
| ENSG00000122707 | 934.8764571 | 640.7741231 | 830.3266112 | 1790.54374 | 1628.921919 | 1612.85368 | 801.9923971 | 1677.43978 | -1.064878553 | 7.21E-09 | 1.60E-06 | RECK | 9 | 36036433 | 36124451 | + | 6045 | protein_coding | reversion inducing cysteine rich protein with kazal motifs [Source:HGNC Symbol;Acc:HGNC:11345] |
| ENSG00000100302 | 75.13358438 | 116.6811757 | 139.7339531 | 223.5615895 | 298.0967532 | 172.7674268 | 110.5162377 | 231.4752565 | -1.062786445 | 0.000970127 | 0.028889233 | RASD2 | 22 | 35540868 | 35554001 | + | 3412 | protein_coding | RASD family member 2 [Source:HGNC Symbol;Acc:HGNC:18229] |
| ENSG00000112414 | 296.2409898 | 205.1644006 | 545.2047301 | 830.6646215 | 557.7946026 | 798.3786677 | 348.8700402 | 728.945964 | -1.060747876 | 0.000385316 | 0.014383616 | ADGRG6 | 6 | 142301854 | 142446266 | + | 8473 | protein_coding | adhesion G protein-coupled receptor G6 [Source:HGNC Symbol;Acc:HGNC:13841] |
| ENSG00000169432 | 592.4819797 | 286.8412236 | 406.2784878 | 1080.88952 | 882.16429 | 695.3620654 | 428.533897 | 886.1386251 | -1.049318107 | 5.11E-05 | 0.003036277 | SCN9A | 2 | 166195185 | 166376001 | - | 13741 | protein_coding | sodium voltage-gated channel alpha subunit 9 [Source:HGNC Symbol;Acc:HGNC:10597] |
| ENSG00000166046 | 317.7077282 | 281.0071648 | 357.0081344 | 832.7156453 | 514.3432115 | 627.7574202 | 318.5743425 | 658.2720923 | -1.046234932 | 4.82E-06 | 0.000471835 | TCP11L2 | 12 | 106301929 | 106347015 | + | 6558 | protein_coding | t-complex 11 like 2 [Source:HGNC Symbol;Acc:HGNC:28627] |
| ENSG00000173894 | 105.1870181 | 119.5982051 | 97.73299607 | 230.7401727 | 243.5298899 | 189.9368605 | 107.5060731 | 221.4023077 | -1.043631547 | 0.000149536 | 0.007073973 | CBX2 | 17 | 79778132 | 79787983 | + | 5687 | protein_coding | chromobox 2 [Source:HGNC Symbol;Acc:HGNC:1552] |
| ENSG00000132434 | 833.9827866 | 799.2660535 | 703.5160295 | 1528.012699 | 1887.609271 | 1391.79722 | 778.9216232 | 1602.473064 | -1.041462531 | 8.78E-09 | 1.82E-06 | LANCL2 | 7 | 55365448 | 55433742 | + | 5235 | protein_coding | LanC like 2 [Source:HGNC Symbol;Acc:HGNC:6509] |
| ENSG00000092295 | 954.1965216 | 1031.656062 | 838.4037183 | 1764.905943 | 2130.128664 | 1900.441694 | 941.4187672 | 1931.825434 | -1.037566978 | 7.12E-10 | 1.86E-07 | TGM1 | 14 | 24249114 | 24264432 | - | 3735 | protein_coding | transglutaminase 1 [Source:HGNC Symbol;Acc:HGNC:11777] |
| ENSG00000211455 | 1326.644433 | 1239.737492 | 1401.378084 | 3057.05091 | 2691.965256 | 2396.209093 | 1322.58667 | 2715.075086 | -1.03750029 | 1.68E-10 | 5.20E-08 | STK38L | 12 | 27243968 | 27325959 | + | 6921 | protein_coding | serine/threonine kinase 38 like [Source:HGNC Symbol;Acc:HGNC:17848] |
| ENSG00000095951 | 909.116371 | 721.478603 | 1267.298106 | 2189.46786 | 1753.213108 | 2002.385207 | 965.96436 | 1981.688725 | -1.035781778 | 8.50E-07 | 0.000111639 | HIVEP1 | 6 | 12008762 | 12164999 | + | 10979 | protein_coding | human immunodeficiency virus type I enhancer binding protein 1 [Source:HGNC Symbol;Acc:HGNC:4920] |
| ENSG00000120162 | 68.69356286 | 124.4599207 | 118.7334746 | 255.3524577 | 191.994519 | 192.0830397 | 103.9623194 | 213.1433388 | -1.032240917 | 0.00098687 | 0.029167296 | MOB3B | 9 | 27325209 | 27529781 | - | 7087 | protein_coding | MOB kinase activator 3B [Source:HGNC Symbol;Acc:HGNC:23825] |
| ENSG00000100311 | 191.0539717 | 197.3856555 | 181.73491 | 382.5159307 | 452.7028659 | 325.146151 | 190.0581791 | 386.7883158 | -1.025928969 | 1.05E-05 | 0.000887127 | PDGFB | 22 | 39223359 | 39244751 | - | 3761 | protein_coding | platelet derived growth factor subunit B [Source:HGNC Symbol;Acc:HGNC:8800] |
| ENSG00000135245 | 306.974359 | 265.4496747 | 186.5811743 | 563.0060213 | 621.4559432 | 354.1195704 | 253.001736 | 512.8605116 | -1.022102001 | 0.000135882 | 0.006579728 | HILPDA | 7 | 128455849 | 128458418 | + | 2070 | protein_coding | hypoxia inducible lipid droplet associated [Source:HGNC Symbol;Acc:HGNC:28859] |
| ENSG00000152779 | 374.594585 | 290.7305961 | 570.2437622 | 727.0879218 | 761.9150913 | 1016.215858 | 411.8563144 | 835.072957 | -1.017772277 | 5.17E-05 | 0.003061266 | SLC16A12 | 10 | 89430299 | 89556641 | - | 4835 | protein_coding | solute carrier family 16 member 12 [Source:HGNC Symbol;Acc:HGNC:23094] |
| ENSG00000091986 | 28882.42317 | 13070.23636 | 17390.81932 | 47428.89887 | 41577.92884 | 30748.3096 | 19781.15962 | 39918.3791 | -1.012967809 | 2.97E-05 | 0.002071007 | CCDC80 | 3 | 112596794 | 112649530 | - | 13071 | protein_coding | coiled-coil domain containing 80 [Source:HGNC Symbol;Acc:HGNC:30649] |
| ENSG00000122786 | 14457.84831 | 10156.124 | 11257.87189 | 26035.69557 | 25997.06837 | 20338.26732 | 11957.2814 | 24123.67709 | -1.012623704 | 8.60E-09 | 1.81E-06 | CALD1 | 7 | 134744252 | 134970728 | + | 9843 | protein_coding | caldesmon 1 [Source:HGNC Symbol;Acc:HGNC:1441] |
| ENSG00000187398 | 236.1341223 | 238.224067 | 369.1237951 | 543.5212956 | 647.7288773 | 511.8637426 | 281.1606615 | 567.7046385 | -1.011092187 | 2.09E-05 | 0.001550309 | LUZP2 | 11 | 24496970 | 25082631 | + | 8851 | protein_coding | leucine zipper protein 2 [Source:HGNC Symbol;Acc:HGNC:23206] |
| ENSG00000004399 | 2775.649274 | 2085.676015 | 1749.501401 | 4156.399643 | 5795.202982 | 3354.478112 | 2203.608897 | 4435.360246 | -1.009610767 | 4.27E-06 | 0.000428885 | PLXND1 | 3 | 129555175 | 129606818 | - | 9738 | protein_coding | plexin D1 [Source:HGNC Symbol;Acc:HGNC:9107] |
| ENSG00000138641 | 249.0141654 | 217.8048613 | 218.8896028 | 626.5877577 | 399.1465001 | 353.0464808 | 228.5695431 | 391.5935795 | -1.008626007 | 8.27E-05 | 0.004461722 | HERC3 | 4 | 88592423 | 88708542 | + | 6981 | protein_coding | HECT and RLD domain containing E3 ubiquitin protein ligase 3 [Source:HGNC Symbol;Acc:HGNC:4876] |
| ENSG00000130518 | 130.9471042 | 121.5428913 | 140.5416638 | 158.9543412 | 285.9707836 | 344.4617639 | 131.0105521 | 263.1289629 | -1.005091903 | 0.001163263 | 0.032722998 | IQCN | 19 | 18257097 | 18274509 | - | 9154 | protein_coding | IQ motif containing N [Source:HGNC Symbol;Acc:HGNC:29350] |
| ENSG00000163235 | 341.3211405 | 268.3667041 | 273.0062204 | 653.2510666 | 708.3587254 | 407.7740507 | 294.231355 | 589.7946142 | -1.004642258 | 4.09E-05 | 0.002591123 | TGFA | 2 | 70447280 | 70554193 | - | 4963 | protein_coding | transforming growth factor alpha [Source:HGNC Symbol;Acc:HGNC:11765] |

| gene_id | tf_family | CH_0323_count | CH_0407_count | CH_0603_count | DMSO_0323_count | DMSO_0407_count | DMSO_0603_count | CH_0323_fpkm | CH_0407_fpkm | CH_0603_fpkm | DMSO_0323_fpkm | DMSO_0407_fpkm | DMSO_0603_fpkm |
|---|---|---|---|---|---|---|---|---|---|---|---|---|---|
| ENSG00000109193 | - | 0 | 0 | 0 | 35 | 49 | 10 | 0 | 0 | 0 | 0.738979616 | 1.019424413 | 0.220932552 |
| ENSG00000101203 | - | 0 | 0 | 0 | 27 | 18 | 23 | 0 | 0 | 0 | 0.108621865 | 0.071354363 | 0.096822573 |
| ENSG00000198959 | - | 38 | 68 | 128 | 665 | 678 | 300 | 0.218503043 | 0.354214461 | 0.553864636 | 3.653424736 | 3.67031001 | 1.724626551 |
| ENSG00000188869 | - | 113 | 57 | 68 | 421 | 639 | 645 | 1.077034594 | 0.492163664 | 0.487730479 | 3.833874853 | 5.733914719 | 6.146258823 |
| ENSG00000159251 | - | 52 | 82 | 144 | 371 | 782 | 560 | 0.522120265 | 0.745872407 | 1.088051551 | 3.559145503 | 7.392190003 | 5.621538041 |
| ENSG00000111186 | - | 4 | 6 | 11 | 47 | 37 | 38 | 0.042933931 | 0.058341205 | 0.088849119 | 0.481995684 | 0.373888021 | 0.407778367 |
| ENSG00000130720 | - | 10 | 20 | 11 | 68 | 121 | 42 | 0.120615096 | 0.218532053 | 0.099842197 | 0.783637496 | 1.373998317 | 0.506466681 |
| ENSG00000156466 | - | 13 | 26 | 8 | 74 | 183 | 13 | 0.162307313 | 0.294070572 | 0.075163068 | 0.882736496 | 2.151022667 | 0.162269915 |
| ENSG00000179772 | Fork | 29 | 58 | 29 | 205 | 270 | 113 | 1.015937574 | 1.840689356 | 0.764516384 | 6.861633433 | 8.904959548 | 3.957741175 |
| ENSG00000172575 | - | 147 | 144 | 243 | 1016 | 767 | 817 | 1.114076364 | 0.988652343 | 1.385873764 | 7.356918764 | 5.472580408 | 6.190412691 |
| ENSG00000259207 | - | 65 | 38 | 159 | 415 | 346 | 410 | 0.683545022 | 0.362010233 | 1.258260799 | 4.16971615 | 3.425539885 | 4.310598221 |
| ENSG00000186994 | - | 10 | 7 | 9 | 36 | 54 | 30 | 0.128568847 | 0.081529968 | 0.087075912 | 0.442224577 | 0.653625022 | 0.385617667 |
| ENSG00000173597 | - | 15 | 36 | 13 | 133 | 96 | 57 | 0.094224395 | 0.204860431 | 0.061451887 | 0.798230592 | 0.567730851 | 0.357970202 |
| ENSG00000171385 | - | 83 | 49 | 137 | 387 | 399 | 323 | 0.487757142 | 0.260858569 | 0.60585108 | 2.17290699 | 2.207484193 | 1.89770188 |
| ENSG00000238271 | - | 10 | 19 | 10 | 47 | 72 | 43 | 0.816389461 | 1.40518813 | 0.61435184 | 3.666062582 | 5.53387121 | 3.509665815 |
| ENSG00000081052 | - | 2379 | 1650 | 2243 | 9601 | 9314 | 6635 | 10.90988327 | 6.854773171 | 7.740601495 | 42.06750534 | 40.21249845 | 30.42051132 |
| ENSG00000168398 | - | 15 | 19 | 9 | 72 | 39 | 65 | 0.101258532 | 0.116192328 | 0.045719624 | 0.46438426 | 0.247858683 | 0.438685869 |
| ENSG00000166448 | - | 12 | 10 | 32 | 25 | 93 | 88 | 0.118913086 | 0.089770137 | 0.238626218 | 0.236697157 | 0.86762192 | 0.871828371 |
| ENSG00000113578 | - | 48 | 21 | 19 | 111 | 144 | 109 | 0.420467656 | 0.166645708 | 0.12524625 | 0.929006949 | 1.187553026 | 0.954591967 |
| ENSG00000165879 | - | 25 | 5 | 21 | 59 | 83 | 62 | 0.436085725 | 0.079010598 | 0.27565821 | 0.983305574 | 1.363041618 | 1.081243407 |
| ENSG00000146674 | - | 9480 | 11744 | 8833 | 41875 | 52030 | 26868 | 95.8320849 | 107.5479469 | 67.19401684 | 404.4473986 | 495.1714389 | 271.5425297 |
| ENSG00000169891 | - | 44 | 34 | 73 | 232 | 198 | 152 | 0.179748591 | 0.125827398 | 0.224416845 | 0.905535394 | 0.761512734 | 0.620806602 |
| ENSG00000120937 | - | 1184 | 2062 | 1153 | 3798 | 7658 | 5939 | 78.83263669 | 124.3731098 | 57.77014154 | 241.6094616 | 480.030546 | 395.3371218 |
| ENSG00000236333 | - | 9 | 4 | 36 | 48 | 55 | 73 | 0.047360072 | 0.019068359 | 0.142558172 | 0.241332442 | 0.272478158 | 0.384054293 |
| ENSG00000160963 | - | 15 | 11 | 12 | 53 | 62 | 32 | 0.219618077 | 0.145899228 | 0.132214122 | 0.741408094 | 0.854609423 | 0.468410611 |
| ENSG00000139946 | - | 65 | 58 | 41 | 328 | 204 | 97 | 0.478416188 | 0.386726547 | 0.227089022 | 2.306593055 | 1.413584708 | 0.713779654 |
| ENSG00000164849 | - | 13 | 14 | 22 | 54 | 90 | 36 | 0.190094736 | 0.185454876 | 0.242085731 | 0.754440727 | 1.238991735 | 0.526294897 |
| ENSG00000072657 | - | 64 | 55 | 159 | 238 | 257 | 480 | 0.247471898 | 0.192660236 | 0.462660916 | 0.879280617 | 0.935574121 | 1.855611577 |
| ENSG00000101605 | - | 8 | 14 | 15 | 34 | 52 | 48 | 0.05623078 | 0.089144723 | 0.07934054 | 0.228332465 | 0.344101548 | 0.337306942 |
| ENSG00000170379 | - | 44 | 26 | 36 | 197 | 132 | 56 | 0.317231442 | 0.169816693 | 0.195319599 | 1.357045478 | 0.895976547 | 0.403656078 |
| ENSG00000141052 | - | 42 | 15 | 74 | 81 | 188 | 180 | 0.218672792 | 0.070748983 | 0.289932535 | 0.402935104 | 0.92151513 | 0.936953169 |
| ENSG00000138623 | - | 235 | 247 | 256 | 833 | 954 | 815 | 2.857577951 | 2.720887964 | 2.342555901 | 9.677871523 | 10.92138609 | 9.908040049 |
| ENSG00000010310 | - | 12 | 5 | 33 | 38 | 70 | 59 | 0.117080178 | 0.044193217 | 0.242290196 | 0.354234088 | 0.64298276 | 0.57551157 |
| ENSG00000233818 | - | 15 | 33 | 16 | 107 | 75 | 45 | 0.588382557 | 1.17264337 | 0.472289496 | 4.010115994 | 2.769675921 | 1.764740955 |
| ENSG00000176678 | Fork | 14 | 13 | 46 | 71 | 86 | 83 | 0.172877341 | 0.145424263 | 0.427452285 | 0.837670099 | 0.99978738 | 1.024679507 |
| ENSG00000186479 | - | 958 | 438 | 1436 | 3714 | 2660 | 3006 | 10.10891127 | 4.186937405 | 11.40284155 | 37.4442729 | 26.42528463 | 31.71230227 |
| ENSG00000149591 | - | 905 | 1560 | 1063 | 3218 | 5644 | 3166 | 10.67600647 | 16.67125635 | 9.43655092 | 36.27028063 | 62.68251538 | 37.339722 |
| ENSG00000172061 | - | 7 | 15 | 39 | 41 | 86 | 59 | 0.054453452 | 0.105706603 | 0.228303073 | 0.304730454 | 0.629832386 | 0.458859057 |
| ENSG00000235750 | - | 9 | 15 | 24 | 44 | 64 | 42 | 0.08694727 | 0.13127686 | 0.174479519 | 0.406135304 | 0.582093266 | 0.405660434 |
| ENSG00000198542 | - | 48 | 58 | 80 | 222 | 270 | 100 | 0.359243098 | 0.393240891 | 0.450564485 | 1.587467332 | 1.902436288 | 0.748250673 |
| ENSG00000169031 | - | 1072 | 1110 | 794 | 3475 | 3887 | 2397 | 5.429043599 | 5.092546522 | 3.025999281 | 16.81465574 | 18.53285152 | 12.13658491 |
| ENSG00000100027 | - | 137 | 49 | 104 | 396 | 300 | 232 | 1.343188448 | 0.435207071 | 0.76730759 | 3.70950699 | 2.769088253 | 2.274072395 |
| ENSG00000152822 | - | 26 | 39 | 36 | 112 | 146 | 59 | 0.165208846 | 0.224495707 | 0.172140169 | 0.679958802 | 0.873397547 | 0.374810616 |
| ENSG00000106366 | - | 5403 | 6477 | 9026 | 17642 | 26503 | 20124 | 78.2632908 | 84.99248297 | 98.38711837 | 244.1605993 | 361.4242894 | 291.4320873 |
| ENSG00000213694 | - | 21 | 12 | 22 | 48 | 69 | 54 | 0.071465644 | 0.036994976 | 0.056340471 | 0.156071599 | 0.221068197 | 0.183726456 |
| ENSG00000135318 | - | 668 | 551 | 1141 | 2732 | 2118 | 2283 | 6.809332856 | 5.08818839 | 8.752528228 | 26.60807301 | 20.32606074 | 23.26665412 |
| ENSG00000165072 | - | 790 | 1140 | 1026 | 2764 | 4139 | 2173 | 9.471727644 | 12.38198986 | 9.25697829 | 31.66246889 | 46.71933995 | 26.04724273 |
| ENSG00000126561 | STAT | 42 | 8 | 41 | 72 | 73 | 131 | 0.337282069 | 0.058199256 | 0.247769357 | 0.552434874 | 0.551907105 | 1.051756439 |
| ENSG00000113083 | - | 9085 | 9094 | 7962 | 32068 | 30645 | 16946 | 75.67990642 | 68.62685345 | 49.91115294 | 255.2302564 | 240.3335725 | 141.1311165 |
| ENSG00000005187 | - | 32 | 37 | 36 | 145 | 138 | 33 | 0.189036639 | 0.198007306 | 0.16003622 | 0.818405686 | 0.767492707 | 0.194899116 |
| ENSG00000118523 | - | 10448 | 12822 | 13618 | 32703 | 44760 | 31078 | 206.40336 | 229.4683821 | 202.4494973 | 617.2709237 | 832.4781404 | 613.8137062 |
| ENSG00000111859 | - | 623 | 605 | 957 | 2245 | 2047 | 2013 | 2.778971125 | 2.444752194 | 3.212385814 | 9.567907321 | 8.596328276 | 8.977174829 |
| ENSG00000249456 | - | 29 | 14 | 53 | 55 | 114 | 102 | 2.414453442 | 1.055923257 | 3.32059937 | 4.375101713 | 8.935623273 | 8.490258825 |
| ENSG00000131738 | - | 1263 | 1425 | 2569 | 3878 | 5938 | 4975 | 36.31627486 | 37.11900195 | 55.58809899 | 106.5394316 | 160.7449355 | 143.018082 |
| ENSG00000165959 | - | 64 | 8 | 69 | 134 | 146 | 121 | 0.188350371 | 0.021328483 | 0.152811265 | 0.376787019 | 0.404518627 | 0.35601787 |
| ENSG00000187689 | - | 555 | 1242 | 813 | 2028 | 3457 | 1922 | 24.73022384 | 50.13489877 | 27.26122605 | 86.33912478 | 145.0219049 | 85.62259148 |
| ENSG00000169991 | - | 599 | 458 | 657 | 1354 | 1895 | 1600 | 4.472911835 | 3.098221589 | 3.691889205 | 9.660215326 | 13.32207553 | 11.94492477 |
| ENSG00000140015 | - | 31 | 49 | 95 | 171 | 147 | 153 | 0.116269233 | 0.1664879 | 0.268130669 | 0.612778967 | 0.519062446 | 0.573712701 |
| ENSG00000260604 | - | 466 | 409 | 642 | 1587 | 1371 | 1192 | 3.049966269 | 2.425023267 | 3.16201497 | 9.924089524 | 8.447841094 | 7.799832845 |
| ENSG00000079841 | - | 121 | 220 | 414 | 490 | 768 | 720 | 0.43721652 | 0.720140699 | 1.125722401 | 1.691655403 | 2.612591893 | 2.601019512 |
| ENSG00000261051 | - | 29 | 28 | 24 | 100 | 68 | 54 | 0.639628478 | 0.559462919 | 0.39834632 | 2.107339076 | 1.41201031 | 1.190757906 |
| ENSG00000106688 | - | 159 | 151 | 154 | 456 | 454 | 349 | 1.837672664 | 1.580998509 | 1.339403797 | 5.035475654 | 4.939989723 | 4.032704301 |
| ENSG00000164694 | - | 92 | 100 | 54 | 237 | 347 | 99 | 0.620236825 | 0.610735553 | 0.273957538 | 1.526590986 | 2.202410947 | 0.667274972 |
| ENSG00000139211 | - | 147 | 149 | 335 | 530 | 675 | 416 | 1.717018096 | 1.576620933 | 2.944571598 | 5.914772354 | 7.422675458 | 4.857924814 |
| ENSG00000146005 | - | 23 | 23 | 38 | 90 | 58 | 70 | 0.235492089 | 0.213333867 | 0.292787046 | 0.880431604 | 0.55908217 | 0.716549913 |
| ENSG00000206538 | - | 5819 | 4427 | 6839 | 16926 | 13885 | 14122 | 21.14851953 | 14.5755388 | 18.70440904 | 58.77472111 | 47.50908425 | 51.31303958 |
| ENSG00000166106 | - | 669 | 497 | 688 | 1521 | 1825 | 1520 | 5.449129822 | 3.6672533 | 4.217054846 | 11.83681549 | 13.99468307 | 12.37783091 |
| ENSG00000171401 | - | 80 | 59 | 100 | 168 | 283 | 166 | 1.26900497 | 0.847830098 | 1.193694273 | 2.546169108 | 4.226286488 | 2.632578588 |
| ENSG00000186352 | - | 64 | 139 | 130 | 344 | 273 | 229 | 1.146236117 | 2.255238086 | 1.752093349 | 5.886500486 | 4.603158439 | 4.100431089 |
| ENSG00000011028 | - | 268 | 144 | 335 | 599 | 652 | 652 | 1.805196567 | 0.878689991 | 1.698064904 | 3.854971346 | 4.1346282 | 4.390734949 |
| ENSG00000082497 | - | 289 | 276 | 376 | 817 | 817 | 764 | 2.442657575 | 2.113281305 | 2.391510906 | 6.597682775 | 6.501086806 | 6.455918303 |
| ENSG00000118515 | - | 217 | 369 | 392 | 675 | 827 | 898 | 1.00350867 | 1.545863858 | 1.364166048 | 2.982426813 | 3.600526955 | 4.151811739 |
| ENSG00000108821 | - | 1670 | 3001 | 1774 | 4288 | 8031 | 3916 | 7.858922965 | 12.79369633 | 6.282318497 | 19.27995271 | 35.58077048 | 18.42422227 |
| ENSG00000049192 | - | 26 | 25 | 39 | 63 | 78 | 77 | 0.14023564 | 0.12215424 | 0.158295816 | 0.324661081 | 0.396076265 | 0.415217548 |
| ENSG00000171219 | - | 35 | 44 | 101 | 113 | 133 | 176 | 0.264605288 | 0.301346835 | 0.57460794 | 0.816231866 | 0.94663221 | 1.330280005 |
| ENSG00000103647 | - | 60 | 55 | 46 | 175 | 147 | 80 | 0.693461383 | 0.575860384 | 0.400081654 | 1.932474209 | 1.599512091 | 0.924402132 |
| ENSG00000166482 | - | 84 | 81 | 61 | 146 | 260 | 152 | 1.803644076 | 1.575578532 | 0.985646452 | 2.995222134 | 5.255863327 | 3.262984886 |
| ENSG00000123843 | - | 29 | 93 | 58 | 140 | 177 | 115 | 0.416285076 | 1.209370233 | 0.626528183 | 1.920107335 | 2.392022564 | 1.650405282 |
| ENSG00000138356 | - | 277 | 251 | 547 | 927 | 839 | 750 | 2.303728805 | 1.891075148 | 3.42340818 | 7.366071748 | 6.569203357 | 6.236095646 |
| ENSG00000159263 | Others | 74 | 50 | 90 | 227 | 190 | 93 | 0.446859072 | 0.273522052 | 0.408979126 | 1.309692509 | 1.080168783 | 0.561463758 |
| ENSG00000166145 | - | 45 | 62 | 69 | 101 | 121 | 185 | 0.58705363 | 0.732724031 | 0.677382743 | 1.25890009 | 1.486106028 | 2.412886607 |
| ENSG00000113389 | - | 1517 | 1111 | 2316 | 3909 | 3313 | 4066 | 7.104905244 | 4.713790217 | 8.162647435 | 17.49214251 | 14.60808567 | 19.03881897 |
| ENSG00000115380 | - | 10425 | 10538 | 9120 | 29961 | 26814 | 14623 | 121.5837161 | 111.3373874 | 80.04126833 | 333.856842 | 294.4151243 | 170.5044625 |
| ENSG00000101333 | - | 5133 | 4007 | 7055 | 14830 | 11220 | 11091 | 34.37446868 | 24.3090258 | 35.55343136 | 94.88782849 | 70.73864694 | 74.25665054 |
| ENSG00000079931 | - | 91 | 101 | 167 | 320 | 297 | 196 | 0.794425761 | 0.798760872 | 1.097104943 | 2.669110366 | 2.440998641 | 1.710676615 |
| ENSG00000119681 | - | 4637 | 4023 | 5632 | 10474 | 12375 | 10014 | 23.56889701 | 18.52403862 | 21.54192865 | 50.86503638 | 59.21701779 | 50.8873307 |
| ENSG00000146094 | - | 104 | 93 | 99 | 274 | 288 | 123 | 0.996391234 | 0.807166241 | 0.713758885 | 2.508139851 | 2.597695298 | 1.178152721 |
| ENSG00000243753 | - | 59 | 75 | 44 | 149 | 197 | 69 | 0.495231783 | 0.570297205 | 0.277925853 | 1.194943559 | 1.556760792 | 0.579035924 |
| ENSG00000027075 | - | 70 | 69 | 94 | 182 | 201 | 142 | 0.366187597 | 0.32699278 | 0.370043873 | 0.909665251 | 0.98992162 | 0.742666536 |
| ENSG00000175274 | - | 924 | 811 | 597 | 1734 | 2245 | 1410 | 5.812915257 | 4.621961855 | 2.82628612 | 10.42259208 | 13.29650581 | 8.868313837 |
| ENSG00000162373 | - | 107 | 84 | 81 | 231 | 224 | 165 | 2.312543429 | 1.644632575 | 1.317379536 | 4.770047752 | 4.557779386 | 3.565249967 |
| ENSG00000074590 | - | 440 | 299 | 678 | 926 | 1129 | 1038 | 2.90116483 | 1.785970981 | 3.364102312 | 5.83358246 | 7.008302372 | 6.842534591 |
| ENSG00000153162 | - | 2006 | 1786 | 1660 | 4725 | 4664 | 2996 | 33.3426377 | 26.89267241 | 20.76332428 | 75.03700036 | 72.9838431 | 49.78640347 |
| ENSG00000004799 | - | 55 | 78 | 79 | 231 | 122 | 116 | 0.56238557 | 0.722519486 | 0.607880686 | 2.256774086 | 1.174439019 | 1.185848993 |
| ENSG00000175161 | - | 161 | 253 | 199 | 584 | 430 | 342 | 0.744687748 | 1.060113429 | 0.692661713 | 2.580868081 | 1.872474662 | 1.581518801 |
| ENSG00000141738 | - | 188 | 156 | 142 | 245 | 547 | 293 | 2.415078391 | 1.815438175 | 1.372718546 | 3.007073852 | 6.615457304 | 3.763058101 |
| ENSG00000198948 | - | 304 | 325 | 444 | 723 | 808 | 775 | 1.60703851 | 1.556393985 | 1.766263298 | 3.651704483 | 4.021269898 | 4.095946954 |
| ENSG00000177181 | - | 679 | 295 | 564 | 1553 | 1065 | 731 | 3.493485126 | 1.374974432 | 2.183675001 | 7.634229419 | 5.158671744 | 3.760160837 |
| ENSG00000157680 | - | 36 | 49 | 104 | 100 | 161 | 127 | 0.106982775 | 0.131914009 | 0.232575772 | 0.283933074 | 0.450439424 | 0.377324494 |
| ENSG00000075618 | - | 769 | 1209 | 1431 | 1561 | 2454 | 3072 | 9.624506453 | 13.70761542 | 13.47756814 | 18.6663609 | 28.91517655 | 38.43910439 |
| ENSG00000187498 | - | 10573 | 9201 | 11852 | 20359 | 27350 | 19771 | 33.14923425 | 26.13327156 | 27.96318818 | 60.98686901 | 80.72940672 | 61.97318624 |
| ENSG00000257671 | - | 127 | 133 | 158 | 299 | 361 | 229 | 3.466255594 | 3.288455625 | 3.245142137 | 7.797092345 | 9.276053102 | 6.248737277 |
| ENSG00000144583 | - | 174 | 194 | 310 | 309 | 581 | 509 | 1.807989648 | 1.826130892 | 2.423975127 | 3.067678182 | 5.683580104 | 5.287670619 |
| ENSG00000115414 | - | 7924 | 4852 | 10026 | 18783 | 15940 | 13018 | 21.14636828 | 11.72993483 | 20.13440495 | 47.89177244 | 40.04781292 | 34.73245747 |
| ENSG00000058335 | - | 45 | 69 | 78 | 146 | 126 | 127 | 0.209421287 | 0.290898099 | 0.273163511 | 0.649180988 | 0.552049495 | 0.590897226 |
| ENSG00000108846 | - | 60 | 61 | 112 | 120 | 242 | 117 | 0.288377222 | 0.265596911 | 0.405086248 | 0.551055798 | 1.095025494 | 0.562206012 |
| ENSG00000185100 | - | 125 | 231 | 223 | 400 | 443 | 352 | 1.000165443 | 1.674392593 | 1.342723127 | 3.057922331 | 3.337065415 | 2.815816932 |
| ENSG00000079308 | - | 722 | 327 | 570 | 1207 | 1203 | 1025 | 2.04913203 | 0.840744392 | 1.217383396 | 3.272989721 | 3.21438234 | 2.908416035 |
| ENSG00000020181 | - | 79 | 66 | 127 | 175 | 189 | 187 | 0.516981141 | 0.391268628 | 0.625419048 | 1.094183811 | 1.164417101 | 1.223458201 |
| ENSG00000073464 | - | 158 | 147 | 256 | 404 | 399 | 336 | 1.075387784 | 0.906376901 | 1.31119641 | 2.627205255 | 2.556701754 | 2.286373669 |
| ENSG00000136490 | - | 46 | 84 | 64 | 93 | 180 | 126 | 0.44795608 | 0.741037818 | 0.46900487 | 0.865296963 | 1.650248211 | 1.226727411 |
| ENSG00000214517 | - | 2945 | 3721 | 4429 | 6908 | 8485 | 7181 | 17.18851964 | 19.67416584 | 19.4526436 | 38.52211573 | 46.62342608 | 41.90231555 |
| ENSG00000198053 | - | 2138 | 1975 | 2982 | 4108 | 5297 | 4983 | 19.49327974 | 16.31277314 | 20.45995281 | 35.78592739 | 45.46805211 | 45.42218504 |
| ENSG00000181274 | - | 61 | 105 | 102 | 171 | 220 | 151 | 1.273685608 | 1.986118643 | 1.602700427 | 3.41140392 | 4.324682703 | 3.15216741 |
| ENSG00000112139 | - | 5104 | 4963 | 5265 | 8817 | 12476 | 10161 | 16.62393827 | 14.64370628 | 12.90450652 | 27.43776422 | 38.25584329 | 33.08717008 |
| ENSG00000261068 | - | 43 | 73 | 65 | 136 | 94 | 135 | 0.980231213 | 1.507531565 | 1.115047075 | 2.96212654 | 2.017377103 | 3.076760994 |
| ENSG00000144642 | - | 773 | 584 | 870 | 1819 | 1431 | 1272 | 1.947680266 | 1.333013352 | 1.649594488 | 4.379005556 | 3.394509009 | 3.204241213 |
| ENSG00000235609 | - | 125 | 129 | 187 | 347 | 238 | 297 | 0.97418712 | 0.910763386 | 1.096714918 | 2.583845086 | 1.746258323 | 2.314135263 |
| ENSG00000125730 | - | 128 | 62 | 205 | 241 | 231 | 306 | 0.710203934 | 0.31163647 | 0.855946734 | 1.277599619 | 1.206658137 | 1.697440075 |
| ENSG00000105856 | HMG | 902 | 534 | 1087 | 2002 | 1677 | 1403 | 6.562634925 | 3.519625432 | 5.951425837 | 13.91683275 | 11.48692888 | 10.2053828 |
| ENSG00000133424 | - | 402 | 297 | 611 | 788 | 1043 | 777 | 1.738137239 | 1.163316647 | 1.988011973 | 3.255283791 | 4.245623528 | 3.358759831 |
| ENSG00000164176 | - | 1324 | 1093 | 1374 | 3255 | 2494 | 1968 | 10.66206343 | 7.973645453 | 8.32644818 | 25.04429995 | 18.90814341 | 15.84449097 |
| ENSG00000054277 | - | 222 | 179 | 234 | 441 | 496 | 353 | 2.760521219 | 2.016390435 | 2.189645187 | 5.239397298 | 5.806560475 | 4.388466034 |
| ENSG00000185499 | - | 343 | 254 | 289 | 715 | 740 | 371 | 3.360021562 | 2.254058293 | 2.130421622 | 6.69204131 | 6.824625521 | 3.633471639 |
| ENSG00000213713 | - | 97 | 55 | 123 | 260 | 140 | 150 | 5.007979236 | 2.572390856 | 4.778764963 | 12.82533626 | 6.804841011 | 7.742513388 |
| ENSG00000164197 | - | 417 | 356 | 522 | 1052 | 807 | 739 | 3.545930682 | 2.742380213 | 3.340292233 | 8.547016717 | 6.460511189 | 6.282587495 |
| ENSG00000001617 | - | 145 | 106 | 119 | 227 | 282 | 246 | 1.388335744 | 0.919424031 | 0.857419268 | 2.076619784 | 2.541995475 | 2.354840686 |
| ENSG00000114796 | - | 1160 | 1162 | 1614 | 3195 | 2217 | 2419 | 6.196631956 | 5.623249114 | 6.488152029 | 16.30696739 | 11.14968423 | 12.91913695 |
| ENSG00000133216 | - | 738 | 559 | 814 | 1362 | 1817 | 1070 | 2.724611767 | 1.86957786 | 2.261478406 | 4.804298695 | 6.315421702 | 3.949407663 |
| ENSG00000152409 | - | 752 | 471 | 1058 | 1712 | 1329 | 1431 | 3.842971454 | 2.180488032 | 4.068691482 | 8.359065057 | 6.3940122 | 7.311203501 |
| ENSG00000165312 | - | 750 | 630 | 945 | 1095 | 1458 | 2005 | 11.815797 | 8.991368341 | 11.20349024 | 16.48240206 | 21.62511588 | 31.58028574 |
| ENSG00000108379 | - | 60 | 70 | 166 | 166 | 173 | 220 | 0.639699725 | 0.676092995 | 1.331842213 | 1.69097672 | 1.736481571 | 2.345025205 |
| ENSG00000170442 | - | 524 | 498 | 298 | 800 | 1099 | 784 | 10.11395371 | 8.70768078 | 4.3283833 | 14.75313405 | 19.97038919 | 15.12884097 |
| ENSG00000168453 | - | 345 | 268 | 343 | 343 | 728 | 809 | 1.814849385 | 1.277143713 | 1.357798697 | 1.723932434 | 3.605388213 | 4.254709723 |
| ENSG00000278948 | - | 58 | 143 | 230 | 238 | 317 | 253 | 1.449455555 | 3.237400498 | 4.325382508 | 5.682749253 | 7.458222465 | 6.321168267 |
| ENSG00000133107 | - | 452 | 324 | 594 | 1100 | 741 | 794 | 4.966909595 | 3.225346018 | 4.911951156 | 11.54901998 | 7.665936018 | 8.723047624 |
| ENSG00000102452 | - | 171 | 185 | 242 | 470 | 406 | 278 | 0.864497487 | 0.847271963 | 0.920667399 | 2.270231315 | 1.932381274 | 1.405116525 |
| ENSG00000154736 | - | 2998 | 2413 | 5781 | 6239 | 6436 | 8093 | 15.29709752 | 11.15368079 | 22.19729993 | 30.41564422 | 30.91666242 | 41.28448466 |
| ENSG00000173801 | - | 1914 | 1611 | 1801 | 3582 | 3693 | 3065 | 17.89587811 | 13.64552038 | 12.67198256 | 31.99935969 | 32.50794744 | 28.65111178 |
| ENSG00000151414 | - | 2039 | 1247 | 2588 | 4759 | 3439 | 2997 | 17.17728077 | 9.516716335 | 16.40669992 | 38.30518115 | 27.27523658 | 25.24200522 |
| ENSG00000073910 | - | 210 | 161 | 332 | 505 | 404 | 415 | 0.514807424 | 0.357548421 | 0.612467945 | 1.182827768 | 0.93240809 | 1.017123114 |
| ENSG00000011201 | - | 1604 | 1238 | 1807 | 3579 | 3050 | 2304 | 10.39364186 | 7.26720667 | 8.811326839 | 22.15795671 | 18.60640137 | 14.9260805 |
| ENSG00000091592 | - | 3930 | 4519 | 4630 | 5811 | 8979 | 10166 | 14.17832908 | 14.76924604 | 12.5699418 | 20.03032538 | 30.49717663 | 36.6676036 |
| ENSG00000127329 | - | 587 | 440 | 1154 | 1561 | 1264 | 1205 | 1.865210199 | 1.266560207 | 2.759403833 | 4.739114595 | 3.781254562 | 3.828041593 |
| ENSG00000122707 | - | 871 | 659 | 1028 | 1746 | 1612 | 1503 | 6.65787551 | 4.563377663 | 5.913306693 | 12.75164994 | 11.60063372 | 11.48620113 |
| ENSG00000100302 | - | 70 | 120 | 173 | 218 | 295 | 161 | 0.947987996 | 1.472209197 | 1.763074546 | 2.820758586 | 3.761196089 | 2.179870001 |
| ENSG00000112414 | - | 276 | 211 | 675 | 810 | 552 | 744 | 1.505170494 | 1.04241956 | 2.770130067 | 4.220522891 | 2.834097935 | 4.056481226 |
| ENSG00000169432 | - | 552 | 295 | 503 | 1054 | 873 | 648 | 1.856241845 | 0.898671521 | 1.272867624 | 3.386419209 | 2.763814471 | 2.178564425 |
| ENSG00000166046 | - | 296 | 289 | 442 | 812 | 509 | 585 | 2.085614893 | 1.844691443 | 2.343605193 | 5.466420854 | 3.376442455 | 4.120958063 |
| ENSG00000173894 | - | 98 | 123 | 121 | 225 | 241 | 177 | 0.796263242 | 0.905355587 | 0.739836471 | 1.746697656 | 1.843515514 | 1.437817547 |
| ENSG00000132434 | - | 777 | 822 | 871 | 1490 | 1868 | 1297 | 6.858326452 | 6.572830524 | 5.78542228 | 12.56573886 | 15.52291103 | 11.4455596 |
| ENSG00000092295 | - | 889 | 1061 | 1038 | 1721 | 2108 | 1771 | 10.99828515 | 11.89110135 | 9.663631084 | 20.3427055 | 24.55234527 | 21.90492125 |
| ENSG00000211455 | - | 1236 | 1275 | 1735 | 2981 | 2664 | 2233 | 8.252080238 | 7.711496014 | 8.716943371 | 19.01566748 | 16.74473788 | 14.9050561 |
| ENSG00000095951 | zf-C2H2 | 847 | 742 | 1569 | 2135 | 1735 | 1866 | 3.564794063 | 2.829035669 | 4.96928326 | 8.585261776 | 6.8746355 | 7.851680076 |
| ENSG00000120162 | - | 64 | 128 | 147 | 249 | 190 | 179 | 0.417283644 | 0.756040115 | 0.721254435 | 1.551155589 | 1.166283551 | 1.166821276 |
| ENSG00000100311 | - | 178 | 203 | 225 | 373 | 448 | 303 | 2.18690788 | 2.259383784 | 2.080236822 | 4.378485805 | 5.181883716 | 3.721800041 |
| ENSG00000135245 | - | 286 | 273 | 231 | 549 | 615 | 330 | 6.38424446 | 5.520642246 | 3.880388682 | 11.70901727 | 12.92461909 | 7.36473858 |
| ENSG00000152779 | - | 349 | 299 | 706 | 709 | 754 | 947 | 3.335360407 | 2.588642117 | 5.077405129 | 6.473933058 | 6.784031407 | 9.048305218 |
| ENSG00000091986 | - | 26909 | 13442 | 21531 | 46249 | 41146 | 28654 | 95.12672925 | 43.04794055 | 57.27814981 | 156.2111321 | 136.9404622 | 101.2721857 |
| ENSG00000122786 | - | 13470 | 10445 | 13938 | 25388 | 25727 | 18953 | 63.23447704 | 44.41997012 | 49.23869905 | 113.8726565 | 113.7037122 | 88.9537413 |
| ENSG00000187398 | - | 220 | 245 | 457 | 530 | 641 | 477 | 1.148534805 | 1.158700106 | 1.795384428 | 2.643637943 | 3.150494104 | 2.489658497 |
| ENSG00000004399 | - | 2586 | 2145 | 2166 | 4053 | 5735 | 3126 | 12.27079133 | 9.22050758 | 7.734322496 | 18.37491256 | 25.61985305 | 14.82971978 |
| ENSG00000138641 | - | 232 | 224 | 271 | 611 | 395 | 329 | 1.535621442 | 1.343159794 | 1.349849182 | 3.864043615 | 2.461458058 | 2.177168295 |
| ENSG00000130518 | - | 122 | 125 | 174 | 155 | 283 | 321 | 0.615832696 | 0.571605512 | 0.660955065 | 0.747548265 | 1.344895405 | 1.619973333 |
| ENSG00000163235 | - | 318 | 276 | 338 | 637 | 701 | 380 | 2.960715419 | 2.327888151 | 2.368132619 | 5.66648319 | 6.144502498 | 3.537146623 |

### Supplemental Table 8 - List of up regulated genes by RNAseq

| gene_id | CH_0323 | CH_0407 | CH_0603 | DMSO_0323 | DMSO_0407 | DMSO_0603 | CHIR | DMSO | log2FoldChange | pvalue | padj |
|---|---|---|---|---|---|---|---|---|---|---|---|
| ENSG00000203837 | 34.3467814 | 6.80640192 | 67.039989 | 3.076535635 | 0 | 2.146179214 | 36.0643908 | 1.74090495 | 4.382717783 | 4.69E-05 | 0.00285117 |
| ENSG00000259070 | 30.0534338 | 21.3915489 | 27.4621642 | 2.051023757 | 3.031492406 | 0 | 26.3023823 | 1.69417205 | 3.940635771 | 0.00013616 | 0.00657973 |
| ENSG00000154545 | 22.5400753 | 294.619969 | 70.2708319 | 8.204095028 | 18.18895443 | 10.73089607 | 129.143625 | 12.3746485 | 3.382914894 | 0.00114513 | 0.03244458 |
| ENSG00000172348 | 24.6867492 | 16.5298332 | 23.4236106 | 1.025511878 | 4.041989874 | 2.146179214 | 21.546731 | 2.40456032 | 3.160243114 | 0.00156051 | 0.04076723 |
| ENSG00000187950 | 24.6867492 | 40.8384115 | 16.1542142 | 4.102047514 | 0 | 5.365448036 | 27.2264583 | 3.15583185 | 3.115987071 | 0.00126038 | 0.03489426 |
| ENSG00000119686 | 59.0335306 | 59.312931 | 67.039989 | 0 | 10.10497469 | 13.95016489 | 61.7954835 | 8.01837986 | 2.951150115 | 2.50E-05 | 0.00181347 |
| ENSG00000042781 | 16.1000538 | 35.0043527 | 58.1551712 | 0 | 6.062984811 | 8.584716857 | 36.4198592 | 4.88256722 | 2.908127233 | 0.00103856 | 0.0301593 |
| ENSG00000223861 | 15.0267169 | 49.5894997 | 20.1927678 | 3.076535635 | 4.041989874 | 4.292358429 | 28.2696614 | 3.80362798 | 2.894722021 | 0.00104095 | 0.0301593 |
| ENSG00000196872 | 86.9402905 | 117.653519 | 144.580217 | 9.229606906 | 32.33591899 | 10.73089607 | 116.391342 | 17.4321407 | 2.736786501 | 6.14E-08 | 1.04E-05 |
| ENSG00000133083 | 165.293886 | 33.0596664 | 189.004306 | 19.48472569 | 12.12596962 | 26.82724018 | 129.119286 | 19.4793118 | 2.731855813 | 1.38E-06 | 0.00016445 |
| ENSG00000183036 | 1811.79272 | 2897.58253 | 2070.16255 | 279.9647428 | 323.3591899 | 430.3089325 | 2259.84593 | 344.544288 | 2.713988665 | 1.20E-34 | 1.89E-30 |
| ENSG00000251003 | 125.58042 | 122.515234 | 142.157085 | 15.38267818 | 22.23094431 | 22.53488175 | 130.084246 | 20.0495014 | 2.699260167 | 2.26E-11 | 8.09E-09 |
| ENSG00000270605 | 18.2467276 | 29.1702939 | 31.5007177 | 3.076535635 | 6.062984811 | 3.219268821 | 26.3059131 | 4.11959642 | 2.674367472 | 0.00166726 | 0.04256784 |
| ENSG00000129151 | 24.6867492 | 65.1469898 | 26.6544535 | 3.076535635 | 11.11547215 | 5.365448036 | 38.8293975 | 6.51915194 | 2.570987394 | 0.00079579 | 0.02467741 |
| ENSG00000261235 | 21.4667384 | 53.4788722 | 24.2313213 | 6.153071271 | 2.020994937 | 8.584716857 | 33.0589773 | 5.58626102 | 2.570244509 | 0.00149089 | 0.03933995 |
| ENSG00000270127 | 19.3200646 | 58.3405878 | 22.6158999 | 5.127559392 | 7.07348228 | 5.365448036 | 33.4255174 | 5.85549657 | 2.511512252 | 0.00141079 | 0.03779626 |
| ENSG00000107562 | 48.3001614 | 98.2066562 | 102.57926 | 15.38267818 | 9.094477217 | 25.75415057 | 83.0286926 | 16.7437687 | 2.316564444 | 1.56E-05 | 0.00122651 |
| ENSG00000077498 | 845.789493 | 1160.9777 | 1184.1039 | 255.3524577 | 223.3199405 | 182.4252332 | 1063.6237 | 220.365877 | 2.271061013 | 7.85E-26 | 2.47E-22 |
| ENSG00000165092 | 42.9334768 | 132.238666 | 105.810103 | 17.43370193 | 13.13646709 | 27.90032979 | 93.6607486 | 19.4901663 | 2.269669866 | 1.75E-05 | 0.00134478 |
| ENSG00000140961 | 129.873767 | 141.962097 | 119.541185 | 6.153071271 | 35.3674114 | 41.85049468 | 130.459017 | 27.7903258 | 2.232403235 | 8.00E-06 | 0.00072817 |
| ENSG00000166473 | 1142.03048 | 1455.59767 | 1228.52799 | 258.4289934 | 324.3696874 | 263.9800434 | 1275.38538 | 282.259575 | 2.175584295 | 1.76E-29 | 1.39E-25 |
| ENSG00000278035 | 41.8601399 | 66.1193329 | 31.5007177 | 4.102047514 | 16.1679595 | 10.73089607 | 46.4933968 | 10.3336344 | 2.167197855 | 0.0011618 | 0.032723 |
| ENSG00000107165 | 2462.23489 | 1786.19433 | 2062.08545 | 767.0828851 | 428.4509267 | 314.4152549 | 2103.50489 | 503.316356 | 2.062867039 | 4.13E-15 | 2.25E-12 |
| ENSG00000283031 | 46.1534875 | 35.0043527 | 52.5011962 | 8.204095028 | 7.07348228 | 17.16943371 | 44.5530122 | 10.8156703 | 2.048021661 | 0.00123416 | 0.03434928 |
| ENSG00000132386 | 383.18128 | 317.956204 | 683.323262 | 111.7807948 | 63.66134052 | 160.9634411 | 461.486915 | 112.135192 | 2.043586767 | 2.95E-09 | 6.93E-07 |
| ENSG00000164175 | 398.207997 | 412.273487 | 457.971973 | 112.8063066 | 101.0497469 | 103.0166023 | 422.817819 | 105.624219 | 2.001665261 | 1.26E-17 | 1.17E-14 |
| ENSG00000041353 | 787.829299 | 387.964909 | 1192.18101 | 155.8778055 | 151.5746203 | 298.3189108 | 789.325073 | 201.923779 | 1.967341127 | 0.0003037 | 0.01215436 |
| ENSG00000109819 | 266.187556 | 281.979508 | 453.125709 | 61.53071271 | 83.87128989 | 113.7474984 | 333.764258 | 86.383167 | 1.952853394 | 5.43E-10 | 1.56E-07 |
| ENSG00000125965 | 191.053972 | 165.298332 | 182.542621 | 23.5867732 | 78.81880255 | 42.92358429 | 179.631642 | 48.4430533 | 1.889381645 | 2.11E-06 | 0.00023634 |
| ENSG00000148053 | 103.040344 | 103.068372 | 163.157564 | 17.43370193 | 44.46188862 | 40.77740507 | 123.08876 | 34.2243319 | 1.849223341 | 2.16E-05 | 0.00158636 |
| ENSG00000197915 | 81.5736059 | 94.3172837 | 299.660674 | 34.86740387 | 33.34641646 | 65.45846604 | 158.517188 | 44.5574288 | 1.836037585 | 0.00018297 | 0.00823513 |
| ENSG00000139289 | 1193.55065 | 1363.22507 | 1890.85078 | 470.7099522 | 326.3906823 | 474.3056064 | 1482.54217 | 423.80208 | 1.807500206 | 4.47E-16 | 3.20E-13 |
| ENSG00000118322 | 827.542765 | 592.156967 | 682.515551 | 226.6381251 | 211.1939709 | 183.4983228 | 700.738428 | 207.11014 | 1.757627217 | 5.53E-16 | 3.79E-13 |
| ENSG00000158270 | 203.934015 | 175.021764 | 165.580696 | 76.91339088 | 46.48288355 | 38.63122586 | 181.512158 | 54.0091668 | 1.746830662 | 5.30E-07 | 7.51E-05 |
| ENSG00000135253 | 1702.31235 | 2653.5244 | 2284.20589 | 477.8885354 | 853.8703609 | 696.435155 | 2213.34755 | 676.064684 | 1.711117603 | 5.74E-14 | 2.74E-11 |
| ENSG00000198203 | 701.962346 | 921.781288 | 951.483218 | 273.8116715 | 252.6243671 | 261.8338641 | 858.40895 | 262.756634 | 1.708685395 | 2.92E-17 | 2.30E-14 |
| ENSG00000182612 | 6755.58257 | 9798.30173 | 7038.39114 | 2289.968025 | 2872.844303 | 2653.750598 | 7864.09181 | 2605.52098 | 1.593694902 | 1.35E-18 | 1.52E-15 |
| ENSG00000257681 | 82.6469428 | 80.7044798 | 63.8091462 | 13.33165442 | 38.3989038 | 23.60797136 | 75.7201896 | 25.1128432 | 1.589790061 | 0.00134214 | 0.03645298 |
| ENSG00000276772 | 96.6003228 | 77.7874505 | 71.8862533 | 18.45921381 | 40.41989874 | 24.68106096 | 82.0913422 | 27.8533912 | 1.556525477 | 0.000766 | 0.02418185 |
| ENSG00000272899 | 582.821947 | 830.381034 | 823.057215 | 272.7861597 | 244.5403874 | 250.0298785 | 745.420065 | 255.785475 | 1.543894571 | 5.44E-13 | 2.32E-10 |
| ENSG00000170989 | 319.854402 | 263.504988 | 462.818238 | 133.3165442 | 74.77681267 | 167.4019787 | 348.725876 | 125.165112 | 1.480566735 | 3.96E-06 | 0.00040656 |
| ENSG00000166963 | 205.007352 | 75.8427642 | 105.002392 | 39.99496326 | 44.46188862 | 53.65448036 | 128.617503 | 46.0371107 | 1.48012574 | 0.00053066 | 0.0183165 |
| ENSG00000143772 | 1230.04411 | 784.680906 | 1268.91353 | 388.6690019 | 361.7580937 | 434.6012909 | 1094.54618 | 395.009462 | 1.470764805 | 1.51E-11 | 5.54E-09 |
| ENSG00000164627 | 247.940828 | 255.726243 | 251.198031 | 113.8318185 | 82.86079242 | 90.139527 | 251.621701 | 95.6107126 | 1.395995701 | 2.73E-07 | 4.14E-05 |
| ENSG00000180332 | 81.5736059 | 107.930088 | 90.4635997 | 41.02047514 | 39.40940127 | 27.90032979 | 93.322431 | 36.1100687 | 1.368915697 | 0.00077666 | 0.02437207 |
| ENSG00000285437 | 97.6736597 | 106.957744 | 76.7325176 | 27.68882072 | 27.28343165 | 54.72756996 | 93.7879739 | 36.5666074 | 1.359961176 | 0.00183551 | 0.04567909 |
| ENSG00000170289 | 172.807244 | 96.2619699 | 125.19516 | 69.73480773 | 43.45139115 | 41.85049468 | 131.421458 | 51.6788979 | 1.344596851 | 0.00051303 | 0.01787994 |
| ENSG00000138316 | 481.928277 | 601.880398 | 538.743044 | 184.5921381 | 190.9840216 | 276.8571186 | 540.850573 | 217.477759 | 1.31540053 | 1.93E-08 | 3.70E-06 |
| ENSG00000163762 | 257.600861 | 256.698587 | 440.202338 | 181.5156025 | 127.322681 | 79.40863093 | 318.167262 | 129.415638 | 1.298578395 | 8.70E-05 | 0.00464823 |
| ENSG00000108924 | 122.360409 | 246.002812 | 243.120924 | 48.19905829 | 95.99725951 | 105.1627815 | 203.828048 | 83.1196998 | 1.296409044 | 0.00052311 | 0.01815092 |
| ENSG00000138642 | 148.120495 | 81.676823 | 142.157085 | 42.04598702 | 42.44089368 | 67.60464525 | 123.984801 | 50.6971753 | 1.291822173 | 0.00108243 | 0.03105918 |
| ENSG00000235718 | 1907.31971 | 1865.92647 | 2276.9365 | 648.1235072 | 696.2327558 | 1136.401894 | 2016.72756 | 826.919386 | 1.2867246 | 3.55E-09 | 8.09E-07 |
| ENSG00000248174 | 173.880581 | 175.994107 | 189.812017 | 66.6582721 | 65.68233545 | 93.35879582 | 179.895568 | 75.2331345 | 1.259420555 | 4.20E-05 | 0.00262486 |
| ENSG00000119922 | 180.320603 | 92.3725974 | 136.50311 | 65.63276022 | 58.60885318 | 49.36212193 | 136.39877 | 57.8679118 | 1.235211934 | 0.00084628 | 0.02593672 |
| ENSG00000210194 | 153.48718 | 399.633027 | 177.696356 | 89.21953343 | 101.0497469 | 121.2591256 | 243.605521 | 103.842802 | 1.230375667 | 0.00070239 | 0.02267361 |
| ENSG00000189007 | 178.173929 | 292.675282 | 276.237063 | 115.8828423 | 94.98676204 | 121.2591256 | 249.028758 | 110.709577 | 1.171504615 | 6.25E-05 | 0.00355509 |
| ENSG00000135069 | 275.847588 | 449.222526 | 479.780163 | 196.8982807 | 181.8895443 | 156.6710826 | 401.616759 | 178.486303 | 1.171247177 | 9.98E-06 | 0.00085008 |
| ENSG00000132481 | 373.521248 | 478.39282 | 534.704491 | 176.3880431 | 162.6900924 | 277.9302083 | 462.206186 | 205.669448 | 1.170084577 | 8.61E-06 | 0.00075805 |
| ENSG00000155761 | 343.467814 | 482.282193 | 446.664023 | 234.8422202 | 189.9735241 | 147.0132762 | 424.13801 | 190.609673 | 1.154049239 | 5.43E-06 | 0.00052847 |
| ENSG00000253417 | 2250.78752 | 2000.10982 | 2468.36393 | 1068.583377 | 892.2692647 | 1068.797249 | 2239.75376 | 1009.8833 | 1.149431286 | 7.58E-12 | 3.06E-09 |
| ENSG00000087303 | 1070.11691 | 436.582066 | 864.250461 | 495.3222373 | 360.7475963 | 212.4717422 | 790.316479 | 356.180525 | 1.149294698 | 0.0002972 | 0.01203176 |
| ENSG00000203727 | 457.241528 | 540.622781 | 382.047166 | 173.3115075 | 174.8160621 | 274.7109394 | 459.970492 | 207.612836 | 1.147860492 | 7.68E-06 | 0.00071177 |
| ENSG00000210174 | 157.780527 | 194.468626 | 124.38745 | 56.40315331 | 76.79780761 | 82.62789975 | 158.878868 | 71.9429536 | 1.142077005 | 0.00066135 | 0.02156993 |
| ENSG00000143387 | 1970.64658 | 1620.896 | 1788.27151 | 943.4709282 | 784.1460356 | 726.481664 | 1793.27137 | 818.032876 | 1.132113549 | 1.20E-10 | 3.78E-08 |
| ENSG00000280744 | 177.100592 | 159.464273 | 220.505024 | 87.16850967 | 57.59835571 | 118.0398568 | 185.689963 | 87.6022407 | 1.086462512 | 0.00117806 | 0.03308018 |
| ENSG00000183828 | 459.388202 | 467.697046 | 264.121403 | 170.2349718 | 240.4983975 | 151.3056346 | 397.068883 | 187.346335 | 1.081595047 | 0.00014838 | 0.00704037 |
| ENSG00000175592 | 335.954456 | 534.788722 | 659.09194 | 156.9033174 | 259.6978494 | 310.1228965 | 509.945039 | 242.241354 | 1.075546006 | 0.0002611 | 0.01082395 |
| ENSG00000005249 | 217.887395 | 208.08143 | 315.007177 | 105.6277235 | 114.1862139 | 133.0631113 | 246.992001 | 117.625683 | 1.072708688 | 0.00016348 | 0.00757456 |
| ENSG00000157388 | 309.121033 | 317.956204 | 419.201859 | 170.2349718 | 107.1127317 | 221.0564591 | 348.759699 | 166.134721 | 1.071919118 | 0.00020916 | 0.00917812 |
| ENSG00000002587 | 302.681011 | 303.371057 | 268.159956 | 125.1124492 | 173.8055646 | 116.9667672 | 291.404008 | 138.62826 | 1.070447162 | 4.40E-05 | 0.00271303 |
| ENSG00000178445 | 178.173929 | 199.330342 | 231.812974 | 72.81134337 | 92.96576711 | 126.6245736 | 203.105748 | 97.467228 | 1.061565082 | 0.00056605 | 0.01909416 |
| ENSG00000132541 | 448.654832 | 723.423289 | 537.935334 | 257.4034815 | 253.6348646 | 313.3421653 | 570.004485 | 274.793504 | 1.05322331 | 7.28E-06 | 0.00068262 |
| ENSG00000184261 | 138.460463 | 125.432264 | 156.695878 | 62.55622459 | 60.62984811 | 80.48172054 | 140.196201 | 67.8892644 | 1.048055488 | 0.00144631 | 0.03855119 |
| ENSG00000279759 | 229.694101 | 325.734949 | 237.466949 | 97.42362845 | 129.343676 | 162.0365307 | 264.298666 | 129.601278 | 1.028679375 | 0.00037156 | 0.01400299 |
| ENSG00000209082 | 271.554241 | 718.561574 | 378.816324 | 273.8116715 | 199.0680013 | 203.8870254 | 456.310713 | 225.588899 | 1.016426974 | 0.00128568 | 0.03542995 |
| ENSG00000166501 | 212.52071 | 192.52394 | 224.543578 | 134.3420561 | 91.95526964 | 85.84716857 | 209.862743 | 104.048165 | 1.012128098 | 0.00057178 | 0.01924613 |
| ENSG00000210140 | 1570.29191 | 3304.9943 | 1899.73559 | 759.9043019 | 1232.806912 | 1369.262339 | 2258.3406 | 1120.65785 | 1.01101256 | 0.00013402 | 0.00656877 |
| ENSG00000223812 | 186.760624 | 265.449675 | 267.352245 | 104.6022116 | 102.0602443 | 152.3787242 | 239.854181 | 119.680393 | 1.005307915 | 0.00059742 | 0.01981293 |
| ENSG00000071575 | 778.169267 | 775.929818 | 826.288058 | 424.5619177 | 383.989038 | 377.7275417 | 793.462381 | 395.426166 | 1.004901399 | 2.84E-08 | 5.14E-06 |
| ENSG00000138061 | 6313.36776 | 4449.44217 | 4922.99679 | 2765.805536 | 2253.409355 | 2801.836964 | 5228.60224 | 2607.01729 | 1.00394551 | 1.47E-08 | 2.92E-06 |
| ENSG00000033122 | 450.801506 | 528.954663 | 554.089548 | 206.1278876 | 242.5193924 | 318.7076133 | 511.281906 | 255.784964 | 1.000424229 | 1.52E-05 | 0.00120165 |

| gene_id | gene_name | gene_chr | gene_start | gene_end | gene_strand | gene_length | gene_biotype | gene_description | tf_family |
|---|---|---|---|---|---|---|---|---|---|
| ENSG00000203837 | PNLIPRP3 | 10 | 116427867 | 116477957 | + | 2354 | protein_coding | pancreatic lipase related protein 3 [Source:HGNC Symbol;Acc:HGNC:23492] | - |
| ENSG00000259070 | LINC00639 | 14 | 38749339 | 38948273 | - | 4860 | lincRNA | long intergenic non-protein coding RNA 639 [Source:HGNC Symbol;Acc:HGNC:27502] | - |
| ENSG00000154545 | MAGED4 | X | 52184823 | 52192268 | + | 4809 | protein_coding | MAGE family member D4 [Source:HGNC Symbol;Acc:HGNC:23793] | - |
| ENSG00000172348 | RCAN2 | 6 | 46220736 | 46491972 | - | 3712 | protein_coding | regulator of calcineurin 2 [Source:HGNC Symbol;Acc:HGNC:3041] | - |
| ENSG00000187950 | OVCH1 | 12 | 29412474 | 29497686 | - | 3958 | protein_coding | ovochymase 1 [Source:HGNC Symbol;Acc:HGNC:23080] | - |
| ENSG00000119686 | FLVCR2 | 14 | 75578617 | 75663214 | + | 4913 | protein_coding | feline leukemia virus subgroup C cellular receptor family member 2 [Source:HGNC Symbol;Acc:HGNC:20105] | - |
| ENSG00000042781 | USH2A | 1 | 215622894 | 216423396 | - | 20506 | protein_coding | usherin [Source:HGNC Symbol;Acc:HGNC:12601] | - |
| ENSG00000223861 | AL365436.1 | 1 | 151557446 | 151557940 | + | 495 | processed_pseudogene | ribosomal protein S10 (RPS10) pseudogene | - |
| ENSG00000196872 | KIAA1211L | 2 | 98793846 | 98936259 | - | 4705 | protein_coding | KIAA1211 like [Source:HGNC Symbol;Acc:HGNC:33454] | - |
| ENSG00000133083 | DCLK1 | 13 | 35768652 | 36131306 | - | 14375 | protein_coding | doublecortin like kinase 1 [Source:HGNC Symbol;Acc:HGNC:2700] | - |
| ENSG00000183036 | PCP4 | 21 | 39867317 | 39929397 | + | 1106 | protein_coding | Purkinje cell protein 4 [Source:HGNC Symbol;Acc:HGNC:8742] | - |
| ENSG00000251003 | ZFPM2-AS1 | 8 | 105546089 | 106060524 | - | 2667 | processed_transcript | ZFPM2 antisense RNA 1 [Source:HGNC Symbol;Acc:HGNC:50698] | - |
| ENSG00000270605 | AL353622.1 | 1 | 28239509 | 28241453 | - | 1945 | antisense | novel transcript | - |
| ENSG00000129151 | BBOX1 | 11 | 27040725 | 27127809 | + | 2464 | protein_coding | gamma-butyrobetaine hydroxylase 1 [Source:HGNC Symbol;Acc:HGNC:964] | - |
| ENSG00000261235 | AC092142.1 | 16 | 82044336 | 82139631 | - | 1030 | antisense | uncharacterized LOC105371363 [Source:NCBI gene;Acc:105371363] | - |
| ENSG00000270127 | AC027020.2 | 15 | 100547765 | 100550153 | - | 2389 | lincRNA | novel transcript | - |
| ENSG00000107562 | CXCL12 | 10 | 44370165 | 44386493 | - | 6077 | protein_coding | C-X-C motif chemokine ligand 12 [Source:HGNC Symbol;Acc:HGNC:10672] | - |
| ENSG00000077498 | TYR | 11 | 89177452 | 89295759 | + | 3016 | protein_coding | tyrosinase [Source:HGNC Symbol;Acc:HGNC:12442] | - |
| ENSG00000165092 | ALDH1A1 | 9 | 72900662 | 73080442 | - | 2849 | protein_coding | aldehyde dehydrogenase 1 family member A1 [Source:HGNC Symbol;Acc:HGNC:402] | - |
| ENSG00000140961 | OSGIN1 | 16 | 83948282 | 83966332 | + | 3460 | protein_coding | oxidative stress induced growth inhibitor 1 [Source:HGNC Symbol;Acc:HGNC:30093] | - |
| ENSG00000166473 | PKD1L2 | 16 | 81100875 | 81220370 | - | 18637 | polymorphic_pseudogene | polycystin 1 like 2 (gene/pseudogene) [Source:HGNC Symbol;Acc:HGNC:21715] | - |
| ENSG00000278035 | AL121895.2 | 20 | 36233851 | 36234297 | + | 447 | lincRNA | novel transcript | - |
| ENSG00000107165 | TYRP1 | 9 | 12685439 | 12710290 | + | 3741 | protein_coding | tyrosinase related protein 1 [Source:HGNC Symbol;Acc:HGNC:12450] | - |
| ENSG00000283031 | AC009242.1 | 2 | 23667208 | 23685453 | - | 2706 | lincRNA | novel transcript | - |
| ENSG00000132386 | SERPINF1 | 17 | 1761959 | 1777574 | + | 3069 | protein_coding | serpin family F member 1 [Source:HGNC Symbol;Acc:HGNC:8824] | - |
| ENSG00000164175 | SLC45A2 | 5 | 33944616 | 33984730 | - | 3267 | protein_coding | solute carrier family 45 member 2 [Source:HGNC Symbol;Acc:HGNC:16472] | - |
| ENSG00000041353 | RAB27B | 18 | 54717860 | 54895516 | + | 7432 | protein_coding | RAB27B, member RAS oncogene family [Source:HGNC Symbol;Acc:HGNC:9767] | - |
| ENSG00000109819 | PPARGC1A | 4 | 23755041 | 23904089 | - | 10870 | protein_coding | PPARG coactivator 1 alpha [Source:HGNC Symbol;Acc:HGNC:9237] | - |
| ENSG00000125965 | GDF5 | 20 | 35433347 | 35454746 | - | 2646 | protein_coding | growth differentiation factor 5 [Source:HGNC Symbol;Acc:HGNC:4220] | - |
| ENSG00000148053 | NTRK2 | 9 | 84668551 | 85027070 | + | 20363 | protein_coding | neurotrophic receptor tyrosine kinase 2 [Source:HGNC Symbol;Acc:HGNC:8032] | - |
| ENSG00000197915 | HRNR | 1 | 152212082 | 152224193 | - | 9623 | protein_coding | hornerin [Source:HGNC Symbol;Acc:HGNC:20846] | - |
| ENSG00000139289 | PHLDA1 | 12 | 76025447 | 76033932 | - | 8069 | protein_coding | pleckstrin homology like domain family A member 1 [Source:HGNC Symbol;Acc:HGNC:8933] | - |
| ENSG00000118322 | ATP10B | 5 | 160563120 | 160852214 | - | 11310 | protein_coding | ATPase phospholipid transporting 10B (putative) [Source:HGNC Symbol;Acc:HGNC:13543] | - |
| ENSG00000158270 | COLEC12 | 18 | 316740 | 500722 | - | 7340 | protein_coding | collectin subfamily member 12 [Source:HGNC Symbol;Acc:HGNC:16016] | - |
| ENSG00000135253 | KCP | 7 | 128862451 | 128910719 | - | 8259 | protein_coding | kielin/chordin-like protein [Source:HGNC Symbol;Acc:HGNC:17585] | - |
| ENSG00000198203 | SULT1C2 | 2 | 108288639 | 108309915 | + | 8399 | protein_coding | sulfotransferase family 1C member 2 [Source:HGNC Symbol;Acc:HGNC:11456] | - |
| ENSG00000182612 | TSPAN10 | 17 | 81637171 | 81648749 | + | 4745 | protein_coding | tetraspanin 10 [Source:HGNC Symbol;Acc:HGNC:29942] | - |
| ENSG00000257681 | AC025265.1 | 12 | 103746315 | 103768858 | - | 1063 | antisense | novel transcript, antisense to STAB2 | - |
| ENSG00000276772 | AC025271.4 | 15 | 57553955 | 57555680 | + | 1726 | lincRNA | novel transcript | - |
| ENSG00000272899 | ATP6V1FNB | 7 | 128866308 | 128872044 | + | 3902 | protein_coding | ATP6V1F neighbor [Source:HGNC Symbol;Acc:HGNC:52392] | - |
| ENSG00000170989 | S1PR1 | 1 | 101236865 | 101243713 | + | 5223 | protein_coding | sphingosine-1-phosphate receptor 1 [Source:HGNC Symbol;Acc:HGNC:3165] | - |
| ENSG00000166963 | MAP1A | 15 | 43510958 | 43531620 | + | 10629 | protein_coding | microtubule associated protein 1A [Source:HGNC Symbol;Acc:HGNC:6835] | - |
| ENSG00000143772 | ITPKB | 1 | 226631690 | 226739323 | - | 7036 | protein_coding | inositol-trisphosphate 3-kinase B [Source:HGNC Symbol;Acc:HGNC:6179] | - |
| ENSG00000164627 | KIF6 | 6 | 39329990 | 39725405 | - | 11868 | protein_coding | kinesin family member 6 [Source:HGNC Symbol;Acc:HGNC:21202] | - |
| ENSG00000180332 | KCTD4 | 13 | 45192853 | 45194717 | - | 1865 | protein_coding | potassium channel tetramerization domain containing 4 [Source:HGNC Symbol;Acc:HGNC:23227] | - |
| ENSG00000285437 | POLR2J3 | 7 | 102562133 | 102572583 | - | 7195 | protein_coding | RNA polymerase II subunit J3 [Source:HGNC Symbol;Acc:HGNC:33853] | - |
| ENSG00000170289 | CNGB3 | 8 | 86553977 | 86743675 | - | 5025 | protein_coding | cyclic nucleotide gated channel beta 3 [Source:HGNC Symbol;Acc:HGNC:2153] | - |
| ENSG00000138316 | ADAMTS14 | 10 | 70672803 | 70762441 | + | 5269 | protein_coding | ADAM metallopeptidase with thrombospondin type 1 motif 14 [Source:HGNC Symbol;Acc:HGNC:14899] | - |
| ENSG00000163762 | TM4SF18 | 3 | 149318498 | 149334414 | - | 4482 | protein_coding | transmembrane 4 L six family member 18 [Source:HGNC Symbol;Acc:HGNC:25181] | - |
| ENSG00000108924 | HLF | 17 | 55265012 | 55325065 | + | 6602 | protein_coding | HLF, PAR bZIP transcription factor [Source:HGNC Symbol;Acc:HGNC:4977] | TF_bZIP |
| ENSG00000138642 | HERC6 | 4 | 88378739 | 88443111 | + | 5793 | protein_coding | HECT and RLD domain containing E3 ubiquitin protein ligase family member 6 [Source:HGNC Symbol;Acc:HGNC:26072] | - |
| ENSG00000235718 | MFRP | 11 | 119338942 | 119346673 | - | 4064 | protein_coding | membrane frizzled-related protein [Source:HGNC Symbol;Acc:HGNC:18121] | - |
| ENSG00000248174 | LINC02268 | 4 | 174094660 | 174220398 | - | 1659 | lincRNA | long intergenic non-protein coding RNA 2268 [Source:HGNC Symbol;Acc:HGNC:53183] | - |
| ENSG00000119922 | IFIT2 | 10 | 89283694 | 89309276 | + | 4101 | protein_coding | interferon induced protein with tetratricopeptide repeats 2 [Source:HGNC Symbol;Acc:HGNC:5409] | - |
| ENSG00000210194 | MT-TE | MT | 14674 | 14742 | - | 69 | Mt_tRNA | mitochondrially encoded tRNA glutamic acid [Source:HGNC Symbol;Acc:HGNC:7479] | - |
| ENSG00000189007 | ADAT2 | 6 | 143422832 | 143450673 | - | 7175 | protein_coding | adenosine deaminase, tRNA specific 2 [Source:HGNC Symbol;Acc:HGNC:21172] | - |
| ENSG00000135069 | PSAT1 | 9 | 78297143 | 78330093 | + | 2188 | protein_coding | phosphoserine aminotransferase 1 [Source:HGNC Symbol;Acc:HGNC:19129] | - |
| ENSG00000132481 | TRIM47 | 17 | 75874161 | 75878575 | - | 3142 | protein_coding | tripartite motif containing 47 [Source:HGNC Symbol;Acc:HGNC:19020] | - |
| ENSG00000155761 | SPAG17 | 1 | 117953861 | 118185223 | - | 9552 | protein_coding | sperm associated antigen 17 [Source:HGNC Symbol;Acc:HGNC:26620] | - |
| ENSG00000253417 | LINC02159 | 5 | 160931778 | 160938626 | - | 2323 | lincRNA | long intergenic non-protein coding RNA 2159 [Source:HGNC Symbol;Acc:HGNC:27749] | - |
| ENSG00000087303 | NID2 | 14 | 52004803 | 52069228 | - | 6392 | protein_coding | nidogen 2 [Source:HGNC Symbol;Acc:HGNC:13389] | - |
| ENSG00000203727 | SAMD5 | 6 | 147508927 | 147737547 | + | 6320 | protein_coding | sterile alpha motif domain containing 5 [Source:HGNC Symbol;Acc:HGNC:21180] | - |
| ENSG00000210174 | MT-TR | MT | 10405 | 10469 | + | 65 | Mt_tRNA | mitochondrially encoded tRNA arginine [Source:HGNC Symbol;Acc:HGNC:7496] | - |
| ENSG00000143387 | CTSK | 1 | 150796208 | 150808323 | - | 1887 | protein_coding | cathepsin K [Source:HGNC Symbol;Acc:HGNC:2536] | - |
| ENSG00000280744 | LINC01173 | 2 | 234682668 | 234717764 | + | 331 | lincRNA | long intergenic non-protein coding RNA 1173 [Source:HGNC Symbol;Acc:HGNC:49545] | - |
| ENSG00000183828 | NUDT14 | 14 | 105172938 | 105181323 | - | 1916 | protein_coding | nudix hydrolase 14 [Source:HGNC Symbol;Acc:HGNC:20141] | - |
| ENSG00000175592 | FOSL1 | 11 | 65892049 | 65900573 | - | 1887 | protein_coding | FOS like 1, AP-1 transcription factor subunit [Source:HGNC Symbol;Acc:HGNC:13718] | TF_bZIP |
| ENSG00000005249 | PRKAR2B | 7 | 107044649 | 107161811 | + | 4318 | protein_coding | protein kinase cAMP-dependent type II regulatory subunit beta [Source:HGNC Symbol;Acc:HGNC:9392] | - |
| ENSG00000157388 | CACNA1D | 3 | 53328963 | 53813733 | + | 20588 | protein_coding | calcium voltage-gated channel subunit alpha1 D [Source:HGNC Symbol;Acc:HGNC:1391] | - |
| ENSG00000002587 | HS3ST1 | 4 | 11393150 | 11429765 | - | 8166 | protein_coding | heparan sulfate-glucosamine 3-sulfotransferase 1 [Source:HGNC Symbol;Acc:HGNC:5194] | - |
| ENSG00000178445 | GLDC | 9 | 6532464 | 6645783 | - | 8939 | protein_coding | glycine decarboxylase [Source:HGNC Symbol;Acc:HGNC:4313] | - |
| ENSG00000132541 | RIDA | 8 | 98102344 | 98117241 | - | 1567 | protein_coding | reactive intermediate imine deaminase A homolog [Source:HGNC Symbol;Acc:HGNC:16897] | - |
| ENSG00000184261 | KCNK12 | 2 | 47516581 | 47570939 | - | 6685 | protein_coding | potassium two pore domain channel subfamily K member 12 [Source:HGNC Symbol;Acc:HGNC:6274] | - |
| ENSG00000279759 | AC118344.2 | 19 | 40355842 | 40360606 | + | 4765 | TEC | TEC | - |
| ENSG00000209082 | MT-TL1 | MT | 3230 | 3304 | + | 75 | Mt_tRNA | mitochondrially encoded tRNA leucine 1 (UUA/G) [Source:HGNC Symbol;Acc:HGNC:7490] | - |
| ENSG00000166501 | PRKCB | 16 | 23835946 | 24220611 | + | 11086 | protein_coding | protein kinase C beta [Source:HGNC Symbol;Acc:HGNC:9395] | - |
| ENSG00000210140 | MT-TC | MT | 5761 | 5826 | - | 66 | Mt_tRNA | mitochondrially encoded tRNA cysteine [Source:HGNC Symbol;Acc:HGNC:7477] | - |
| ENSG00000223812 | AC073365.1 | 3 | 191425526 | 191591097 | + | 2535 | lincRNA | novel transcript | - |
| ENSG00000071575 | TRIB2 | 2 | 12716889 | 12742734 | + | 4588 | protein_coding | tribbles pseudokinase 2 [Source:HGNC Symbol;Acc:HGNC:30809] | - |
| ENSG00000138061 | CYP1B1 | 2 | 38066973 | 38109902 | - | 6014 | protein_coding | cytochrome P450 family 1 subfamily B member 1 [Source:HGNC Symbol;Acc:HGNC:2597] | - |
| ENSG00000033122 | LRRC7 | 1 | 69568398 | 70151945 | + | 14652 | protein_coding | leucine rich repeat containing 7 [Source:HGNC Symbol;Acc:HGNC:18531] | - |

| gene_id | CH_0323_count | CH_0407_count | CH_0603_count | DMSO_0323_count | DMSO_0407_count | DMSO_0603_count | CH_0323_fpkm | CH_0407_fpkm | CH_0603_fpkm | DMSO_0323_fpkm | DMSO_0407_fpkm | DMSO_0603_fpkm |
|---|---|---|---|---|---|---|---|---|---|---|---|---|
| ENSG00000203837 | 32 | 7 | 83 | 3 | 0 | 2 | 0.628141288 | 0.124476935 | 1.226041663 | 0.056264342 | 0 | 0.039249785 |
| ENSG00000259070 | 28 | 22 | 34 | 2 | 3 | 0 | 0.266216876 | 0.189488874 | 0.243263103 | 0.018168211 | 0.026853319 | 0 |
| ENSG00000154545 | 21 | 303 | 87 | 8 | 18 | 10 | 0.201780102 | 2.637455582 | 0.62906869 | 0.073443549 | 0.162828608 | 0.096063624 |
| ENSG00000172348 | 23 | 17 | 29 | 1 | 4 | 2 | 0.286308136 | 0.191707126 | 0.271658704 | 0.011893522 | 0.046877561 | 0.024890623 |
| ENSG00000187950 | 23 | 42 | 20 | 4 | 0 | 5 | 0.26851334 | 0.444192072 | 0.175706489 | 0.044617235 | 0 | 0.058359015 |
| ENSG00000119686 | 55 | 61 | 83 | 0 | 10 | 13 | 0.517284834 | 0.519733097 | 0.587441904 | 0 | 0.088545444 | 0.122239152 |
| ENSG00000042781 | 15 | 36 | 72 | 0 | 6 | 8 | 0.033800578 | 0.07348841 | 0.122091418 | 0 | 0.012728677 | 0.018022821 |
| ENSG00000223861 | 14 | 51 | 25 | 3 | 4 | 4 | 1.306882846 | 4.312829407 | 1.756177483 | 0.267568204 | 0.351534355 | 0.373309063 |
| ENSG00000196872 | 81 | 121 | 179 | 9 | 32 | 10 | 0.795498219 | 1.076522337 | 1.322899944 | 0.084450326 | 0.295871636 | 0.098187028 |
| ENSG00000133083 | 154 | 34 | 234 | 19 | 12 | 25 | 0.495024493 | 0.099007562 | 0.566032801 | 0.058353135 | 0.036315027 | 0.080342603 |
| ENSG00000183036 | 1688 | 2980 | 2563 | 273 | 320 | 401 | 70.52305802 | 112.7868429 | 80.57996491 | 10.89747717 | 12.5865827 | 16.74954396 |
| ENSG00000251003 | 117 | 126 | 176 | 15 | 22 | 21 | 2.027106969 | 1.977629047 | 2.294685899 | 0.248305701 | 0.35884975 | 0.363755878 |
| ENSG00000270605 | 17 | 30 | 39 | 3 | 6 | 3 | 0.403871434 | 0.645652671 | 0.697234063 | 0.068095764 | 0.134197562 | 0.071255008 |
| ENSG00000129151 | 23 | 67 | 33 | 3 | 11 | 5 | 0.431321347 | 1.138233601 | 0.465700636 | 0.053752541 | 0.194207038 | 0.093743905 |
| ENSG00000261235 | 20 | 55 | 30 | 6 | 2 | 8 | 0.897235795 | 2.235232831 | 1.012785849 | 0.257177206 | 0.084470634 | 0.358811624 |
| ENSG00000270127 | 18 | 60 | 28 | 5 | 7 | 5 | 0.348153027 | 1.051313893 | 0.407544913 | 0.09240007 | 0.127466151 | 0.096686891 |
| ENSG00000107562 | 45 | 101 | 127 | 15 | 9 | 24 | 0.342166193 | 0.695711913 | 0.726688152 | 0.108973392 | 0.064426755 | 0.182446588 |
| ENSG00000077498 | 788 | 1194 | 1466 | 249 | 221 | 170 | 12.07281931 | 16.57182325 | 16.90192724 | 3.644907049 | 3.187674137 | 2.603942113 |
| ENSG00000165092 | 40 | 136 | 131 | 17 | 13 | 26 | 0.648755963 | 1.998222119 | 1.598867376 | 0.263435876 | 0.198501542 | 0.421594212 |
| ENSG00000140961 | 121 | 146 | 148 | 6 | 35 | 39 | 1.61593204 | 1.766339007 | 1.487370663 | 0.076558532 | 0.440052941 | 0.520717591 |
| ENSG00000166473 | 1064 | 1497 | 1521 | 252 | 321 | 246 | 2.638028258 | 3.362351387 | 2.837832797 | 0.596956909 | 0.749276324 | 0.609779533 |
| ENSG00000278035 | 39 | 68 | 39 | 4 | 16 | 10 | 4.031539363 | 6.367936037 | 3.033826068 | 0.395067147 | 1.557132043 | 1.033489856 |
| ENSG00000107165 | 2294 | 1837 | 2553 | 748 | 424 | 293 | 28.33475919 | 20.5550601 | 23.72994334 | 8.827390465 | 4.930501903 | 3.618209032 |
| ENSG00000283031 | 43 | 36 | 65 | 8 | 7 | 16 | 0.734267801 | 0.556893324 | 0.835255144 | 0.130521075 | 0.112533864 | 0.273152973 |
| ENSG00000132386 | 357 | 327 | 846 | 109 | 63 | 150 | 5.375082671 | 4.460136673 | 9.585330003 | 1.568007217 | 0.89301066 | 2.25791772 |
| ENSG00000164175 | 371 | 424 | 567 | 110 | 100 | 96 | 5.247332637 | 5.432678754 | 6.034864441 | 1.486490022 | 1.331569527 | 1.357487502 |
| ENSG00000041353 | 734 | 399 | 1476 | 152 | 150 | 278 | 4.563564357 | 2.247317831 | 6.90580405 | 0.902934682 | 0.878008136 | 1.7280362 |
| ENSG00000109819 | 248 | 290 | 561 | 60 | 83 | 106 | 1.054231424 | 1.116775189 | 1.794596895 | 0.243691372 | 0.33217086 | 0.450495091 |
| ENSG00000125965 | 178 | 170 | 226 | 23 | 78 | 40 | 3.108450694 | 2.689406093 | 2.9699709 | 0.383757118 | 1.28238298 | 0.698367295 |
| ENSG00000148053 | 96 | 106 | 202 | 17 | 44 | 38 | 0.217842841 | 0.217902096 | 0.344939717 | 0.036857477 | 0.093999144 | 0.086209589 |
| ENSG00000197915 | 76 | 97 | 371 | 34 | 33 | 61 | 0.364936185 | 0.42194763 | 1.34059322 | 0.155986452 | 0.149182004 | 0.292841815 |
| ENSG00000139289 | 1112 | 1402 | 2341 | 459 | 323 | 442 | 6.367938968 | 7.273201189 | 10.08823738 | 2.511374139 | 1.74138897 | 2.530557997 |
| ENSG00000118322 | 771 | 609 | 845 | 221 | 209 | 171 | 3.149964024 | 2.253990029 | 2.597931518 | 0.862676795 | 0.803890069 | 0.698469179 |
| ENSG00000158270 | 190 | 180 | 205 | 75 | 46 | 36 | 1.196110662 | 1.026534969 | 0.971161363 | 0.451111243 | 0.272630697 | 0.226579275 |
| ENSG00000135253 | 1586 | 2729 | 2828 | 466 | 845 | 649 | 8.873389338 | 13.8316303 | 11.90653886 | 2.491018186 | 4.450842488 | 3.630203506 |
| ENSG00000198203 | 654 | 948 | 1178 | 267 | 250 | 244 | 3.598023435 | 4.724741572 | 4.876983698 | 1.403466749 | 1.294867736 | 1.342072527 |
| ENSG00000182612 | 6294 | 10077 | 8714 | 2233 | 2843 | 2473 | 61.29207755 | 88.89807251 | 63.85794428 | 20.77643138 | 26.06475369 | 24.07695942 |
| ENSG00000257681 | 77 | 83 | 79 | 13 | 38 | 22 | 3.347119988 | 3.268452146 | 2.584207732 | 0.539918906 | 1.555117879 | 0.956099646 |
| ENSG00000276772 | 90 | 80 | 89 | 18 | 40 | 23 | 2.409436797 | 1.940199992 | 1.793010406 | 0.460415739 | 1.008166314 | 0.615603083 |
| ENSG00000272899 | 543 | 854 | 1019 | 266 | 242 | 233 | 6.430228191 | 9.1615279 | 9.08072479 | 3.009628004 | 2.697994645 | 2.758559763 |
| ENSG00000170989 | 298 | 271 | 573 | 130 | 74 | 156 | 2.636392447 | 2.17193372 | 3.814768526 | 1.098858506 | 0.616346134 | 1.379806904 |
| ENSG00000166963 | 191 | 78 | 130 | 39 | 44 | 50 | 0.830337746 | 0.307184642 | 0.425289382 | 0.161990911 | 0.180083222 | 0.217315818 |
| ENSG00000143772 | 1146 | 807 | 1571 | 379 | 358 | 405 | 7.526145454 | 4.801147039 | 7.763971794 | 2.378109385 | 2.213452355 | 2.659150597 |
| ENSG00000164627 | 231 | 263 | 311 | 111 | 82 | 84 | 0.899390431 | 0.927631556 | 0.911205738 | 0.41291807 | 0.300572537 | 0.326975709 |
| ENSG00000180332 | 76 | 111 | 112 | 40 | 39 | 26 | 1.882992442 | 2.491388443 | 2.088203317 | 0.946890159 | 0.90970117 | 0.644033196 |
| ENSG00000285437 | 91 | 110 | 95 | 27 | 27 | 51 | 0.584419118 | 0.639969372 | 0.459120201 | 0.165672877 | 0.163247278 | 0.327456126 |
| ENSG00000170289 | 161 | 99 | 155 | 68 | 43 | 39 | 1.480483701 | 0.82470083 | 1.072578845 | 0.59743587 | 0.372259142 | 0.358543854 |
| ENSG00000138316 | 449 | 619 | 667 | 180 | 189 | 258 | 3.937603324 | 4.917674204 | 4.401809365 | 1.508213258 | 1.560438252 | 2.262065878 |
| ENSG00000163762 | 240 | 264 | 545 | 177 | 126 | 74 | 2.474304871 | 2.465638357 | 4.22822651 | 1.74349161 | 1.222958374 | 0.762734883 |
| ENSG00000108924 | 114 | 253 | 301 | 47 | 95 | 98 | 0.797890239 | 1.604140132 | 1.585347859 | 0.314297398 | 0.625980879 | 0.685747601 |
| ENSG00000138642 | 138 | 84 | 176 | 41 | 42 | 63 | 1.100751735 | 0.606978154 | 1.056434886 | 0.312463128 | 0.315397861 | 0.502401309 |
| ENSG00000235718 | 1777 | 1919 | 2819 | 632 | 689 | 1059 | 20.20447403 | 19.76599032 | 24.11987048 | 6.865652634 | 7.375279867 | 12.0380461 |
| ENSG00000248174 | 162 | 181 | 235 | 65 | 65 | 87 | 4.512138783 | 4.566984017 | 4.925553844 | 1.72975828 | 1.704433074 | 2.422627305 |
| ENSG00000119922 | 168 | 95 | 169 | 64 | 58 | 46 | 1.892924678 | 0.96968603 | 1.432948328 | 0.688983232 | 0.615249411 | 0.51818138 |
| ENSG00000210194 | 143 | 411 | 220 | 87 | 100 | 113 | 95.7636669 | 249.338897 | 110.868248 | 55.66581982 | 63.04692241 | 75.65595087 |
| ENSG00000189007 | 166 | 301 | 342 | 113 | 94 | 113 | 1.069054887 | 1.756070281 | 1.657439922 | 0.695304413 | 0.569926604 | 0.727562454 |
| ENSG00000135069 | 257 | 462 | 594 | 192 | 180 | 146 | 5.427497427 | 8.838772602 | 9.440015819 | 3.874113666 | 3.578806107 | 3.082614943 |
| ENSG00000132481 | 348 | 492 | 662 | 172 | 161 | 259 | 5.117842115 | 6.5547514 | 7.326311895 | 2.416800009 | 2.229116044 | 3.808091059 |
| ENSG00000155761 | 320 | 496 | 553 | 229 | 188 | 137 | 1.547994756 | 2.173625226 | 2.013095659 | 1.058423847 | 0.856202552 | 0.662582551 |
| ENSG00000253417 | 2097 | 2057 | 3056 | 1042 | 883 | 996 | 41.71219473 | 37.06656871 | 45.74437885 | 19.80327219 | 16.53577203 | 19.80723571 |
| ENSG00000087303 | 997 | 449 | 1070 | 483 | 357 | 198 | 7.207293576 | 2.940403138 | 5.820772239 | 3.336021279 | 2.429654004 | 1.431008341 |
| ENSG00000203727 | 426 | 556 | 473 | 169 | 173 | 256 | 3.114629132 | 3.682603963 | 2.602421614 | 1.180560026 | 1.190808723 | 1.871270746 |
| ENSG00000210174 | 147 | 200 | 154 | 55 | 76 | 77 | 104.5003629 | 128.7994303 | 82.38363663 | 37.3566377 | 50.86431709 | 54.72567283 |
| ENSG00000143387 | 1836 | 1667 | 2214 | 920 | 776 | 677 | 44.95878824 | 36.97949726 | 40.7980411 | 21.52456458 | 17.88968952 | 16.57412118 |
| ENSG00000280744 | 165 | 164 | 273 | 85 | 57 | 110 | 23.03402167 | 20.74021037 | 28.67928026 | 11.33729323 | 7.491345795 | 15.35247619 |
| ENSG00000183828 | 428 | 481 | 327 | 166 | 238 | 141 | 10.32195794 | 10.50864871 | 5.934523346 | 3.824996402 | 5.403739873 | 3.399674589 |
| ENSG00000175592 | 313 | 550 | 816 | 153 | 257 | 289 | 7.664542875 | 12.20079394 | 15.03667639 | 3.579628674 | 5.924806969 | 7.075215687 |
| ENSG00000005249 | 203 | 214 | 390 | 103 | 113 | 124 | 2.172337106 | 2.074571648 | 3.140621242 | 1.053108297 | 1.138436438 | 1.326639086 |
| ENSG00000157388 | 288 | 327 | 519 | 166 | 106 | 206 | 0.646386308 | 0.664861057 | 0.876570383 | 0.355969162 | 0.223977652 | 0.462239231 |
| ENSG00000002587 | 282 | 312 | 332 | 122 | 172 | 109 | 1.595708481 | 1.599346342 | 1.413716422 | 0.659582163 | 0.916288115 | 0.616638822 |
| ENSG00000178445 | 166 | 205 | 287 | 71 | 92 | 118 | 0.858090258 | 0.959980092 | 1.116417291 | 0.350661317 | 0.447725544 | 0.609827228 |
| ENSG00000132541 | 418 | 744 | 666 | 251 | 251 | 292 | 12.32596999 | 19.87473021 | 14.77878828 | 7.071689321 | 6.968153473 | 8.608502228 |
| ENSG00000184261 | 129 | 129 | 194 | 61 | 60 | 75 | 0.89166582 | 0.80776606 | 1.009099319 | 0.402853249 | 0.390447657 | 0.518290911 |
| ENSG00000279759 | 214 | 335 | 294 | 95 | 128 | 151 | 2.075222603 | 2.942925072 | 2.145448135 | 0.880195508 | 1.168584299 | 1.46395519 |
| ENSG00000209082 | 253 | 739 | 469 | 267 | 197 | 190 | 155.873784 | 412.4587088 | 217.4428712 | 157.169563 | 114.2662422 | 117.0323913 |
| ENSG00000166501 | 198 | 198 | 278 | 131 | 91 | 80 | 0.825285351 | 0.747631548 | 0.871973961 | 0.521692831 | 0.35709149 | 0.333371795 |
| ENSG00000210140 | 1463 | 3399 | 2352 | 741 | 1220 | 1276 | 1024.26943 | 2155.780464 | 1239.158832 | 495.6700978 | 804.1348376 | 893.1419333 |
| ENSG00000223812 | 174 | 273 | 331 | 102 | 101 | 142 | 3.171648901 | 4.507980059 | 4.54029033 | 1.776399555 | 1.733230778 | 2.587760753 |
| ENSG00000071575 | 725 | 798 | 1023 | 414 | 380 | 352 | 7.301774523 | 7.280761167 | 7.753286265 | 3.983780299 | 3.6030739 | 3.544320572 |
| ENSG00000138061 | 5882 | 4576 | 6095 | 2697 | 2230 | 2611 | 45.19344178 | 31.85076699 | 35.24064762 | 19.79866787 | 16.13074485 | 20.05659428 |
| ENSG00000033122 | 420 | 544 | 686 | 201 | 240 | 297 | 1.324543425 | 1.554172759 | 1.628023991 | 0.605644245 | 0.712569639 | 0.936425606 |
